# CEP-IP with CMRF in SEA-AD Microglia: A Mitochondrial Co-expression Module Linked to Cognition

**DOI:** 10.64898/2026.10.22.761284

**Authors:** Kah Keng Wong

## Abstract

Single-nucleus RNA-seq (snRNA-seq) studies of Alzheimer’s disease (AD) routinely filter nuclei by mitochondrial percentage (%MT), yet whether these reads carry information about cognition is rarely examined. In the Seattle Alzheimer’s Disease Brain Cell Atlas (SEA-AD) microglia snRNA-seq dataset, a transcriptome-wide screen identified a module of *MT-CO3* and eight co-expressed mitochondrial genes. Three microglial populations were analyzed: PVM_3, M4 (Mic.13-like), and PVM_1, with a housekeeping-gene control. The CEP-IP pipeline with a copula monotone recovery filter (CMRF) used per-cell Spearman and Kendall scores, an inflection point (IP), and a cross-validated copula space boundary to isolate *MT-CO3* Module cells. The donor percentage of Module cells past the IP (Module post-IP%) was significantly associated with worse cognition (MMSE and CASI; FDR<0.05) in all three populations. In PVM_3 and M4, these associations remained significant after adjustment for tissue quality, plaque burden, tangle burden, and contamination, and after doublet removal. PVM_3 and M4 Module post-IP% repeated in each donor’s regions beyond chance, and no per-cell %MT cutoff reproduced the Module cells. In two external datasets without cognitive scores, the *MT-CO3* module and its IP replicated. However, at the donor level, Module post-IP% could not be separated from mean %MT, and the source of the mitochondrial transcripts (e.g., retained cytoplasm or a true change in mitochondrial content) remains unresolved. Routine %MT filtering in AD snRNA-seq may therefore remove cells that carry information about cognition. The R package cepip, the pipeline code, and a software container that reproduced the findings are publicly available.

## 1. Introduction

Monotonicity is one of the simpler and more explainable relationships for downstream characterization. Non-parametric statistics such as Spearman r_s_ and Kendall’s *τ*_b_ are widely used to identify monotonic relationships at the population level [1], and have been applied to measure gene-gene co-expression in single-cell transcriptomics [2]. However, a population-level statistic summarizes the relationship for all cells in a single value. Decomposition of these population-level metrics into contributions of single cells uncovers the strength of the monotonic relationship held by each individual cell. Such cell subpopulations may demonstrate clinically relevant properties.

Flow cytometry gating, through the expression of biomarkers, enables the identification of immune cell subpopulations with clinical implications [3]. The gates are drawn on two-dimensional plots of paired markers and set within each experiment with controls [4, 5]. In contrast, single-cell transcriptomic studies often combine cells from many donors and experiments in one analysis. Expression levels then differ between processing batches, and donors are often processed in separate batches [6–8]. Consequently, a fixed expression cutoff applied to pooled samples can select cells from some donors or batches more than others. One approach to mitigate this is to gate pooled samples by how two genes vary together in ranks, such as a monotonic relationship measured by rank correlation, which is less affected by shifts in the absolute expression level of either gene [9–11].

The relationship between two genes is described by their joint distribution, which combines the marginal distribution of each gene (the raw counts) with the dependency between them [12, 13]. The marginal distributions can be distorted by outliers or technical factors such as sequencing depth and postmortem interval (PMI) [14–16]. Transforming the depth-normalized expression values into empirical copula space separates the dependency structure from the marginal distributions [12, 17]. In copula space, both axes are ranks from 0 to 1 that enable interpretation of the joint dependency in the whole expression range (outliers are capped) and log-transformation or scaling of either gene does not change the dependency [12, 17].

In this study, the published cell explanatory power with inflection point (CEP-IP) framework [18] was applied to microglial populations in the Seattle Alzheimer’s Disease Brain Cell Atlas (SEA-AD) immune single-nucleus RNA-seq (snRNA-seq) dataset [19, 20]. Firstly, a transcriptome-wide screen identified the dual-filtered genes (DFGs) of each gene of interest (GOI), i.e., partner genes with both r_s_≥0.6 and τ_b_≥0.5. *MT-CO3* was the top-ranked mitochondrial GOI, and its eight DFGs were averaged to form the composite DFG (cDFG). Secondly, r_s_ and *τ*_b_ of *MT-CO3* with cDFG were decomposed into the cell-level scores S_i_ and K_i_. Cells that passed cutoffs on both scores formed the Spearman-Kendall cell-level score (SKCS) core. Thirdly, the inflection point (IP) was located on a generalized additive model (GAM) fit of cDFG on *MT-CO3* [21–23]. The fitted cDFG value at the IP (IP_y_) split the cells into pre-IP and post-IP groups. Next, the copula monotone recovery filter (CMRF) recovered cells in copula space that followed the same monotonic relationship as the SKCS-core and formed the Module, validated in the copula pitcher plot and a raw-count UMAP. The donor percentage of Module cells at or above IP_y_ (i.e., Module post-IP%) was tested for association with clinical outcomes. Five Batteries of analyses then tested whether these associations could arise from contamination, measurement artifacts, per-cell mitochondrial percentage (%MT), or the 11 severely affected (SA) donors denoted by SEA-AD.

## 2. Materials and Methods

### 2.1 Dataset and definitions

All results were computed on the SEA-AD immune snRNA-seq dataset, Multiregion release of June 2026 (https://brain-map.org/consortia/sea-ad/our-data) [19, 20]. Battery 1 also used the Gerrits et al. [24] dataset (GSE148822) for technical checks. The pipeline was built upon the published CEP-IP framework [18]. The GOI/DFG screen was extended from a targeted panel, as originally used by the framework, to the whole transcriptome. SKCS-core, copula space, pitcher plots, and CMRF were developed for this study. Development started in April 2026 on the earlier snRNA-seq dataset pre-release of July 2025. After the June 2026 release, every step was rerun from the start on the new release and refined over several rounds of reruns, with the final run order recorded in the deposited logs.

The term “region set” denotes the neocortical regions (NeoC; DFC, MTG, AnG, STG, V1C, and ITG) or the non-neocortical regions (NonNeoC; HIP, MEC, LEC, and FI). Following Mesulam (1998) [25], NeoC comprises primary and association isocortex, while NonNeoC comprises limbic (HIP) and paralimbic (MEC, LEC, and FI) cortex. “Arm” denotes one of four groups of cells analyzed in parallel: M4 (Mic.13-like cells with *MT-CO3*>0 shortlisted from Micro-PVM_3 cells), PVM_3 (all Micro-PVM_3 cells with *MT-CO3*>0), PVM_1 (all Micro-PVM_1 cells with *MT-CO3*>0), and HKG8 (the control group consisting of the M4 cells with *ACTB*>0, analyzed with *ACTB* as the GOI and eight other housekeeping genes [HKG8] as its partners). M4, PVM_3, and PVM_1 are the main arms.

“Unit” denotes one donor in one of the brain regions of NeoC or NonNeoC (e.g., MTG or LEC) in one arm, with at least 10 cells of the arm. “Donor percentage” denotes the cells of one cell population in a donor, as a percentage of that donor’s cells of the arm’s supertype. Module post-IP% is the donor percentage of Module post-IP cells. Multiple testing was corrected by the Benjamini-Hochberg procedure, yielding the false discovery rate (FDR); 95% confidence intervals (CIs), from donor bootstraps unless stated otherwise, are provided in the Supplementary Tables.

### 2.2 Selection of cell groups and GOI-DFG screen

M4 cells were selected as the cells ranked in the top 3.0% on each of two Mic.13 marker scores [26], which weight each marker gene by its published fold change alone or by both its fold change and its adjusted *p*-value (Supplementary Methods S1). Target and control supertypes were chosen if they were myeloid, had at least 25 donors with ≥5 cells in each region set, and had the module emerge in a transcriptome-wide screen with module r_s_≥0.6. In this screen, each gene was tested as a GOI, and a gene became a DFG partner of a GOI when the pair reached Spearman r_s_≥0.6 and Kendall’s *τ*_b_≥0.5 (Supplementary Methods S2). Gene-level and technical controls, including ambient RNA and DecontX [27], are in Supplementary Methods S3.

### 2.3 Cell-level scores and the SKCS-core

For each arm and region set, the composite cDFG (only GOI^+^ cells of each arm were included) was fitted on the GOI by GAMs [21–23] with a donor random intercept (to remove donor-to-donor differences in the mean composite) and inverse-donor weights (so that each donor carries the same total weight regardless of the number of cells). Deviance explained (DE) was measured against a donor-only model.

Spearman r_s_ is the Pearson correlation of ranks, and Kendall’s *τ*_b_ measures the net agreement between all pairs of observations. For n cells with ranks R_x,i_ on the GOI and R_y,i_ on the composite cDFG or cHKG8 (ties averaged; middle rank *R̄* = (*n* + 1)⁄2):

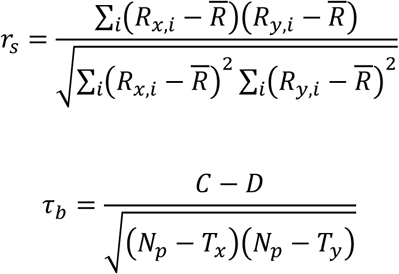

where C and D are the numbers of concordant and discordant pairs, respectively, *N_p_* = *n*(*n* − 1)⁄2 is the number of pairs, and T_x_ and T_y_ are the pairs tied on the GOI and on the composite. Both statistics add up contributions of individual cells, thus each cell *i* can be assigned its own score [1, 28, 29]:

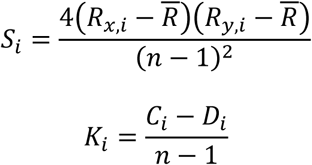

where C_i_ and D_i_ are the concordant and discordant pairs, respectively, involving cell *i*. Without ties, r_s_ is close to three times the mean S_i_ and *τ*_b_ is the mean K_i_ (derivation and ties descriptions are provided in Supplementary Methods S4).

S_i_ grows with the distance of a cell from the middle of both rank orders in the direction of the relationship, and is near zero for a cell near the middle GOI rank regardless of its composite value (Supplementary Methods S4). K_i_ measures how consistently a cell is ordered against all other cells. Cells with S_i_>0.05 and K_i_>0.5 (without ties; more than three quarters of their pairs concordant) formed the SKCS-core. K_i_ plotted against S_i_ is the pitcher plot.

Within each fitted population, cells were ranked by explanatory power (EP; the reduction in squared error of the population-level fit over the mean), and split into top-ranked explanatory power (TREP) or non-TREP cells according to the fitted DE (Supplementary Methods S4) [18].

### 2.4 Inflection point (IP), pre-IP and post-IP groups, and copula coordinates

The IP was located on all cells with GOI>0 with the IP detection procedure of the CEP-IP framework and graded by its IP reliability score (IPRS, 0 to 1; Supplementary Methods S5). IP_y_ was the IP on the composite cDFG or cHKG8 axis fitted at IP_x_, i.e., IP_y_ = f(IP_x_). SKCS-core cells below IP_y_ were defined as pre-IP and the others as post-IP, and each pre-IP and post-IP group was split into TREP and non-TREP cells. In copula space, each cell’s rank was divided by n + 1, and both axes range from 0 to 1. This was applied to the GOI and the composite (u, v), and to S_i_ and K_i_ (u_SK_, v_SK_) (Supplementary Methods S6).

### 2.5 Copula monotone recovery filter (CMRF)

The SKCS-core cells followed the *MT-CO3*/cDFG monotone relationship, but the fixed Si>0.05 and K_i_>0.5 cutoffs also excluded cells that may belong to it. CMRF recovered these cells in three stages (Supplementary Methods S7). Stage 1 admitted cells within a tolerance band (δ_base_ and δ_adapt_) around a spline g(u) = E[v | u] that was fitted to the TREP cells of the SKCS-core as a penalized spline with a donor random intercept. The band extended δ_base_ above and below the spline, where δ_base_ was the donor-weighted 95th percentile of |v - g(u)| over the SKCS-core. δ_base_ was widened around IP_u_ as δ_adapt_(u) = δ_base_[1 + κ·K(u)], where K is a Gaussian kernel centered at IP_u_ with bandwidth 0.10 and κ=0.50. SKCS-core cells within the band formed the reference set (Original + Refined, OriRef), SKCS-core cells outside it were labeled Original-only, and non-SKCS cells within it were candidates.

Stage 2 tested whether the candidates were as monotone-coherent as OriRef by repeated stratified k-fold cross-validation (RSk-fCV): five folds repeated 40 times yielded 200 root mean squared errors (RMSE) of isotonic (PAVA) fits for Curve A (OriRef only) and Curve B (OriRef plus candidates). In each RSk-fCV iteration, when a one-sided paired Wilcoxon signed-rank test yielded *p*<0.05 and ΔRMSE=median(RMSE_B_ - RMSE_A_) exceeded the background variability BV = IQR(RMSE_A_) (interquartile range, IQR), the candidate with the largest |v - PAVA(u)| was removed and Curve B was refitted; the gate closed when either criterion failed. OriRef plus Recovered (the candidates that passed Stage 2) formed the Module. For validation of the Recovered cells, after the GAM was refitted on the final Module and TREP recomputed, Recovered cells were compared with the pre-IP and post-IP TREP cells in the copula pitcher plot (u_SK_, v_SK_) and in a UMAP built with Monocle3 from the raw counts, which uses no CMRF input (Supplementary Methods S7). Differential expression between Module and non-Module cells, by paired pseudobulk per donor, is described in Supplementary Methods S8.

### 2.6 Clinical association, benchmark against competitors, and boundary ablation

For each donor, arm, and region set, the donor percentage of four cell populations was computed: Module pre-IP, Module post-IP, non-Module pre-IP, and non-Module post-IP (pre-IP and post-IP were split at IP_y_). Module post-IP% on each arm was computed as:

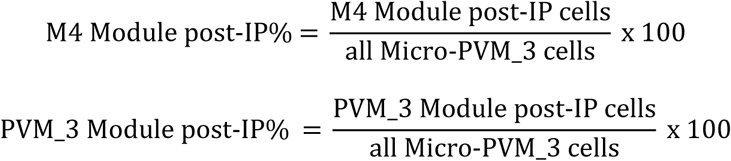

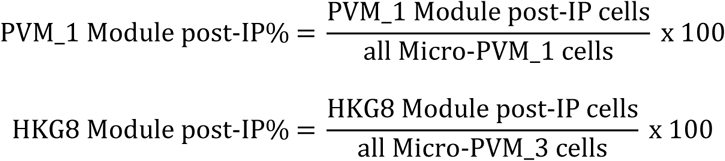

where Module post-IP cells are the arm’s Module cells with a composite at or above IP_y_. The denominator is all of the donor’s cells of the supertype in the region set.

Association of each donor percentage with nine outcomes including cognitive status, Mini-Mental State Examination (MMSE), Cognitive Abilities Screening Instrument (CASI), CERAD score, Braak stage, Thal phase, AD neuropathologic change (ADNC), cerebral amyloid angiopathy (CAA), and *APOE* ε4 dose, was examined by ordinary least squares (OLS) with standardized outcome and predictor, adjusted for age at death, sex, PMI, and *APOE* ε4 dose (termed “cov4” due to four covariates), and effect sizes were the standardized β and ΔR^2^ (R^2^ gained over cov4). Module post-IP% was then tested on cognitive outcomes adjusted for pathology and on pathology outcomes adjusted for cognition, and Module and non-Module post-IP% were fitted in one model (Supplementary Methods S9).

For benchmark against competitors, five donor-level competitors without CMRF input, i.e., cluster proportion (CP), AUCell, UCell, AddModuleScore, and changepoint split (CS), were each fitted alone with cov4 and then with Module post-IP% in one model, called Module only, competitor only, Both, or Neither from the FDR of each coefficient, or unreadable when the variance inflation factor (VIF) was ≥5. A nested F-test examined whether Module post-IP% explained additional variance in MMSE or CASI beyond a model that already held CP and AUCell, adjusted for cov4 alone or for cov4 with Braak, Thal, and CERAD (+BTC) (Supplementary Methods S9).

In terms of CMRF boundary ablation, three simpler boundaries (OLS, fixed spline, and local spline), each keeping about as many cells (within 19) as the CMRF Module, were compared with CMRF on two tests: how far their kept cells overlapped those of the CMRF Module (Jaccard index), and whether their kept and discarded cells showed the same associations with the clinical and negative-control outcomes at cov4, and with the cognitive outcomes after adding +BTC (Supplementary Methods S9).

To separate how many M4 cells a donor has from the state of those cells, the M4 formula for Module post-IP% was written as two factors, using the fact that the M4 cells are a subset of the donor’s Micro-PVM_3 cells:

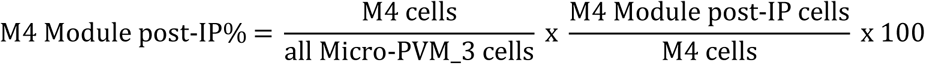

where the first and second factors, each multiplied by 100, are the M4 share and within-M4 fraction, respectively (donors with ≥5 M4 cells):

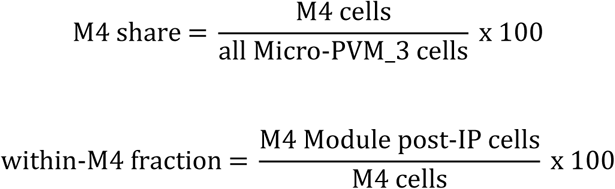

Integrating the M4 share in the same model held the number of M4 cells constant, thus any remaining association of Module post-IP% with an outcome originated from the within-M4 fraction.

### 2.7 Batteries 1-5

As CMRF’s Module post-IP% separated cognition from pathology outcomes, five thematic “Batteries” were constructed to stress-test these observations against contamination, measurement artifacts, the percentage of unique molecular identifiers (UMIs) from mitochondrial genes (%MT), and the 11 donors that SEA-AD denoted as SA. SEA-AD excluded these donors from its analyses of gene expression changes because of systematically lower data quality [19]. Here, the SA donors were retained in the main analyses because all 11 had High ADNC, and excluding them would narrow the range of pathology. Tissue quality was instead adjusted for in the cov7 model, and Battery 5 tested whether the SA donors carried the results.

This section describes the aim of each Battery and the terms used in the Results; the full procedures are in Supplementary Methods S10-S14. The five Batteries shared one set of settings (Supplementary Methods A2), and readings were fixed before each run in pre-specified reading files (Batteries_1-5_Pre-specified_Readings.md in this project’s GitHub repository). Two readings were changed after the first run (the offset share and the donor requirement; Supplementary Methods A2). The readings named M4 as the primary arm, as planned when the analyses were designed, while PVM_3 was recommended after the clinical analyses (see Discussion).

Battery 1 tested whether Module post-IP% and the *MT-CO3*/DFG co-expression that defines the Module could be due to ambient RNA contamination or doublets:

1. *Clinical models adjusted for contamination or doublets.* The clinical models of Section 2.6 were refitted with donor mean DecontX contamination (the estimated fraction of ambient RNA per nucleus) as a covariate, or after removing Module post-IP nuclei with a high SEA-AD doublet score.
2. *Cells within donor.* DecontX contamination was compared between Module post-IP, Module pre-IP, and non-Module cells within each donor, and the r_s_ of *MT-CO3* with cDFG between the least and the most contaminated tertile of Module cells. If ambient RNA produced the co-expression, the r_s_ would rise with contamination.
3. *Ambient RNA in an external dataset.* The Gerrits et al. dataset (GSE148822) [24], which also contains the empty droplets, was used to compare the mitochondrial fraction of ambient RNA with that of the nucleus-containing droplets. r_s_ of *MT-CO3* with each DFG was computed before and after removing the ambient index (a nucleus’s UMIs from other lineages’ marker genes). SEA-AD nuclei were also down-sampled to the Gerrits sequencing depth and mitochondrial fraction.

Battery 2 examined whether Module post-IP% or the MT-CO3 module arose from artifacts of tissue sampling, CMRF cell assignment, or library preparation and sequencing:

1. *Consistency between regions.* The donor share of variance of Module post-IP%, with or without tissue quality removed (TQR; PMI, RIN, brain pH, and fresh brain weight), was compared with a null that randomly shuffled donor labels within each region, and split-half r_s_ measured whether Module post-IP% was reproducible over random halves of a donor’s regions.
2. *CMRF label agreement.* Cohen’s κ compared the Module labels of M4 and HKG8 with those of PVM_3 on the cells they share. M4, whose CMRF run used the same mitochondrial genes as PVM_3, was expected to agree well (pre-specified κ>0.60), and HKG8, whose run used housekeeping genes (*ACTB* with cHKG8), was expected to agree no better than chance (pre-specified |κ|<0.20).
3. *Within-donor co-expression.* A donor with more mitochondrial RNA would have high *MT-CO3* and cDFG levels, so pooling cells over donors could produce a correlation that is absent within donors. The r_s_ of *MT-CO3* with cDFG was therefore computed within each donor and compared with that of a control pair (*ACTB* with cHKG8).
4. *Offset share.* The share of mitochondrial UMIs that every nucleus carries regardless of sequencing depth (a fixed background, as expected of ambient RNA in each droplet) was compared between Module post-IP and pre-IP cells against a pre-specified margin of ±10 percentage points (pp).
5. *Library fields.* Five library fields recorded by SEA-AD, including the intronic mapping rate (a lower rate indicates relatively more cytoplasmic RNA), were tested against Module post-IP% between donors and within donor, alongside %MT, sequencing depth, and the HKG8 control arm.

Battery 3 assessed whether Module post-IP% tracked microglial activation, and whether it carried information beyond %MT on histology and clinical outcomes. Each unit was matched by donor and region to the SEA-AD quantitative neuropathology (28 stain measures, six of them on IBA1⁺ microglia) and to the continuous pseudo-progression scores (CPS; three local scores per region and three global scores per donor). The following was tested:

1. *Microglial activation.* Within donor, the r_s_ of Module post-IP%, %MT, and the cytoplasmic share (own-ER plus ribosomal genes as a percentage of non-mitochondrial UMIs) with the six IBA1 measures were compared with directions set in advance: positive for activated IBA1⁺ cells per area, and negative for inactivated IBA1⁺ cells per area and for process area and length per cell.
2. *Pathology progression.* The r_s_ with the CPS were computed within donor for the three local scores and between donors for all six CPS.
3. *Independence from %MT.* Module post-IP% and %MT were entered together in one model per comparison. The verdict was “Module only” when the Module post-IP% coefficient alone had FDR<0.05, “%MT only” when the %MT coefficient alone did, “Both” when both did, and “Neither” when neither did. It was “unreadable” when the two predictors were too collinear to separate (VIF≥5).

Battery 4 examined whether a simpler quantity could replace Module post-IP%:

1. *Nuclear-encoded panels.* Each of 15 gene panels encoded in the nucleus rather than in mitochondrial DNA (14 for mitochondrial functions such as OXPHOS, the mitoribosome, and the TCA cycle, and one for glycolysis) and a matched reference set was entered with Module post-IP% or with %MT in the clinical model, with verdicts as in Battery 3.
2. *%MT as a cell-level threshold.* The area under the curve (AUC) of per-cell %MT for Module membership measured whether a %MT threshold could reproduce the Module cells, with cells pooled over donors and within each donor.
3. *Donor requirement to reproduce the clinical association observations.* Donors were subsampled from 10 to 83 to find the smallest number at which the association of Module post-IP% with MMSE reached *p*<0.05 in at least 80% of draws, also with the donor effect reduced to 0.75x and 0.5x the observed value. To test whether sampling more regions per donor changed the results, the donor share of variance and split-half r_s_ were recomputed for donors with at least three regions, then at least four regions, and so on.

Battery 5 tested whether the 11 donors that SEA-AD flagged as SA, all with High ADNC, carried the between-donor and clinical results of Batteries 1 to 4. The nine analyses exposed to these donors were recomputed without them. As all SA donors had High ADNC, each change was compared with the changes from 1,000 random removals of the same number of non-SA donors with High ADNC. A comparison that was significant with all donors held if it stayed significant with the same sign. Otherwise, the change toward zero was ranked among the 1,000 random removals. A change larger than in 95% of them was called lost beyond random removal, which means the SA donors carried the result. A smaller change was called lost within random removal, which was interpreted as a loss of power.

### 2.8 M4 against PVM_3 and summary of all tests

A donor can have a high M4 Module post-IP% for two reasons. It may have many M4 cells among its Micro-PVM_3 cells (a high M4 share), or many of its M4 cells may be Module post-IP cells (a high within-M4 fraction). Module post-IP% in M4 depends on both the M4 share and the within-M4 fraction (Section 2.6). The analyses in this section tested which of the two carried the M4 associations. The M4 share alone could produce an association of M4 Module post-IP% with an outcome.

To test this, the M4 share was added as a second predictor to the M4 models, and the change in the Module post-IP% coefficient was estimated by donor bootstrap. The within-M4 fraction was also fitted with %MT in one model. To test whether the M4 results were a property of the M4 cells, each donor’s M4 cells were also compared with as many PVM_3 arm cells drawn at random within the same donor and region (10,000 draws per region set). This kept the M4 share identical in every draw, and all cells were scored on the PVM_3 labels. These analyses were added after Battery 5 was complete, and their readings were fixed before each was first run (Supplementary Methods S15).

### 2.9 Validation in external datasets

The pipeline was applied to two independent snRNA-seq datasets (Supplementary Methods S16): the prefrontal cortex (BA9) data of Pineda et al. [30] (external dataset 1; 63 donors with ALS, FTLD, or no neurological disease [pathologically normal, PN]), and the insular cortex data of Rexach et al. [31] (external dataset 2; 37 donors with AD, bvFTD, PSP, or no neurological disease [controls]). Nuclei kept by the authors’ published quality control were mapped to the SEA-AD taxonomy by MapMyCells [32]. CEP-IP with CMRF was run with the R package cepip (0.1.0) at the SEA-AD settings on the PVM_3 and HKG8 arms. The DFGs replicated if at least five of the eight mitochondrial genes passed tier 1 (r_s_≥0.6 and *τ*_b_≥0.5) with *MT-CO3*. Module post-IP% was compared between diagnoses only if the arm reached an IPRS of at least 0.60 [18]. Each diagnosis was compared with controls by OLS. In external dataset 1, PMI was available for only 8 of the 13 PN donors. Thus, the primary model was adjusted for age at death and sex (cov2), and a secondary model added PMI (cov3). In external dataset 2, the model was adjusted for age at death, sex, and PMI (cov3). The stability of each β was tested post hoc by leave-one-donor-out (LODO) refits and by a donor bootstrap stratified by diagnosis (10,000 resamples). The power of each comparison to detect the SEA-AD NeoC cognitive-status β (0.465) was also estimated.

### 2.10 Software environment and reproducibility check

The study comprised 13 analysis parts, where Parts 1-8 were run in Python (3.13.14) on Fedora 44, and Parts 9-13 in R (4.6.1; Bioconductor 3.23) on Windows 11. The software environment is deposited as a container recipe for Podman or Docker, which installs every package at the version recorded in the lock files (Data and Code Availability). To test reproducibility, the pipeline was rerun inside the container on Fedora 44, and every CSV and Excel table was compared cell by cell with the original results. Numbers agreed when their relative difference was within 10^-6^, and text had to match exactly. In the Python rerun of Parts 1 to 8, 9,253 of 9,262 output tables agreed; the nine that differed were Part 5 differential expression tables, where a few *p*-values and log_2_ fold changes shifted slightly, with no gene crossing the 0.05 cutoff. The count matrices that Part 8 exports as the input for downstream R analyses were identical to the original export.

In the R rerun of Parts 9 to 13 of the scripts, 1,150 of 1,203 tables matched the original results or differed by at most 0.1%, apart from differences in sign, row order, or layout. The other 53 that differed held secondary statistics that depended on random subsamples, permutations, or nearest neighbors, and none of these differences changed a significance call. The whole pipeline was also run with one command (seaad-run all) in a new folder that held only the inputs, and it yielded the same outcome.

Finally, a second person not involved in the analysis ran two checks on a Windows 11 laptop. The reproducibility script of the R package cepip agreed with all 698 published values, and the container rerun of R Parts 9 to 11 yielded identical clinical associations, with differences only in secondary UMAP-based and permutation statistics (Supplementary Methods A3). The analyses of the two external datasets (Section 2.9) were each deposited as a container, and reruns by the author and by a second person on Windows 11 reproduced their results (Supplementary Methods S16). The comparison of every table is deposited with the code (Data and Code Availability).

## 3. Results

### 3.1 Selection of Mic.13-like cells and the *MT-CO3* module

M4 (Mic.13-like) was mapped on the SEA-AD snRNA-seq dataset according to five different cutoffs (top 2.5%, 2.9%, 3.0%, 3.1%, and 5% of PVM_1 to PVM_4, Lymphocyte, and Monocyte populations; Supplementary Table S1). The top 3% was selected due to: (i) M4 cutoff of 3.0% was within the admissible range of an objective selection rule, where no supertype other than PVM_3 or PVM_4 (the disease-associated and lipid-associated microglia of the SEA-AD taxonomy, respectively [19]), had >1% of its cells selected (Supplementary Figure S1A); (ii) Differentially expressed gene (DEG) screening of the selected against the non-selected PVM_3 cells recovered Mic.13 markers that were not used for scoring, 24 of 68 in NeoC (*p*=2.4×10^-9^) and 13 of 68 in NonNeoC (*p*=2.6×10^-5^) (Supplementary Table S2). This yielded n=3,412 and n=2,704 cells in NeoC and NonNeoC, mostly PVM_3 cells (Figure 1A and 1B). As Mic.12 and Mic.13 of Green et al. corresponded to Micro-PVM_3, the disease-associated microglia (DAM) supertype of the SEA-AD taxonomy [19], PVM_3 was prioritized in this study.

**Figure 1.**
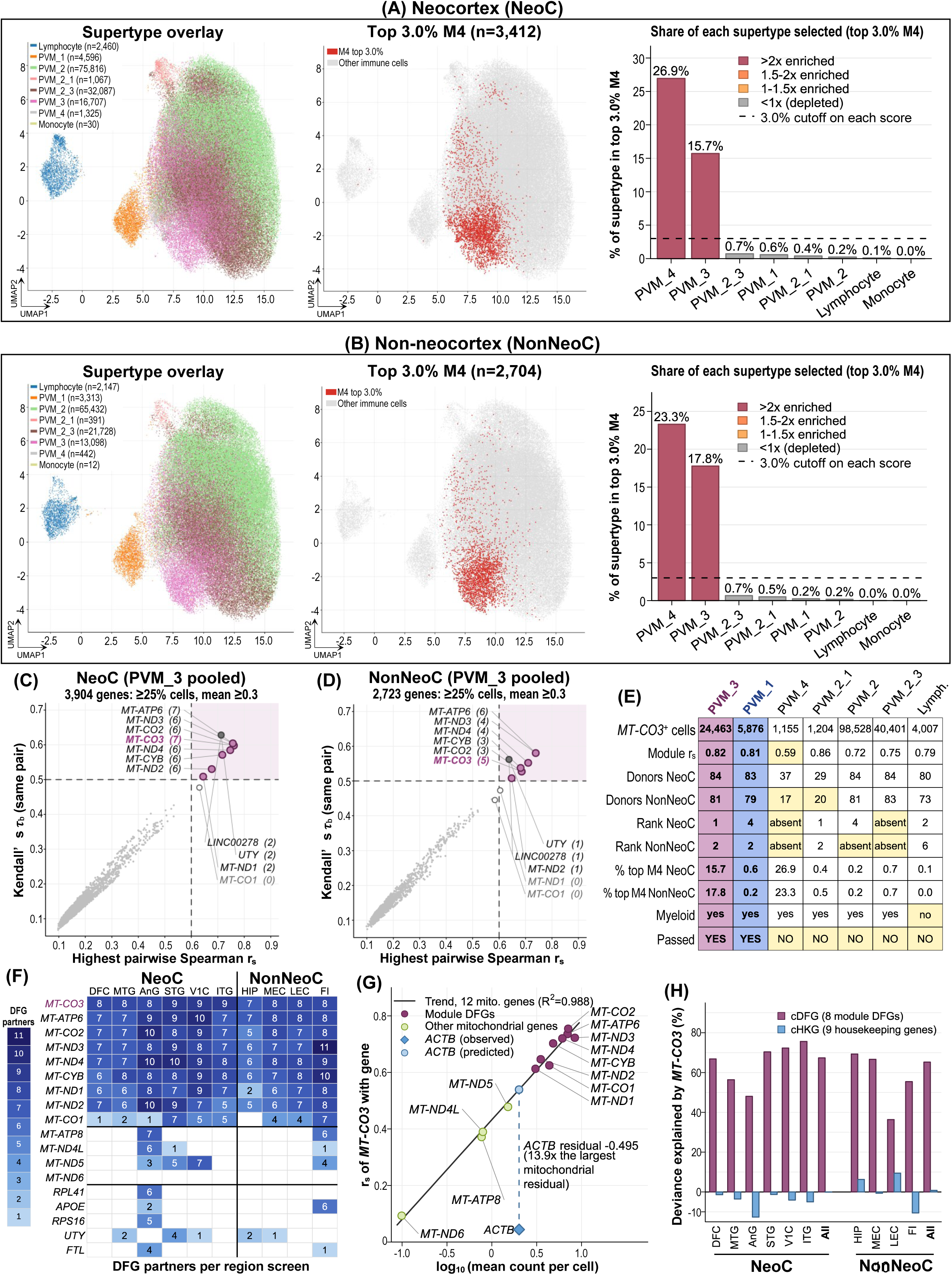
Selection of M4 (Mic.13-like) cells, choice of target and control supertypes, and the *MT-CO3* module. (A, B) Left, UMAP of immune cells colored by supertype. Middle, cells in the M4 top 3% (red; n=3,412 and 2,704). Right, percentage of each supertype’s cells selected, colored by enrichment (percentage selected divided by the 3.0% cutoff); dashed line, the 3.0% cutoff. (C, D) Pooled GOI-DFG screens of 2,500 PVM_3 cells per region set. Dashed lines are DFG gates; shaded indicates both gates passed; plum circles indicate mitochondrial genes with DFG partners; dark gray circles are *UTY* and *LINC00278* (overlapping); open circles are other mitochondrial genes; numbers in brackets indicate DFG partners. (E) Supertype criteria; yellow indicates criterion not met. *MT-CO3*⁺ cells, number of cells with *MT-CO3*>0; Module r_s_ indicates r_s_ of *MT-CO3* with cDFG; both region sets combined; Rank indicates rank of *MT-CO3* in the pooled screen; % top M4, as in A. Monocyte (<50 cells) not shown. (F) DFG partners per gene in the regional screens of M4 cells (blank, none). (G) r_s_ between *MT-CO3* and each mitochondrial gene against log_10_ mean counts per cell in PVM_3 cells with *MT-CO3*>0, with the trend over 12 other mitochondrial genes; *ACTB* observed (diamond) and predicted (light blue circle). (H) Deviance explained (DE) by *MT-CO3* in cDFG and cHKG in GAMs of M4 cells, relative to a donor-only model, per region and pooled (All).

Next, screening for pairwise genes (i.e., GOI-DFG pairs) that passed the expression and correlation gates was conducted in each region set separately. Both NeoC and NonNeoC showed multiple mitochondrial genes passing these cutoffs in the pooled screen of 2,500 subsampled PVM_3 cells (Figures 1C and 1D; Supplementary Table S3). *MT-CO3* was the top-ranked gene compared with 12 other mitochondrial genes and chosen as the GOI (Supplementary Methods S2; Supplementary Table S3). *MT-CO3* had no DFG partners in PVM_2_3 or PVM_4 in either region set (Figure 1E; Supplementary Table S4). A gene was kept as an *MT-CO3* partner DFG if it was *MT-CO3*’s DFG in at least 8 of the 10 brain regional screens in M4 cells, resulting in eight DFGs (*MT-ATP6*, *MT-CO1*, *MT-CO2*, *MT-CYB*, *MT-ND1*, *MT-ND2*, *MT-ND3*, and *MT-ND4*) (Figure 1F).

To examine whether confounding factors including ambient RNA could produce the *MT-CO3* module, the following tests were conducted in pooled PVM_3 cells with *MT-CO3*>0 (Supplementary Figure S1B to S1E; Supplementary Table S5): (i) None of 295 matched gene panels of similar expression levels (maximum r_s_=0.348) reached the observed module’s r_s_=0.817; (ii) In 185 libraries, module r_s_ within each library was inversely associated with the library’s intronic:exonic ratio (lower ratio, more ambient RNA; r_s_=-0.339, *p*=2.3×10^-6^); (iii) With library median mitochondrial percentage regressed out of both, partial r_s_ was -0.077 (*p*=0.296); (iv) In quartiles of the nucleus-level ambient index, module r_s_ (0.793 to 0.831) did not rise consistently as the ambient index rose; (v) After ambient RNA removal with DecontX, module r_s_ changed from 0.817 to 0.802, within a tolerance of 0.05 (Supplementary Table S6). Collectively, these tests did not suggest ambient contribution to the module, but the library-level analysis could not separate ambient from cellular mitochondrial RNA (this was examined further in Battery 1).

Among the seven supertypes in Figure 1E, only PVM_3 and PVM_1 passed the set criteria: having at least 25 donors in each of NeoC and NonNeoC, presence of *MT-CO3* partner DFGs within a supertype’s pooled screen, module r_s_≥0.6, and of myeloid lineage (Figure 1E; Supplementary Tables S1, S4, and S5). PVM_3 and PVM_1 were selected as the target and control supertype, respectively. To test whether the *MT-CO3*/DFG module was dependent on any specific brain region, the GOI-DFG screen was performed separately in each of the 10 regions in M4 cells, where *MT-CO3* showed 7 to 9 DFG partners in all 10 regions (Figure 1F; Supplementary Table S3). In leave-one-region-out sensitivity analysis (Supplementary Figure S1F), the pooled r_s_ of *MT-CO3* with cDFG changed by at most 0.014 except in NonNeoC without MEC, where r_s_ dropped to 0.747, more than twice the matched-null maximum of 0.348. Hence, the *MT-CO3*/DFG module did not depend on any specific brain region.

In order to assess whether the *MT-CO3* and DFG correlation occurred simply due to gene abundance, a linear model of r_s_ (*MT-CO3* r_s_ with the other 12 mitochondrial genes) as a function of log_10_ mean UMI count per cell was constructed (Figure 1G; Supplementary Table S5). The r_s_ followed mitochondrial abundance closely (R^2^=0.988), and the eight module DFGs were the most abundant mitochondrial genes. In contrast, for a control gene *ACTB*, the model predicted r_s_=0.539 but the observed r_s_ was 0.044, a residual 13.9 times the largest residual of any mitochondrial gene, suggesting that abundance alone did not produce the correlation. Finally, cDFG and cHKG (*ACTB* and eight other housekeeping genes combined) were each fitted against *MT-CO3* with a GAM in M4 cells, per region and pooled per region set (Figure 1H; Supplementary Table S7). *MT-CO3* explained about 62% of the variation in cDFG on average (DE=36.3-75.6% in the 12 fits), but did not explain cHKG variation (DE=-12.7% to 9.4%).

### 3.2 Identification of Spearman-Kendall cell-level score (SKCS)-core cells and pitcher plot

Heatmap visualizations, when ordered according to ascending *MT-CO3* expression in the three main arms (M4, PVM_3, and PVM_1), showed a monotonically increasing trend of cDFG and each individual DFG expression in every region of NeoC and NonNeoC, and this was not the case in the control arm HKG8 (Supplementary Figures S2 and S3; Supplementary Table S7). Subsequent analyses focused on the region sets NeoC and NonNeoC, each divided into the three main arms (*MT-CO3*/cDFG modeling) and the control arm HKG8 (*ACTB*/cHKG8 modeling).

GAM’s DE was approximately 65-67% in both region sets (Figure 1H), and hence the next aim was to improve the modest DE. r_s_ and τ_b_ were decomposed into per-observation S_i_ and K_i_ for each cell within each arm and region set. Cells that passed the S_i_>0.05 and K_i_>0.5 thresholds were shortlisted as the SKCS-core cells: 1,419 of 2,130 (66.6%) M4 cells in NeoC and 1,303 of 1,987 (65.6%) in NonNeoC (Supplementary Table S7). Regarding why Si>0.05 was chosen, many cells had S_i_ near zero and formed a narrow peak in the S_i_ distribution, and S_i_>0.05 was the smallest cutoff (in steps of 0.01) that was above two estimates of the upper end of this peak in both region sets (Supplementary Figure S4; Supplementary Table S7). K_i_>0.5 corresponds to GOI/cDFG concordance of above 75%.

In M4, restricting *MT-CO3*^+^ cells to SKCS-core improved DE in both NeoC (from 67.4% to 87.7%) and NonNeoC (from 65.2% to 87.8%), but the cell population was discontinuous along the fitted GAM curve due to the fixed S_i_ and K_i_ thresholds (Figure 2). When plotted on a K_i_ (y-axis) versus S_i_ (x-axis) graph, SKCS-core cells formed a distinctive bulb-like edge in the top right quadrant, while the non-SKCS-core population formed a stalk-like edge, producing an overall shape that resembles the outline of a tropical pitcher plant. Hence, the plot was termed a pitcher plot.

**Figure 2.**
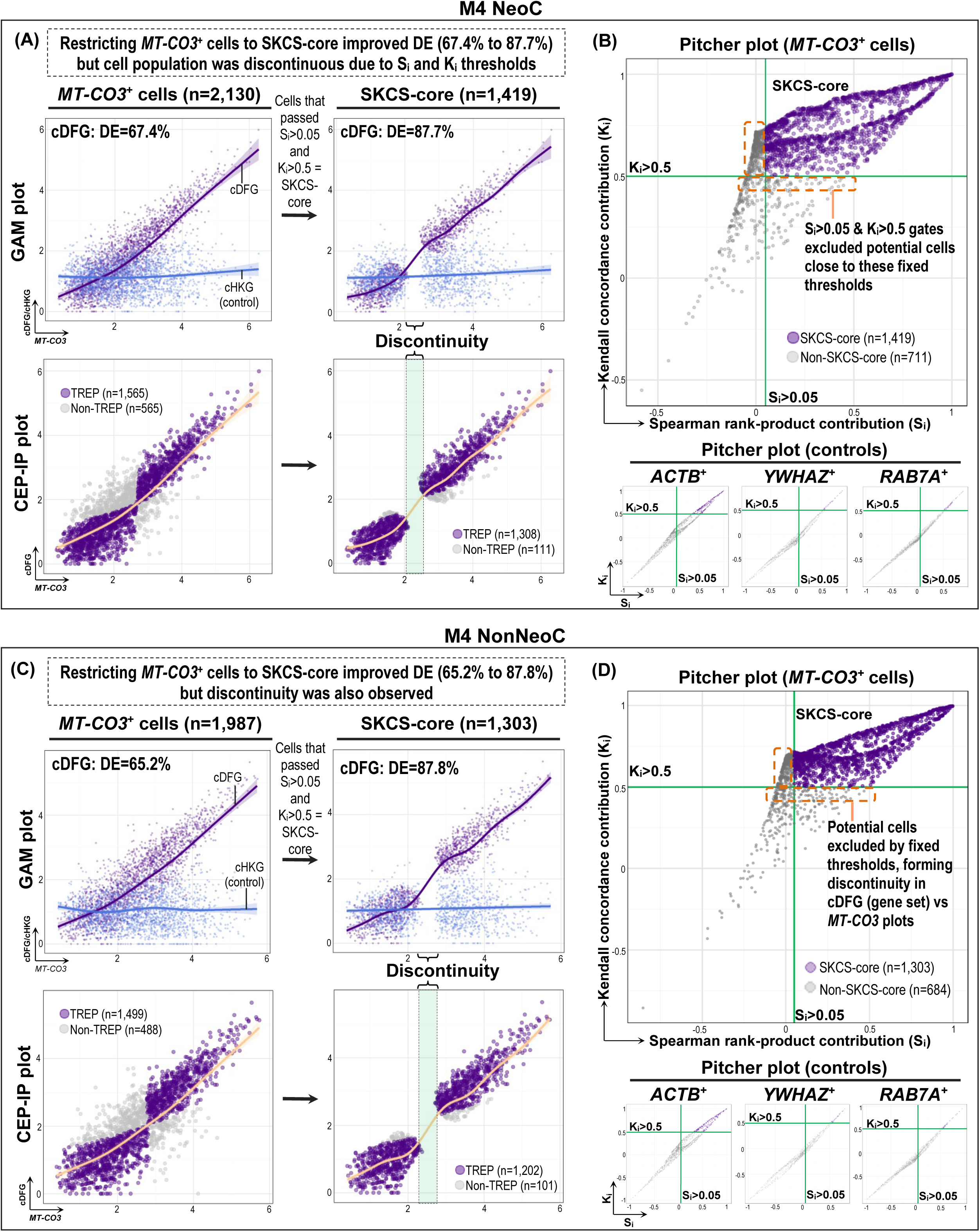
GAM, CEP-IP and pitcher plots of *MT-CO3*^+^ and SKCS-core M4 cells. (A [NeoC] and C [NonNeoC]) GAM (top) and CEP-IP plots (bottom) of *MT-CO3*^+^ cells (left; NeoC: 2,130 cells from 78 donors; NonNeoC: 1,987 cells from 80 donors) and SKCS-core cells (right; NeoC: 1,419 cells from 74 donors; NonNeoC: 1,303 cells from 79 donors). DE, the deviance in cDFG explained by *MT-CO3*; cHKG (control), the composite of the nine housekeeping genes. TREP cells, those best explained by the GAM fit (orange curve), were defined separately for *MT-CO3*^+^ and SKCS-core cells. Shaded indicates the discontinuous region left by the SKCS-core. (B [NeoC] and D [NonNeoC]) Pitcher plots of *MT-CO3*^+^ cells (top), with the SKCS-core in the top-right quadrant beyond the S_i_>0.05 and K_i_>0.5 gates (green lines; orange dashed boxes indicate cells just outside the gates), and control panels (bottom) with *ACTB*, *YWHAZ*, or *RAB7A* as the GOI and the other eight housekeeping genes as the composite, in HKG8 cells expressing that gene.

In the *ACTB*/cHKG8 control cells, the pitcher plots showed less distinctive bulbous boundaries, and most cells fell outside of the top right quadrant (SKCS-core: 17.5% and 18.4% of *ACTB*^+^ cells in NeoC and NonNeoC), also when *ACTB* was replaced by *YWHAZ* or *RAB7A* as the GOI (SKCS-core in both NeoC and NonNeoC: 6.3% to 8.2%) (Figure 2B and 2D; Supplementary Table S7). PVM_3, PVM_1, and HKG8 are shown in Supplementary Figure S5. In HKG8, DE also rose in the SKCS-core (from 11.3% to 78.5% in NeoC and from 13.0% to 64.8% in NonNeoC), as the S_i_/K_i_ gates select concordant cells regardless of the genes.

### 3.3 CMRF reclaims monotone-coherent cells

SKCS-core cells passed both the S_i_ and K_i_ gates and were taken as monotonic, but the fixed thresholds excluded cells that may be members of the same monotonic relationship. Hence, the next aim was to objectively recover the cells that aligned with the *MT-CO3*/cDFG monotonicity.

To achieve this, the first step was to identify the cDFG’s IP_y_ that separates SKCS-core cells with cDFG below IP_y_ (i.e., pre-IP; *MT-CO3*^low^/cDFG^low^) from those at or above it (i.e., post-IP; *MT-CO3*^hi^/cDFG^hi^); each population was also decomposed into CEP-IP’s TREP and non-TREP populations. IP_x_, determined on the *MT-CO3* axis (*ACTB* in HKG8), was computed in each arm and region set separately (Supplementary Figure S6). In the main arms, IP_x_ ranged from 2.383 to 2.803 with IPRS of 0.761-0.819 (strong or very strong tier), whereas in HKG8 the 95% CIs of IP_x_ were 10-18 times wider (IPRS: 0.571-0.639) (Supplementary Table S8). Each IP_x_ was then transformed into IP_y_, the fitted cDFG value at IP_x_ on the GAM curve, i.e., IP_y_ = f(IP_x_). On the pitcher plot, the two distinctive spines in the SKCS-core quadrant represented the pre-IP and post-IP populations (Figure 3 and Supplementary Figure S7; Supplementary Table S7).

**Figure 3.**
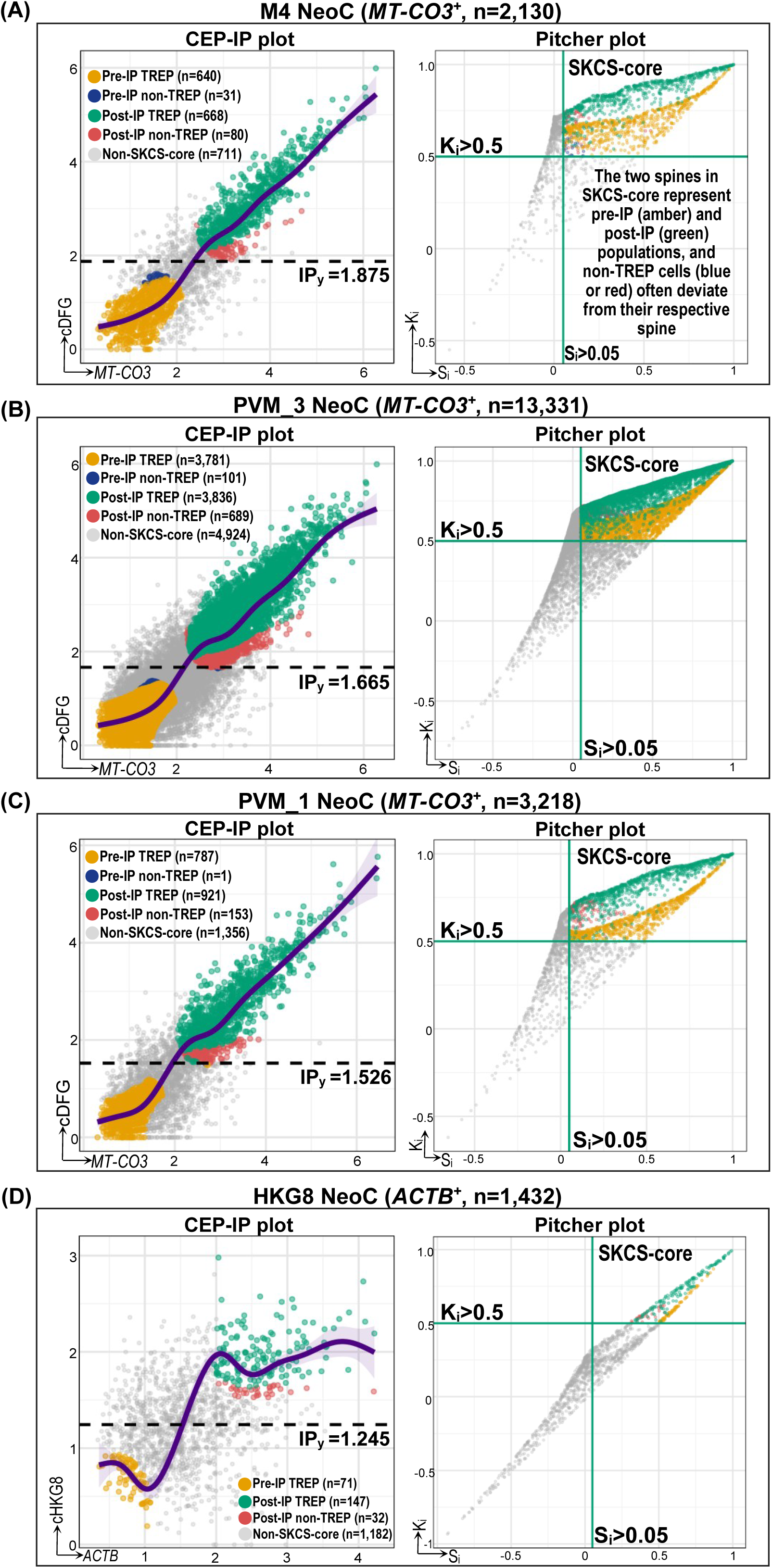
CEP-IP and pitcher plots of M4 (A), PVM_3 (B), and PVM_1 (C) NeoC *MT-CO3*^+^ cells and HKG8 (D) NeoC *ACTB*^+^ cells. Left, composite (cDFG; cHKG8 in HKG8) against GOI expression; right, K_i_ against S_i_. Cells: M4 2,130; PVM_3 13,331; PVM_1 3,218; HKG8 1,432. IP_y_ = f(IP_x_), the population-level GAM fit of all GOI^+^ cells at the GOI inflection point IP_x_ (Supplementary Figure S6), separates pre-IP and post-IP cells (dashed line); purple curve indicates the GAM fit of the SKCS-core, which defines TREP and non-TREP cells. Cell group colors: pre-IP TREP (amber), pre-IP non-TREP (blue), post-IP TREP (green), post-IP non-TREP (red), and non-SKCS-core (gray). The two spines of the SKCS-core quadrant are the pre-IP and post-IP groups.

In order to limit the influence of outliers, the expression values were transformed into copula space (rank-based) u and v, with IP_x_ and IP_y_ transformed into IP_u_ and IP_v_, while S_i_ and K_i_ were transformed into u_SK_ and v_SK_ (Figure 4 and Supplementary Figures S8-S10; Supplementary Table S9). In the copula space pitcher plot (v_SK_ versus u_SK_) of the main arms but not in the control arm, the bulbous boundaries dominated the pitcher plot with both pre-IP and post-IP spines being more prominent than their score space version, as the SKCS-core comprised the majority of cells in the main arms (57.9% to 66.6%) but not in HKG8 (17.5% in NeoC and 18.4% in NonNeoC) (Figure 4C and 4D).

**Figure 4.**
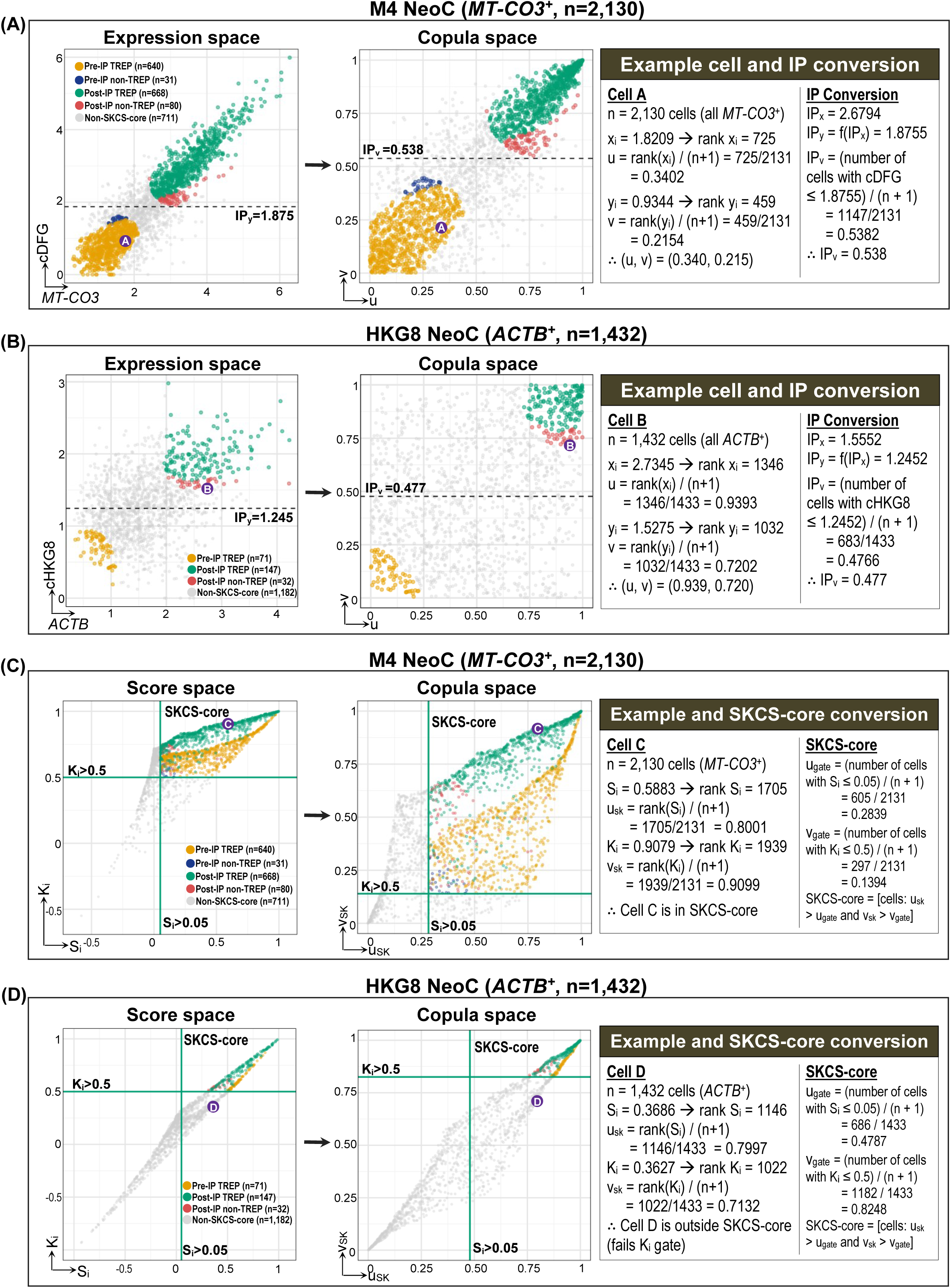
M4 NeoC *MT-CO3*^+^ cells (A, C) and HKG8 NeoC *ACTB*^+^ cells (B, D) in expression, score, and copula space. (A, B) Expression space (composite against GOI; left) and copula space (ranks divided by n + 1; middle). The right-hand box converts one cell (cell A or B, circled) and the IP to copula space: IP_x_ yields IP_y_ = f(IP_x_), whose copula coordinate is IP_v_ (dashed line). (C, D) Score space (K_i_ against S_i_; left) and copula space (middle). The right-hand box converts one cell (cell C or D) and the gates S_i_=0.05 and K_i_=0.5 to u_gate_ and v_gate_, so that the SKCS-core holds the cells with u_SK_ > u_gate_ and v_SK_ > v_gate_. S_i_ in C and D is computed from the copula coordinates, 4(u − 0.5)(v − 0.5). As the ranks are divided by n + 1 instead of being scaled by n − 1, it equals the S_i_ of Figure 2 multiplied by ((n − 1)/(n + 1))², which is 0.998 for M4 and 0.997 for HKG8 (Supplementary Methods S6).

Subsequently, the CMRF pipeline was performed in three stages (Figures 5 and 6; Supplementary Table S10). In Stage 1, SKCS-core pre-IP and post-IP TREP cells were used as the calibration population to establish the GAM spline, g(u) = E[v|u] (Figure 6), and the band was widened at IP_u_ (Supplementary Figure S11). SKCS-core and non-SKCS-core cells within the band were subgrouped as Original+Refined (OriRef) or candidate cells, respectively, and SKCS-core cells outside the band as Original-only. In Stage 2, candidates were removed one at a time via the RSk-fCV gate until an iteration passed the dual stopping rule. In Stage 3, OriRef and Recovered (candidates that survived all RSk-fCV iterations) formed the final set of monotonic Module cells, while Original-only and Excluded formed the non-Module cells retained for downstream analyses.

**Figure 5.**
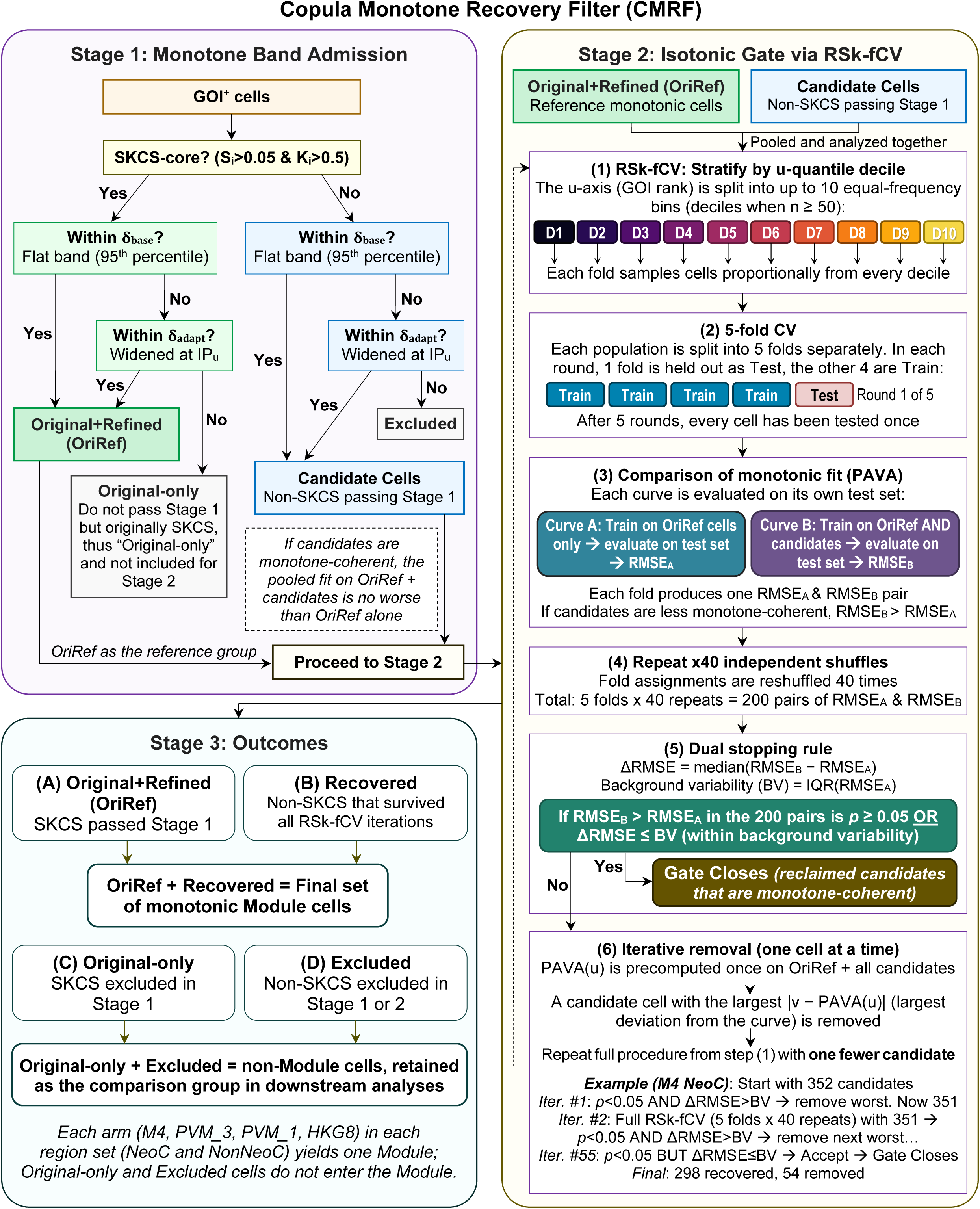
Workflow of the copula monotone recovery filter (CMRF). Stage 1: In copula space, GOI⁺ cells within the flat band δ_base_ or the band widened at IP_u_, δ_adapt_, are admitted; the admitted SKCS-core cells form the reference set (Original + Refined, OriRef), SKCS-core cells not admitted are Original-only, admitted non-SKCS cells are candidates, and the other non-SKCS cells are Excluded. Stage 2 (RSk-fCV): isotonic (PAVA) fits on OriRef (Curve A) and on OriRef plus candidates (Curve B) are compared by five-fold cross-validation stratified by u decile and repeated 40 times (steps 1 to 4); the dual stopping rule is applied (5); and the candidate with the largest |v - PAVA(u)| is removed (6), one per iteration, until the gate closes. Stage 3 assigns each cell to one of four classes (A to D); OriRef plus Recovered forms the Module. Bottom right, M4 NeoC example: of 352 candidates, 54 were removed before the gate closed at iteration 55, and 298 were recovered. Abbreviations: BV, background variability; PAVA, pool-adjacent-violators algorithm; RMSE, root mean squared error; RSk-fCV, repeated stratified k-fold cross-validation.

**Figure 6.**
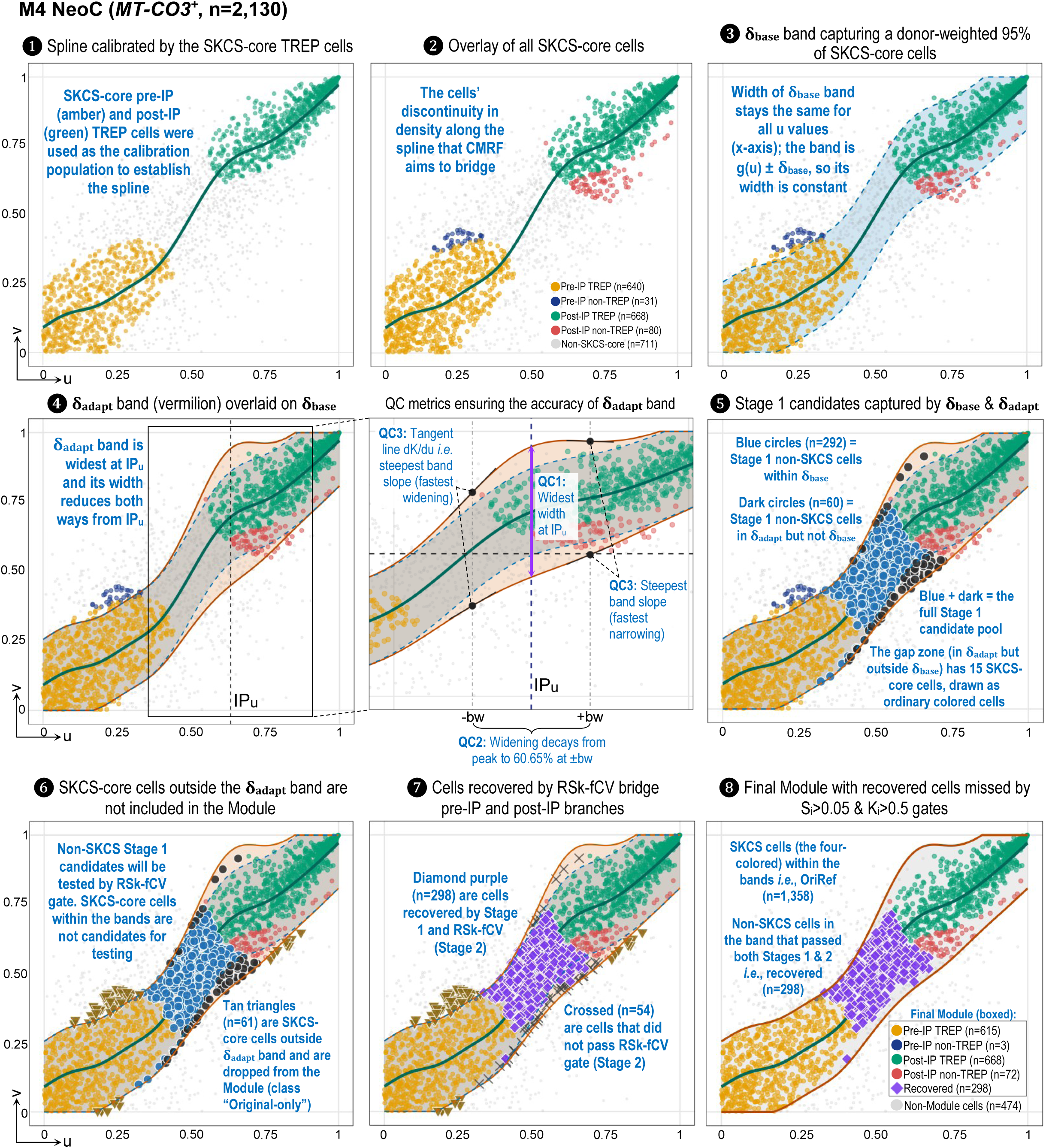
Construction of the CMRF tolerance band and recovery of monotone-coherent cells, M4 NeoC (n=2,130). Copula coordinates u and v of *MT-CO3* and cDFG, respectively. (1) Spline g(u) = E[v | u] fitted to the 1,308 SKCS-core TREP cells (640 pre-IP, 668 post-IP) with a donor random intercept and inverse-donor weights. (2) All 1,419 SKCS-core cells, with the gap in density along the spline. (3) Flat band g(u) ± δ_base_; δ_base_=0.1642, the donor-weighted 95th percentile of |v - g(u)| over the SKCS-core. (4) Adaptive band δ_adapt_(u) (vermilion), widest at IP_u_=0.633 (δ_max_=0.2463), with the kernel checks QC1 to QC3 (middle panel; Supplementary Figure S11). (5) Stage 1 candidates, all non-SKCS-core: 292 within δ_base_ (blue circles) and 60 within δ_adapt_ only (dark circles), 352 in total; the 15 SKCS-core cells in the widened zone are drawn as ordinary core cells. (6) The 61 SKCS-core cells outside δ_adapt_ (tan triangles) leave the module as Original-only. (7) The RSk-fCV gate recovered 298 candidates (purple diamonds) and removed 54 (crosses). (8) Final module: 1,358 OriRef cells and 298 recovered cells; the other 474 cells (413 Excluded and 61 Original-only) are not in the module.

In RSk-fCV of M4 NeoC cells (Figure 7), the gate closed at iteration 55, when ΔRMSE (0.0044) fell below BV (0.0046), after 54 of 352 candidates (15%) were removed; 298 cells were recovered, yielding a Module of 1,656 cells. In all eight arm and region set combinations, the gate closed on ΔRMSE≤BV with *p*<0.05. The other arms recovered 242 cells (M4 NonNeoC), 1,351 and 1,089 (PVM_3), and 381 and 318 (PVM_1) in NeoC and NonNeoC (Supplementary Figure S12). In HKG8, the spline was not monotone and more cells were recovered than OriRef held (287 against 239 in NeoC, and 240 against 191 in NonNeoC).

**Figure 7.**
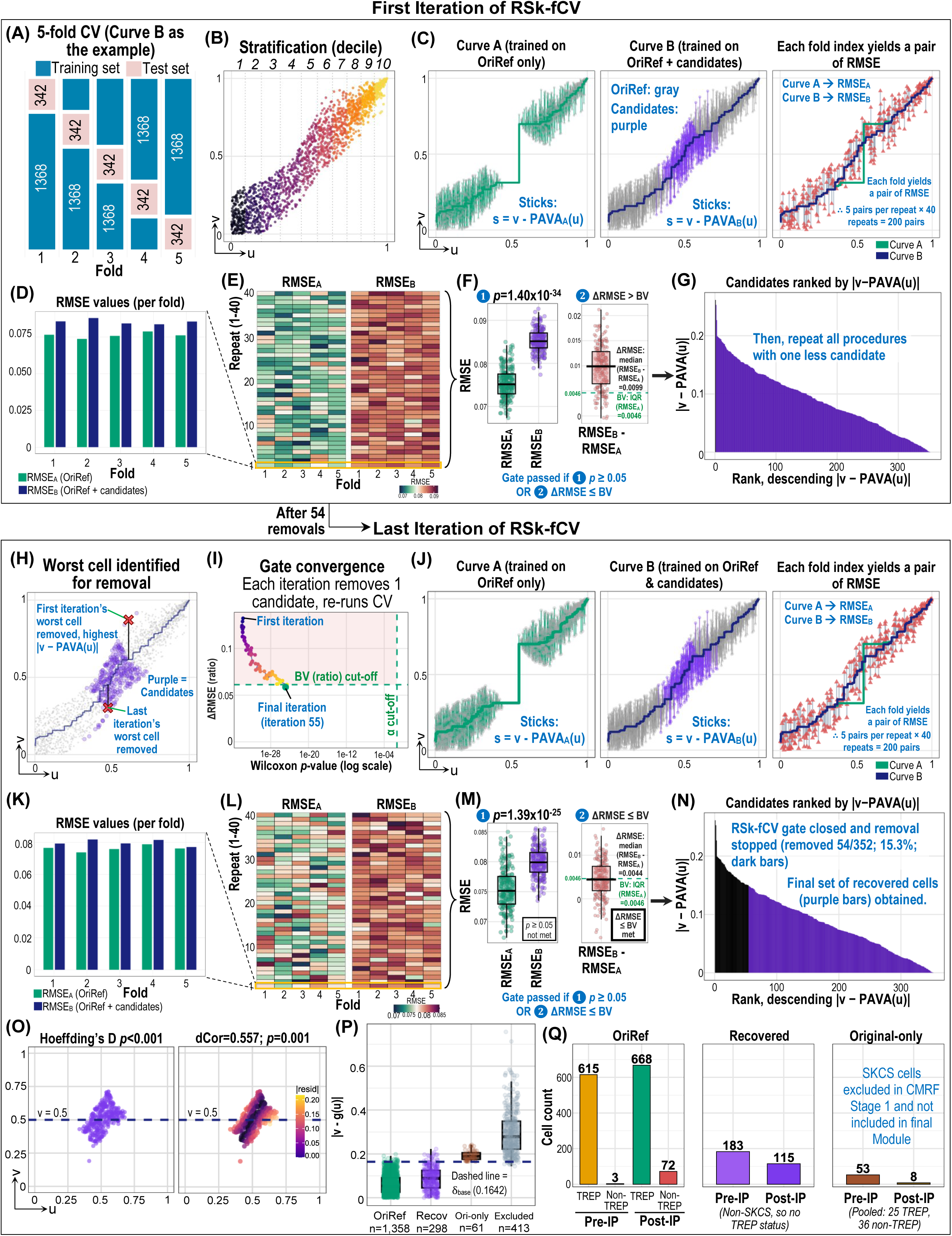
The RSk-fCV gate (M4 NeoC example). Stage 2 tests whether adding the Stage 1 candidates degrades the monotone fit of the CMRF spine. (A to G) First iteration. The pool was 1,358 OriRef cells plus 352 candidates (A, shown for Curve B; Curve A’s folds are drawn from the OriRef cells alone). Folds were stratified by u decile (B). Curve A is an isotonic fit trained on OriRef only, and Curve B on OriRef plus candidates; sticks are residuals to each curve (C). C3 and J3 are schematic: both curves were drawn on one shared fold to compare them on the same cells. The gate runs two separate cross-validations and pairs RMSE_A_ with RMSE_B_ by repeat and fold. Five folds x 40 repeats yield 200 paired RMSE values (D and E; D and K show repeat 1, the highlighted heatmap row). The gate continued removing while *p*<0.05 and ΔRMSE = median(RMSE_B_ - RMSE_A_) exceeded the background variability BV = IQR(RMSE_A_) =0.0046 (F; first iteration, *p*=1.40×10^-34^ and ΔRMSE=0.0099). Each iteration removed the candidate with the largest |v - PAVA(u)| (G and H). (I to N) Convergence and closure. After 54 removals (15.3% of 352), the gate closed at iteration 55 on the effect-size arm: ΔRMSE fell to 0.0044, below BV, while the *p*-value remained significant (I, M, and N). (O to Q) The 298 Recovered cells. Their u and v remain dependent (O), spine residuals increased monotonically over OriRef, Recovered, Original-only, and Excluded cells (P; Jonckheere-Terpstra *p*<0.001), and the final module composition is shown in Q.

### 3.4 Validation of CMRF-derived Recovered cells

In copula space pitcher plots, Recovered cells bridged the pre-IP and post-IP spines, forming a complete bulbous boundary in the three main arms and both region sets (Figure 8A). Conversely, in HKG8, the Recovered cells were located away from both spines. In the main arms, the vast majority of Recovered cells failed the S_i_>0.05 gate (95.0-99.7%) and few failed the K_i_>0.5 gate (0.3-5.8%), against 28-29% that failed the S_i_>0.05 gate and 100% that failed the K_i_>0.5 gate in HKG8 (Figure 8C). This indicates that in the main arms, CMRF reclaimed cells whose K_i_ concordance nearly always passed the gate but whose S_i_ rarely did.

**Figure 8.**
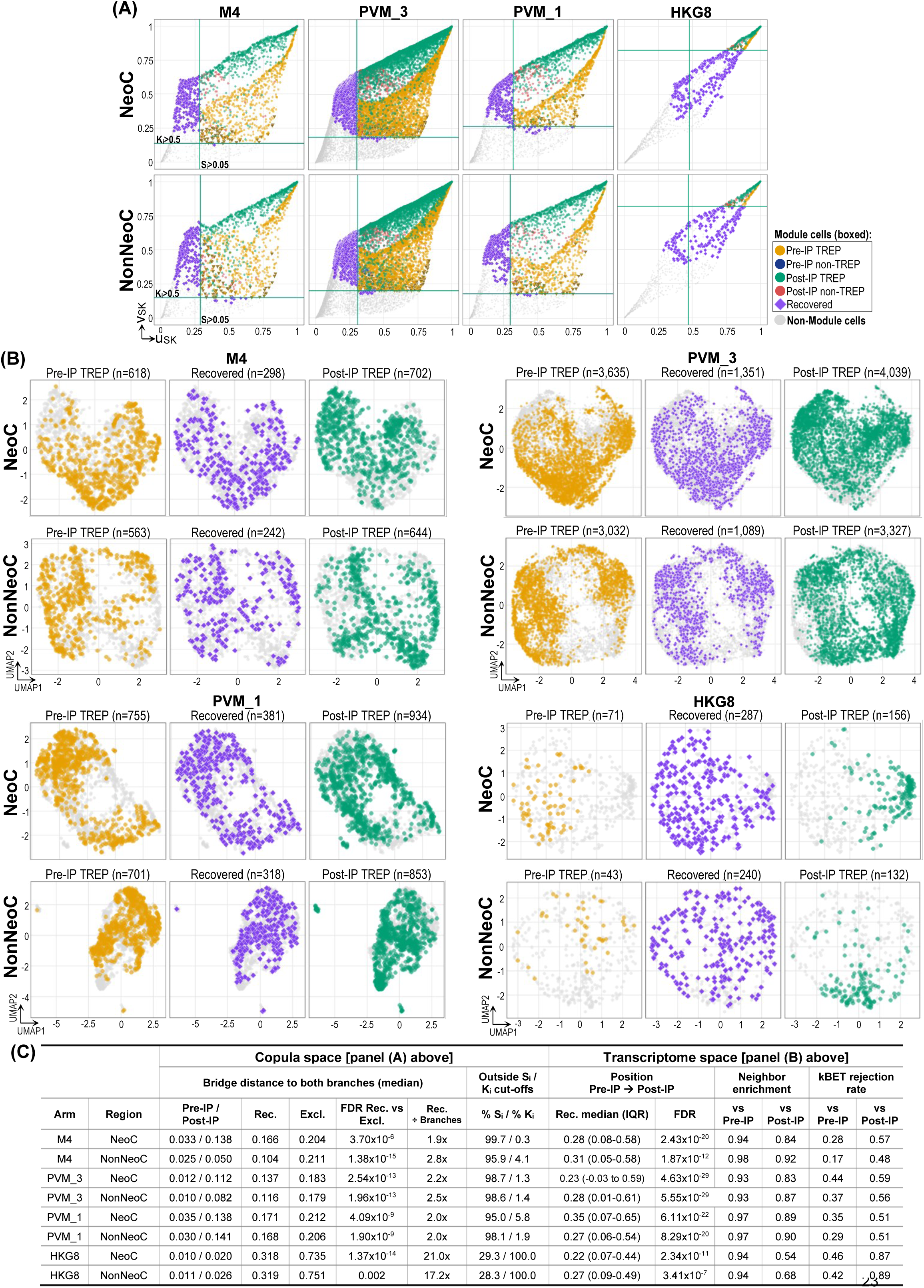
Validation of the Recovered cells in copula and transcriptome space. (A) Pitcher plots in copula score space (u_SK_, v_SK_) per arm and region set: final Module cells (pre-IP and post-IP, TREP and non-TREP, and Recovered) and non-Module cells (gray; tan triangles, Original-only); lines, the S_i_>0.05 and K_i_>0.5 gates. (B) UMAPs from the raw counts of each arm’s GOI⁺ cells: pre-IP TREP, Recovered, and post-IP TREP cells against the other cells (gray); TREP was recomputed on the GAM refitted to the final Module, and the TREP groups hold OriRef cells only. (C) Summary per arm and region set; medians are over cells, and FDRs are from donor-level tests (Supplementary Methods S7). Copula space: median bridge distance (the larger of a cell’s distances to the nearest pre-IP and post-IP TREP cell) of TREP, Recovered, and Excluded cells, the FDR of Recovered against Excluded, and the ratio of the Recovered median to the average of the pre-IP and post-IP TREP medians; percentages of Recovered cells with S_i_≤0.05 and with K_i_≤0.5. Transcriptome space: median (IQR) position of Recovered cells from the pre-IP (0) to the post-IP (1) TREP centroid, with the FDR of the ordered trend; mean neighbor enrichment (1 indicates random mixing) and kBET rejection rate (0 indicates complete mixing) against each TREP group over 50 nearest neighbors.

In UMAP space (Figure 8B), the Recovered cells in the main arms occupied the same neighborhoods as the pre-IP and post-IP TREP cells (mean neighbor enrichment 0.83-0.98, where 1 indicates random mixing) and resided between the pre-IP and post-IP TREP cells (median position=0.23-0.35, where 0 is the pre-IP and 1 the post-IP TREP centroid; ordered trend FDR≤1.87×10^-12^), closer to pre-IP (kBET rejection rate 0.17-0.44 against pre-IP and 0.48-0.59 against post-IP TREP cells). HKG8 Recovered cells also sat between the pre-IP and post-IP TREP cells, but far from both TREP spines in copula space and separated from the post-IP TREP cells in UMAP space (Figure 8C; Supplementary Table S10). Collectively, Recovered cells in the main arms were consistent with a continuum from pre-IP to post-IP that the S_i_/K_i_ gates had excluded.

### 3.5 Differential expression between Module and non-Module cells

Apart from mitochondrial genes, no gene was specific to the Module (Supplementary Table S11). The only exception came from the post-IP against pre-IP comparison: 13 of 14 mostly neuronal genes rose in post-IP Module cells, because these nuclei held about 30% less of their own RNA, which raised the share of the neuronal RNA they carried. In PVM_3, 86 (NeoC) and 99 (NonNeoC) genes differed between post-IP Module and non-Module cells (e.g., *APOE*, *SPP1*, and *FTL*), but 147 of the 156 non-mitochondrial genes tested changed in the same direction in HKG8 control. Of the 368 non-mitochondrial PVM_3 genes that differed between post-IP and pre-IP Module cells, 306 also changed in the same direction in HKG8. The 14 genes that differed from HKG8 in both M4 and PVM_1 were mostly neuronal (e.g., *RBFOX1* and *CNTNAP2*), and short neuron-specific genes (e.g., *SNAP25*) rose with them. Post-IP Module nuclei held 28-33% fewer non-mitochondrial UMIs than pre-IP Module nuclei in M4, PVM_3, and PVM_1 (all FDR≤3.00×10^-5^), which non-Module nuclei did not show, while their neuronal UMIs per nucleus changed little (fold 0.87-1.11).

### 3.6 Association of Module post-IP% with clinical parameters

Pooled over donors, Module post-IP cells made up 5.1% and 5.9% of the arm’s supertype cells in M4 NeoC and NonNeoC, respectively, while the percentage was higher in PVM_3 (29.4% and 31.4%) and PVM_1 (25.8% and 31.6%), and lower in HKG8 (1.7% and 1.9%) (donors of Figure 9; Supplementary Figure S13; Supplementary Table S12).

**Figure 9.**
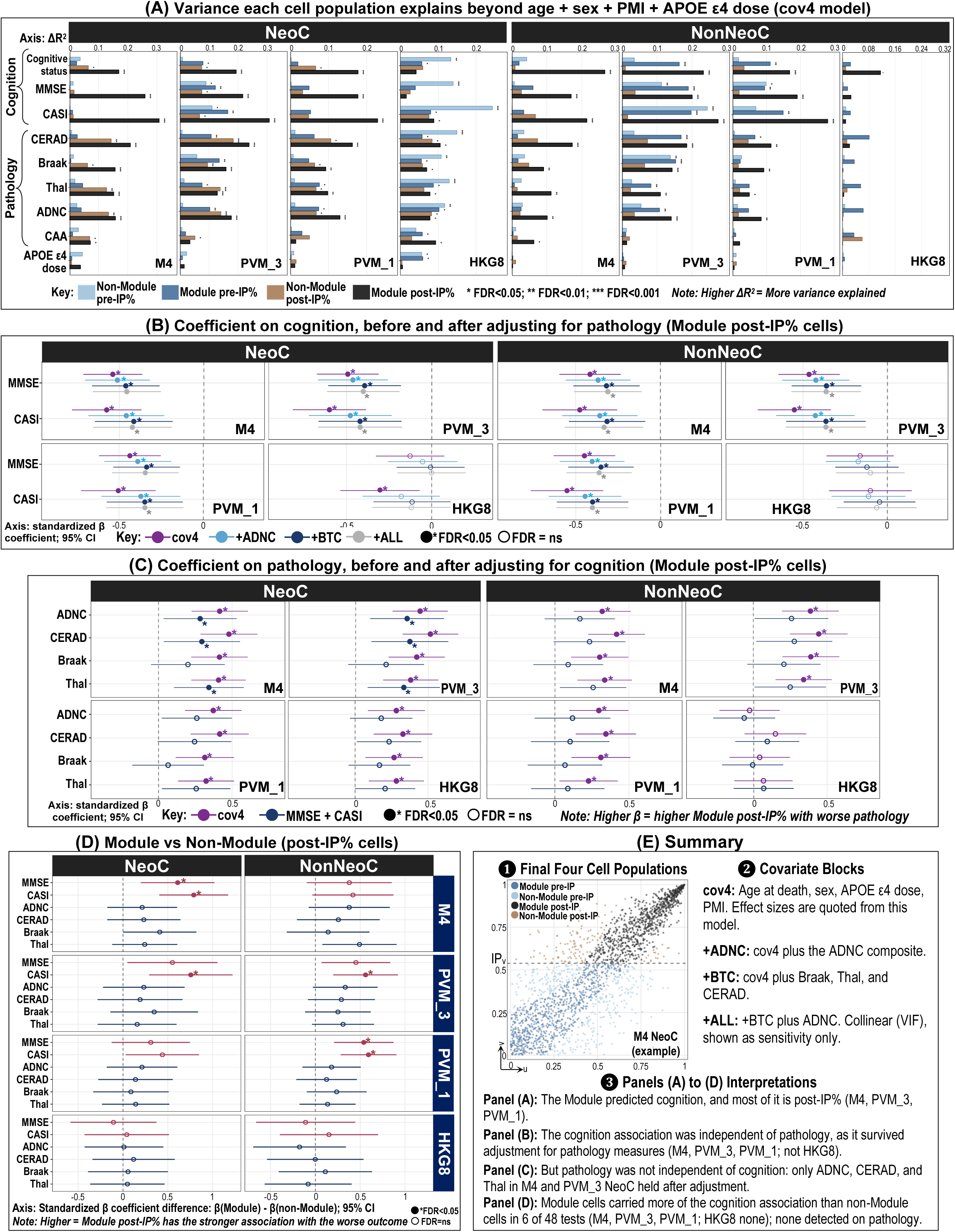
Association of Module post-IP% with cognition and pathology. Donor-level models for four arms and two region sets (84 NeoC and 83 NonNeoC donors); predictors are donor percentages of the arm’s supertype cells. (A) ΔR^2^ beyond age at death, sex, PMI, and *APOE* ε4 dose (cov4 model; APOE ε4 dose left out when it was the outcome) of the donor percentage of each of four cell populations on nine outcomes. * FDR<0.05, ** FDR<0.01, *** FDR<0.001 (BH over 15 outcomes, including six negative-control outcomes not shown). (B) Standardized β of Module post-IP% on MMSE and CASI at cov4 and after adding ADNC (+ADNC), Braak, Thal, and CERAD (+BTC), or all four (+ALL; collinear, sensitivity only); lower β indicates worse cognition. (C) Standardized β of Module post-IP% on ADNC, CERAD, Braak, and Thal at cov4 and after adding MMSE and CASI; higher β indicates worse pathology. In B and C, bars are 95% CIs (β±1.96 SE) and filled circles mark FDR<0.05. (D) Standardized β of Module minus non-Module post-IP% in one cov4 model, with MMSE and CASI multiplied by -1, hence higher values indicate a stronger Module association with the worse outcome; 69 NeoC and 68 NonNeoC donors. (E) Summary of the results: the four cell populations in the M4 NeoC copula plane with IP_v_, the covariate blocks, and the conclusion of each of panels A to D. Values are in Supplementary Table S12.

Associations of four donor percentages (Module pre-IP%, Module post-IP%, non-Module pre-IP%, and non-Module post-IP%) with clinical parameters were tested after adjustment for the four covariates, i.e., age at death, sex, PMI, and *APOE* ε4 dose (cov4 model). All significant associations of Module post-IP% were with worse outcomes, i.e., a higher Module post-IP% with worse cognition (dementia by cognitive status, lower MMSE and CASI) and worse pathology (higher CERAD, Braak, Thal, ADNC, and CAA); associations are in this direction hereafter unless stated. Module post-IP% was significant for all three cognition parameters (FDR<0.05) in all three main arms and both region sets, and its ΔR^2^ was the largest of the four donor percentages in all 18 comparisons (Figure 9A; Supplementary Table S12). In HKG8, it was also significant for CASI in NeoC and for cognitive status in NonNeoC.

In M4, Module post-IP% explained 17-31% of the variation in CASI and MMSE, while the other three donor percentages were not significant for either parameter. Module post-IP% was also significant for CERAD, Braak, Thal, and ADNC in all main arms and both region sets, for CAA in M4 only, and for all five pathology measures in HKG8 NeoC. Non-Module post-IP% was significant for CERAD, Braak, Thal, ADNC, and cognitive status in NeoC of all three main arms, for CAA in M4 and PVM_3 NeoC, for MMSE and CASI in PVM_3 NeoC only, and for no outcome in NonNeoC (Figure 9A).

These cov4 associations were then examined against tissue quality and other potential confounders (Supplementary Figure S14; Supplementary Table S12). Adding RIN, brain pH, and fresh brain weight to cov4 (cov7) reduced the ΔR^2^ of Module post-IP% on cognition to 39-79% of its cov4 value, but MMSE, CASI, and cognitive status remained significant in all three main arms and both region sets (Supplementary Figure S14A to S14C). In HKG8, cov7 removed the NeoC CASI association (FDR from 0.034 to 0.212), whereas NonNeoC cognitive status remained significant (FDR from 0.023 to 0.019). RIN and fresh brain weight were themselves associated with cognition or pathology (5 of 24 pairs, FDR<0.05; Supplementary Figure S14D), hence adjusting for them partly removed disease-related variation.

Furthermore, none of the 12 clinical, biological, and tissue covariates, added one at a time to cov4, removed the association of Module post-IP% with MMSE, CASI, or cognitive status in the main arms (largest attenuation 16.6%, for months from the last CASI; Supplementary Figure S14E). Of six technical covariates, the donor-mean mitochondrial fraction (the donor-level %MT, tested in Battery 3) or sequencing saturation removed 7 of the 18 cognition associations in the main arms (Supplementary Table S12).

To examine whether the significant cognition associations were simply due to pathology confounds, Module post-IP% in the cov4 model was also adjusted for ADNC (+ADNC), for Braak, Thal, and CERAD combined (+BTC), or for all four of these parameters (+ALL). In the three main arms, the associations with MMSE and CASI survived every pathology adjustment in both region sets (12 of 12 tests per arm, including the collinear +ALL sensitivity block; FDR<0.05), while the CASI association in HKG8 NeoC was lost in all three pathology-adjusted models (Figure 9B). Next, the reverse was investigated: when adjusted for MMSE and CASI, only 6 of the 24 pathology associations remained significant (FDR<0.05), all in M4 and PVM_3 NeoC for ADNC, CERAD, and Thal (Figure 9C). Collectively, the association of Module post-IP% with cognition survived adjustment for pathology, while its association with pathology mostly did not survive adjustment for cognition, except for amyloid-related pathology (ADNC, CERAD, and Thal) in M4 and PVM_3 NeoC.

Next, whether the clinical association belonged to the Module post-IP cells or to any post-IP cells of the same arm was assessed. Module post-IP% and non-Module post-IP% were tested together in the same cov4 model, and the difference between their two coefficients, Δβ = β(Module) - β(non-Module), was obtained, with MMSE and CASI sign-reversed, so that a positive value means a stronger association with the worse outcome. Module post-IP% showed stronger associations in 6 of 48 tests, all on MMSE or CASI (FDR≤0.018; Figure 9D), and none in HKG8 (FDR≥0.888). In the same models, Module post-IP% itself remained significant for MMSE and CASI in all three main arms and both region sets, and for 22 of the 24 pathology tests, whereas non-Module post-IP% was significant in none of the 48 tests (FDR≥0.240). The core results of this clinical outcomes arc are summarized in Figure 9E.

### 3.7 Module post-IP% versus competitors

Five competitors were selected for comparison with Module post-IP%, each computed per donor over all cells of the arm’s supertype: AUCell (AUC) [33], UCell (UC) [34], and AddModuleScore (AMS) [35], which measure how strongly each cell expressed the nine Module genes without boundaries or cell selection; cluster proportion (CP), the supertype’s cells as a percentage of all Micro-PVM cells; and changepoint split (CS), the most similar to CMRF’s Module post-IP%, which uses a standard breakpoint from segmented regression [36].

Firstly, Module post-IP% and the competitors were each fitted alone in the same cov4 model on the same donors. In PVM_3, Module post-IP% equaled or exceeded every competitor for MMSE and CASI in both region sets (e.g., NeoC ΔR^2^: 0.216 and 0.308 versus competitors 0.118-0.210 and 0.193-0.290) (Figure 10A; Supplementary Table S12), while competitors explained as much as or more than PVM_3 Module post-IP% for ADNC, CERAD, and Thal. In M4, Module post-IP% had a higher ΔR^2^ than every competitor for MMSE and CASI in NeoC only. In PVM_1, the three gene scores matched or exceeded Module post-IP% on most outcomes (e.g., NeoC CASI: AMS ΔR^2^=0.300 versus Module post-IP% ΔR^2^=0.229). In HKG8, the housekeeping gene scores and CS were near zero and not significant, while CP, the same Micro-PVM_3 abundance as in M4, remained significant on most outcomes. Thus, scores over the same nine Module genes recovered much of the cognitive association, so the Module gene panel itself carried most of the signal, whichever method was used to probe the clinical associations.

**Figure 10.**
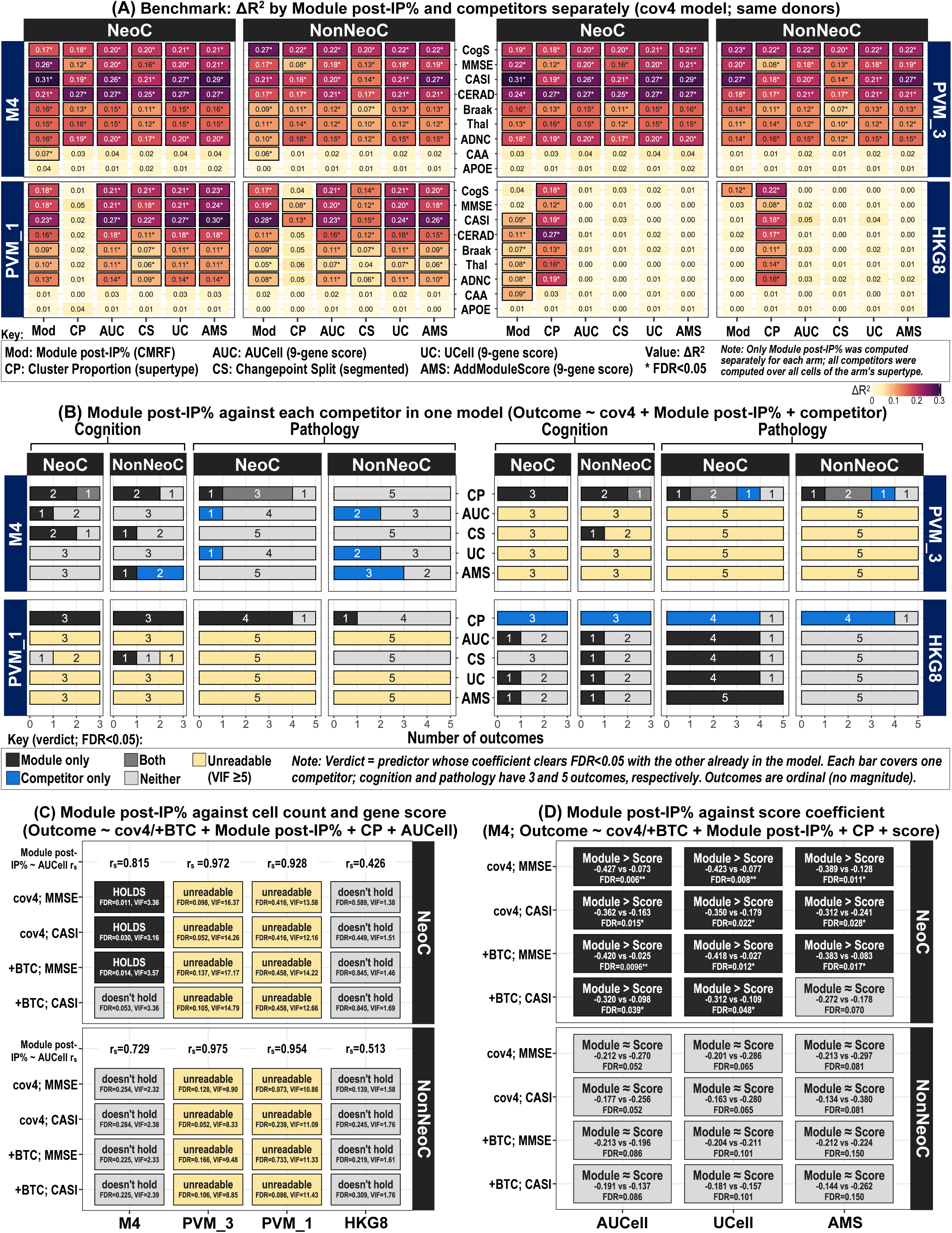
Module post-IP% against five competitors. (A) ΔR^2^ over cov4 of Module post-IP% (Mod) and each competitor fitted alone on the same donors: cluster proportion (CP), the supertype’s cells as a percentage of all Micro-PVM cells; the donor means of AUCell (AUC), UCell (UC), and AddModuleScore (AMS) over the GOI and its eight partners; and changepoint split (CS), the percentage of cells above a segmented-regression breakpoint. Outcomes: cognitive status (CogS), MMSE, CASI, CERAD, Braak, Thal, ADNC, CAA, and *APOE* ε4 dose (APOE). * FDR<0.05 (BH over 15 outcomes per predictor). As competitors use all cells of the supertype, M4 and PVM_3 (both Micro-PVM_3) have identical competitor values, and HKG8 shares only CP with them, as its gene-based competitors use *ACTB* and its eight partners. (B) Outcomes by verdict with Module post-IP% and one competitor in one cov4 model: Module only, competitor only, Both, Neither, or unreadable (joint VIF≥5). (C) Nested F-test of Module post-IP% added to cov4 or +BTC with CP and AUCell, on MMSE and CASI. “HOLDS” indicates FDR<0.05 with the expected sign and all VIFs<5; unreadable indicates VIF≥5; each box shows the FDR and the largest VIF; r_s_ is the correlation of Module post-IP% with AUCell. (D) M4: β of Module post-IP% and of each score in the model with CP; the FDR and label (“Module > Score” at FDR<0.05, otherwise “Module ≈ Score”) refer to the model without CP, hence verdicts can differ from C. * FDR<0.05, ** FDR<0.01. Values are shown in Supplementary Table S12.

Next, Module post-IP% and one competitor were fitted in the same cov4 model to examine which of the two still carried a significant coefficient (FDR<0.05) with the other present: Module only, competitor only, Both, Neither, or unreadable (VIF≥5) (Figure 10B). Against CP (cell abundance), Module post-IP% remained significant but CP did not for most cognitive outcomes in the three main arms: PVM_3 (Module only 5 of 6, including MMSE and CASI in both region sets), PVM_1 (6 of 6), and M4 (4 of 6). In PVM_3, Module post-IP% was also the only significant predictor for Braak in both region sets. Against the gene scores (AUC, UC, and AMS) and CS, the comparison was mostly unreadable in PVM_3 and PVM_1, where Module post-IP% and each gene score or CS were nearly the same quantity per donor (r_s_=0.885-0.978). In M4, where the comparisons were readable (VIF=1.8-3.3), Module post-IP% was the only significant predictor for MMSE in NeoC against AUC or CS, for CASI in NeoC against CS, and for cognitive status in NonNeoC against AMS or CS, whereas AMS was the only significant predictor for MMSE and CASI in NonNeoC. For pathology in M4, no comparison with the gene scores or CS was Module only, and the competitor was the only significant predictor in 9 of 40 tests. In HKG8, CP was the only significant predictor for all six cognitive outcomes.

Collectively, Module post-IP% carried information that cell abundance did not in all three main arms, including MMSE and CASI in both region sets in PVM_3. Against a gene score of the same nine genes, Module post-IP% showed no consistent gain: the two could not be separated in PVM_3 and PVM_1, and in M4, Module post-IP% edged ahead on cognition in NeoC while the gene scores outperformed on pathology staging. MMSE and CASI were therefore the outcomes on which Module post-IP% was tested further.

Module post-IP% was assessed alongside CP and AUC in the same cov4 or +BTC model for MMSE and CASI outcomes, to test whether Module post-IP% was more than a cell count (CP) and a gene score (AUC) combined. In PVM_3 and PVM_1, this three-predictor joint model was unreadable in both region sets (VIF=8.3-17.2), as Module post-IP% and AUC were nearly the same quantity per donor (Figure 10C). In M4 NeoC, where the model was readable, Module post-IP% remained predictive of MMSE (FDR=0.011) and CASI (FDR=0.030) at cov4, and of MMSE (FDR=0.014) but not CASI (FDR=0.053) at +BTC; the CASI coefficient pointed in the same direction, and fewer donors had CASI (69 against 80 for MMSE). None held in M4 NonNeoC (FDR=0.225-0.284) or in HKG8 (FDR=0.139-0.845).

As the joint models were unreadable in PVM_3 and PVM_1, Module post-IP% was then tested alongside CP with each of the three gene scores in turn in M4 only (Figure 10D). In M4 NeoC, Module post-IP% remained significantly associated with MMSE at both cov4 and +BTC against AUC, UC, or AMS (FDR≤0.017), and with CASI at cov4 and +BTC against AUC or UC (FDR≤0.048) and against AMS at cov4 (FDR=0.028) but not at +BTC (FDR=0.070). In M4 NonNeoC, Module post-IP% was not significant in any comparison (FDR=0.052-0.150). These FDRs are from the model without CP (Figure 10D). With CP in the model, as in Figure 10C, the CASI association in M4 NeoC held only at cov4 against AUC (FDR=0.030) and UC (FDR=0.039), and the NonNeoC FDRs were 0.209-0.381. Taken together, in M4 NeoC, the MMSE association of Module post-IP% held against all three gene scores, and the CASI association held against all except AMS at +BTC in the model without CP; AMS was the strongest competitor.

### 3.8 Ablation of CMRF-derived boundaries of Module post-IP%

In this ablation analysis, three simpler boundaries were built in the same copula plane as CMRF, i.e., OLS (a straight line with a constant half-width), fixed spline (a curve with a constant half-width), and local spline (the same curve, but with a half-width that varies along u), each set to hold about the same number of cells as the CMRF Module.

In the three main arms, all four boundaries yielded significance in all 48 tests at cov4 and at +BTC (all FDR<0.05), with standardized β=-0.462 to -0.273 at +BTC (Figure 11A; Supplementary Table S12), whereas the control arm HKG8 yielded no significance in any of its 16 tests at +BTC (FDR≥0.254). CMRF had the most negative β in 7 of the 12 arm, region set, and outcome combinations, but the differences between boundaries were small. Hence, the cognition association of Module post-IP% survived pathology adjustment however the boundary was drawn, as long as it followed the *MT-CO3*/cDFG relationship and held about the same number of cells. Figure 11B shows the four boundaries on the M4 NeoC cells.

**Figure 11.**
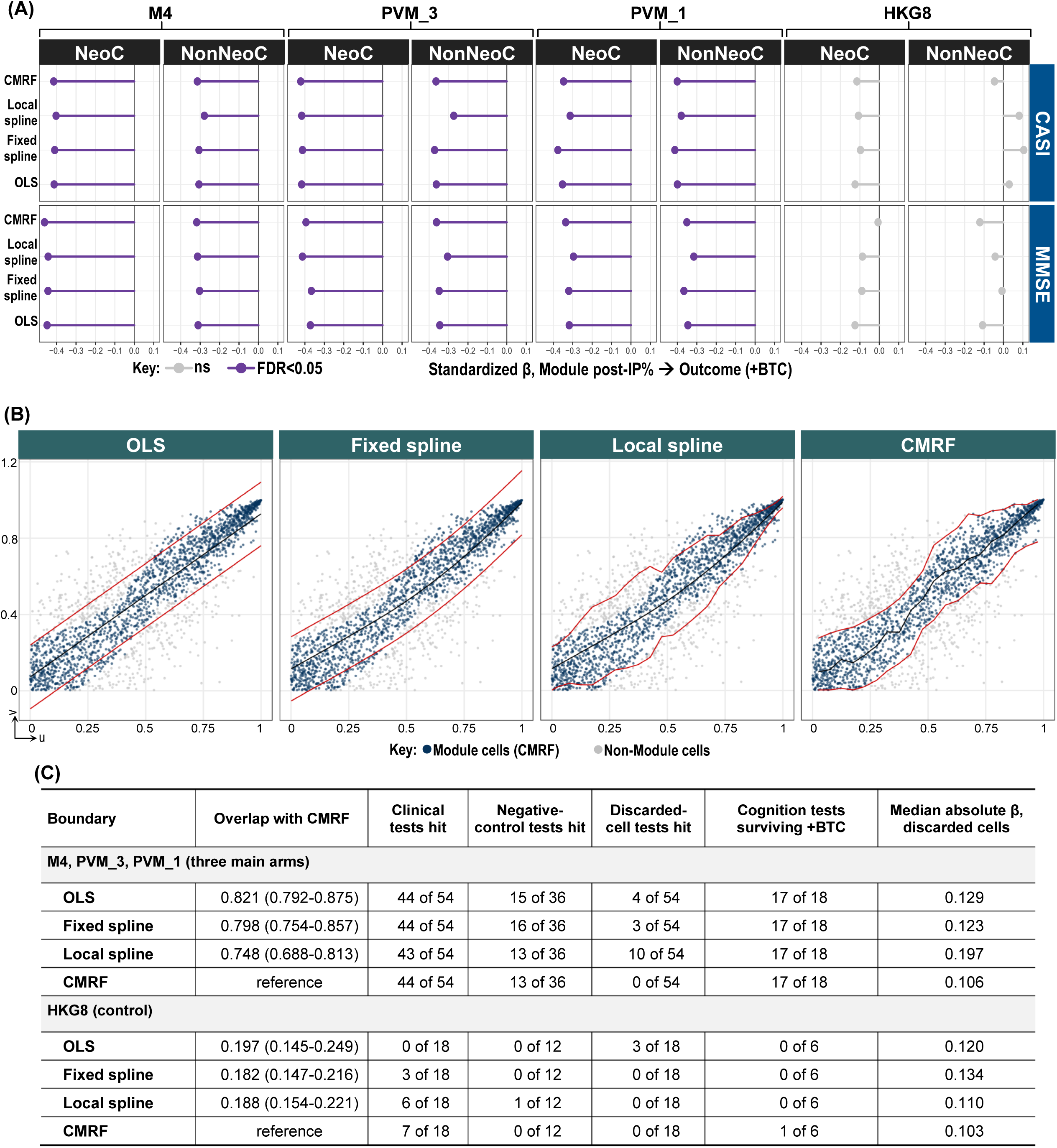
Boundary ablation: CMRF against three simpler boundaries. Each boundary was built in the copula plane (u, v) of each arm and region set to hold about as many cells as the CMRF Module (within 19); cells outside it were discarded and split at the IP as in Figure 9. (A) Standardized β of each boundary’s Module post-IP% on CASI and MMSE at +BTC; violet, FDR<0.05 (BH over 128 tests). (B) The four boundaries on M4 NeoC (n=2,130): OLS line and fixed (penalized) spline, each with a constant half-width; local spline, the same spline with a half-width estimated in 20 equal-count bins of u; and CMRF, the smallest and largest v of Module cells in the same bins, with the median as center line. Red, boundary lines; black, center line; colors, CMRF Module membership. (C) Scorecard for the main arms and HKG8: median (range) Jaccard overlap with the CMRF Module; hits among clinical tests (nine outcomes, cov4), negative-control tests (six outcomes), discarded-cell tests (the discarded cells’ coefficient with both donor percentages in the model), and cognition tests surviving +BTC; and the median |β| of the discarded cells. A hit is a test with FDR<0.05 (BH over the outcomes of each model; Supplementary Table S12).

Figure 11C compares the four boundaries on the same tests. In the main arms, the simpler boundaries shared most cells with the CMRF Module (median Jaccard 0.748-0.821) and yielded nearly the same results: 43-44 of 54 clinical associations were significant at cov4, and 17 of 18 cognition tests were still significant after +BTC. They differed in the cells that were left out. CMRF was the only boundary whose discarded cells carried no clinical association (0 of 54); OLS, fixed spline, and local spline left out signal-bearing cells (in 4, 3, and 10 of 54 tests, respectively). Negative-control hits were similar for every boundary (13-16 of 36), and in HKG8, clinical hits were low in all four boundaries (0-7 of 18). The CMRF boundary therefore did not increase the clinical association, but CMRF provided a cleaner split, leaving out cells with no clinical signal, while every simpler boundary left out some cells that still carried it.

### 3.9 Battery 1: Contamination and doublets did not account for the cognition associations

Battery 1 tested whether Module post-IP% results were confounded by contamination. Of the 48 of 72 models significant before adjustment (FDR<0.05), 35 remained significant after adding donor-mean DecontX contamination (donors with at least five non-Module cells) (Figure 12A; Supplementary Table S13). Battery 1 used the donors with labeled cells of each arm, so M4 and HKG8 had fewer donors than in Figure 9A (M4: 78 and 80; HKG8: 74 and 77), and 48 associations were significant against 51 in Figure 9A.

**Figure 12.**
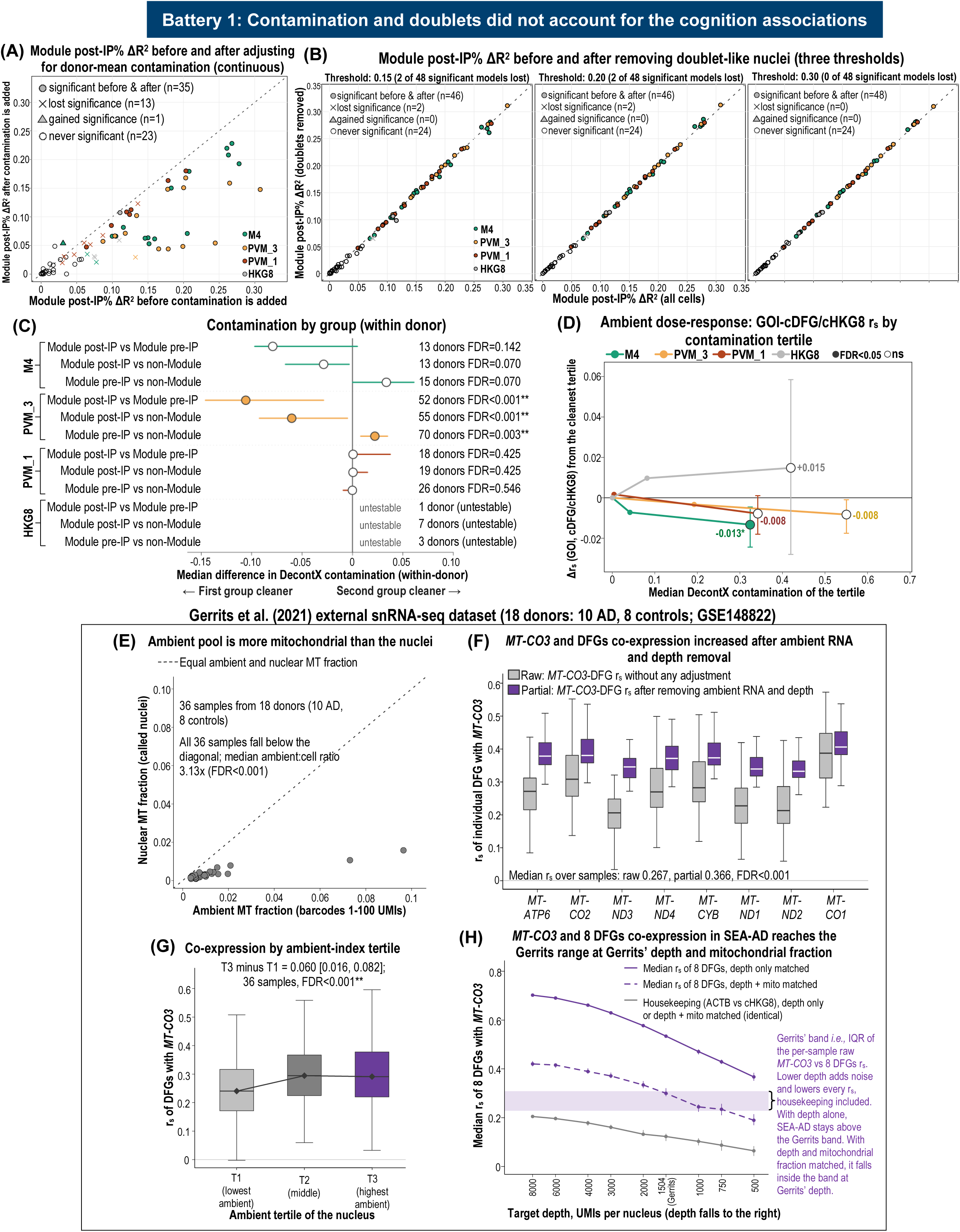
Battery 1. Filled symbols and * mark FDR<0.05; ** marks FDR<0.01. (A) Contamination as a covariate. Module post-IP% ΔR² before (x-axis) and after (y-axis) adding donor-mean DecontX contamination (72 cov4 models), using all donors with labeled cells before and those with a contamination value after. Dashed line: equal ΔR². (B) Doublet removal. Module post-IP% ΔR² before and after removing Module post-IP nuclei with a doublet score ≥0.15, 0.20, or 0.30. (C) Contamination by group. Donor median of the within-donor difference in median DecontX contamination, with donor-bootstrap 95% CIs; negative: the first group was cleaner; untestable: fewer than 10 donors. (D) Co-expression by contamination tertile. Change in r_s_ of the GOI with the composite in Module cells from the cleanest tertile; bars: 95% CIs of most minus least contaminated tertile. (E) Ambient against nuclear mitochondrial fraction, for barcodes with 1-100 UMIs (x-axis) and called nuclei (y-axis). (F) Raw and partial co-expression. r_s_ of MT-CO3 with each DFG per sample, raw r_s_ (gray) and partial r_s_ after removing the ambient index and log depth (purple). (G) Co-expression by ambient-index tertile. Raw r_s_ of MT-CO3 with each DFG per tertile and sample. (H) Depth and mitochondrial thinning. PVM_3 nuclei thinned to each target depth alone (solid) or also to the Gerrits median mitochondrial fraction (dashed); points: median r_s_ of MT-CO3 with the eight DFGs; shaded: Gerrits per-sample IQR; gray: housekeeping control.

Seven of the 13 losses were in PVM_1 NonNeoC, where 20 of 83 donors had no contamination value (66-95% of ΔR^2^ retained on the other 63 donors) and were included only in the model before adjustment, which the pre-specified reading classed as borderline because ΔR^2^ fell by less than half. All six PVM_3 cognition models remained significant (FDR=2.82×10^-4^ to 0.043 after adjustment), with the lowest retention in NeoC cognitive status (ΔR^2^ from 0.199 to 0.048, n=83). All six M4 cognition models also remained significant, with the largest decreases in NeoC (cognitive status: ΔR^2^ from 0.150 to 0.063, FDR from 6.09×10^-4^ to 0.022, n=78; MMSE: ΔR^2^ from 0.263 to 0.208; CASI: ΔR^2^ from 0.278 to 0.193).

In addition, removing Module post-IP nuclei with a doublet score at or above 0.15, 0.20, or 0.30 lost two of the 48 significant models at 0.15 and 0.20 (HKG8 NeoC CERAD and Braak) and none at 0.30; all six PVM_3 and all six M4 cognition models remained significant at every threshold (Figure 12B).

If Module post-IP cells were an ambient artifact, they would be the most contaminated. Within donor, no arm showed significantly more contamination in Module post-IP than in Module pre-IP cells (Figure 12C). In PVM_3, Module post-IP cells were cleaner than Module pre-IP (median difference: -0.106, 52 donors, FDR≤3.00×10^-4^) and non-Module cells (median difference: -0.060, 55 donors, FDR≤3.00×10^-4^); in M4 they were cleaner without reaching significance (median difference: -0.079, 13 donors, FDR=0.142). Likewise, no arm showed a significant rise of co-expression with DecontX tertile (Figure 12D), so higher contamination did not raise the co-expression.

In the external Gerrits dataset (18 donors: 10 AD and 8 controls), the ambient pool was richer in mitochondrial RNA than the called nuclei in all 36 samples (3.13-fold; median log ratio: 1.14, FDR≤2.00×10^-4^; Figure 12E). Removing the ambient index and log depth increased the r_s_ of *MT-CO3* with its eight DFGs (raw r_s_=0.267, partial r_s_=0.366, both FDR≤2.00×10^-4^; Figure 12F).

Co-expression was also stronger in nuclei with more ambient RNA (T3 minus T1 ambient-index tertile Δr_s_=0.060, FDR≤2.00×10^-4^; Figure 12G). By the pre-specified reading, ambient RNA thus contributed to the co-expression in the Gerrits dataset, which persisted once the ambient index and depth were removed. The raw co-expression was higher in the SEA-AD PVM_3 arm (median r_s_ of *MT-CO3* with each of its eight DFGs=0.711) than in Gerrits (r_s_=0.267); matching both depth and mitochondrial fraction lowered SEA-AD r_s_ to 0.300, inside the Gerrits IQR (0.228-0.309; Figure 12H), thus depth and mitochondrial content together accounted for the difference.

Within donor, Module post-IP cells had higher own-ER and ribosomal shares and a lower nuclear-retained *NEAT1* share than Module pre-IP cells in PVM_3 and PVM_1 (FDR<0.05), and a higher other-lineage index in PVM_3 only (0.017 pp, FDR≤4.00×10^-4^; Supplementary Figure S15). M4 showed no significant shift, and HKG8 was untestable.

### 3.10 Battery 2: Module post-IP% behaved as a donor-level property

Battery 2 tested whether Module post-IP% was a donor-level property, that is, whether donors with high Module post-IP% in some brain regions also tended to have high values in their other regions (of the 10 regions, e.g., MTG or LEC). Each region was dissected and sequenced as a separate sample. Technical errors (e.g., tissue sampling, library preparation, or sequencing) affect only that sample and cannot create this donor-level pattern. Additionally, donor-wide confounders that can affect all regions alike, such as PMI and RIN, were adjusted for by TQR.

In M4, PVM_3, and PVM_1, the donor share of variance of Module post-IP% over units was 69.6%, 73.2%, and 73.9%, exceeding the 95th percentile of the within-region shuffle null by 21.8, 51.0, and 33.9 pp (all FDR<0.01; Figure 13A; Supplementary Table S14), and was 58.3%, 66.7%, and 66.1% after TQR (all FDR<0.01). Hence, Module post-IP% repeated between a donor’s brain regions beyond chance, also when adjusted for tissue quality. In HKG8, Module post-IP% did not exceed its null (donor share 41.7%, null 95th percentile 56.4%, FDR=0.520), whereas %MT and the matched non-mitochondrial share exceeded their nulls on all four arms.

**Figure 13.**
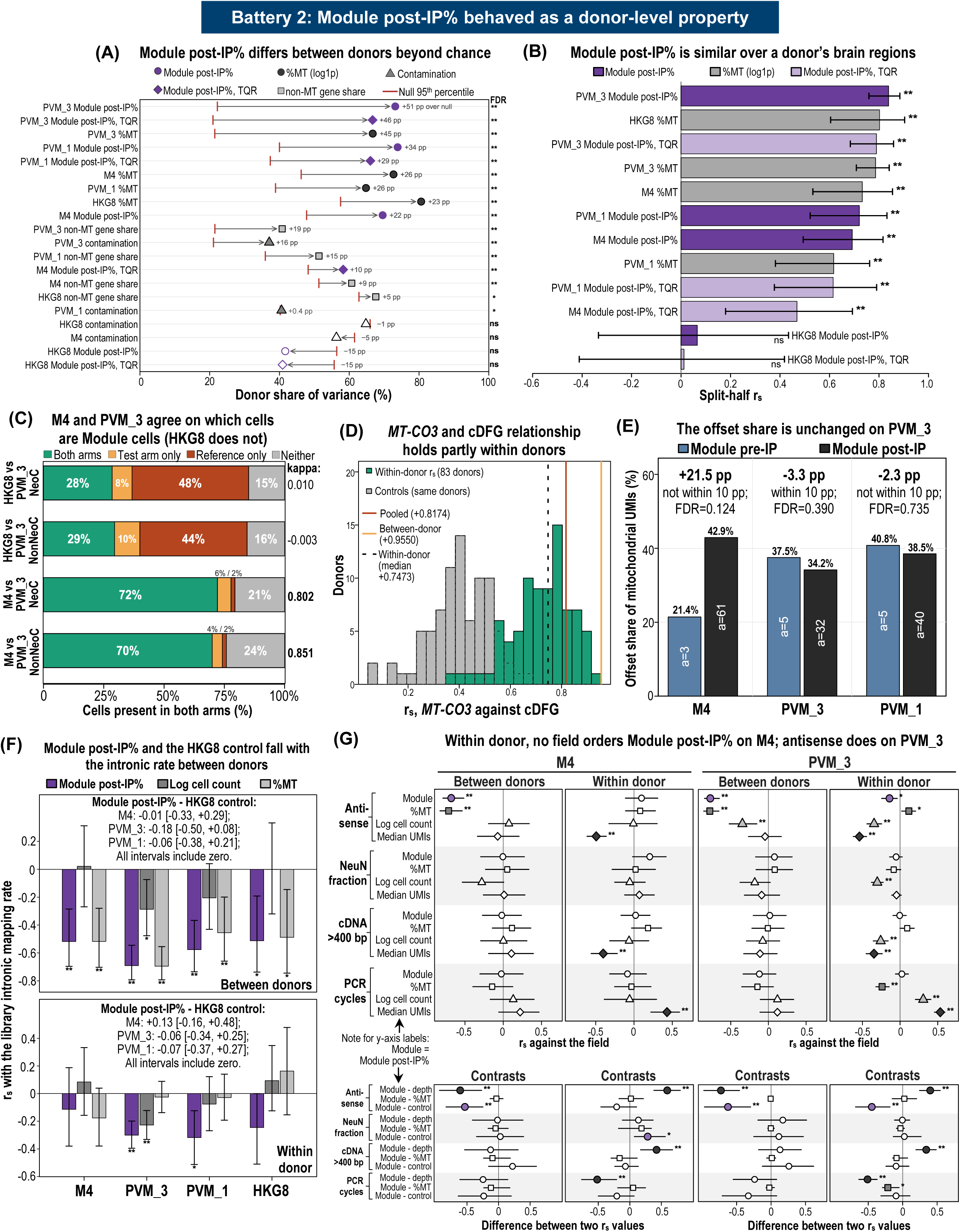
Battery 2. Filled symbols and * mark FDR<0.05; ** marks FDR<0.01. (A) Donor share of variance of Module post-IP%, its residual after TQR, %MT, DecontX contamination, and a matched non-mitochondrial share. Red tick: 95th percentile of within-region shuffles. (B) Split-half repeat. r_s_ over donors between random halves of each donor’s units, median over 200 splits. (C) CMRF label agreement on cells shared by a test arm (M4 or HKG8) and the PVM_3 reference. Both arms: a Module cell in both; Test arm only and Reference only: a Module cell in one arm; Neither: a non-Module cell in both. (D) Within-donor co-expression in PVM_3 cells. r_s_ of MT-CO3 with cDFG per donor, and of ACTB with cHKG8 (control; gray). Dashed line: median within-donor r_s_; red: r_s_ pooled over donors; orange: r_s_ between donor means. (E) Offset share of Module pre-IP and post-IP cells. Above each arm, the post-IP minus pre-IP difference (pp), whether its 90% CI is within the 10 pp margin, and the non-zero test FDR. (F) Library intronic rate. r_s_ of Module post-IP%, log cell count, and %MT with the intronic mapping rate, between donors (top) and within donor (bottom). Above: each arm’s Module post-IP% r_s_ minus that of HKG8. (G) Library fields in M4 and PVM_3. Top: r_s_ of each quantity with the field. Bottom: contrasts of the Module post-IP% r_s_ minus the r_s_ of depth, %MT, or the HKG8 control for the same field, with 95% CIs from the same donor resamples.

Split-half r_s_ (whether donors with high Module post-IP% in one random half of their regions also had high values in the other half; higher values mean more consistency within donors) was 0.692 in M4, 0.839 in PVM_3, and 0.721 in PVM_1 (all FDR<0.01), and 0.469, 0.790, and 0.615 after TQR (all FDR<0.01; Figure 13B). In HKG8, it was 0.066 (FDR=0.970).

M4, built on the same mitochondrial genes as PVM_3, assigned mostly the same cells to the Module (Cohen’s κ=0.802 in NeoC and 0.851 in NonNeoC), above the pre-specified 0.60; HKG8 did not (κ=0.010 and -0.003), inside the pre-specified |κ|<0.20 (Figure 13C). Thus, with housekeeping genes, the same CMRF method selected a different set of cells, ruling out an artifact of the CMRF method that would select the same cells from any genes.

Pooled correlations can arise from donor differences alone. In the PVM_3 arm cells, the median within-donor r_s_ of *MT-CO3* with cDFG was 0.747 (positive in every donor; 0.743 with log depth partialed out) against 0.817 with all cells pooled and 0.955 between donor means (Figure 13D), and it exceeded the control pair (*ACTB* with cHKG8; median r_s_=0.402) in all 83 donors. Thus, the *MT-CO3* relationship with cDFG did not arise from donor differences alone.

The intercept of mitochondrial on total UMIs is a fixed number of mitochondrial UMIs that every nucleus carries regardless of sequencing depth, the pattern expected of background (ambient) mitochondrial RNA. In PVM_3, the offset share was 34.2% in Module post-IP and 37.5% in Module pre-IP cells (difference -3.3 pp, 90% CI: -9.2 to 1.8, FDR=0.390), within the pre-specified 10 pp margin (Figure 13E). The difference was not significant after FDR correction in M4 (+21.5 pp, 90% CI: 4.5-35.0, FDR=0.124) and PVM_1 (-2.3 pp, 90% CI: -17.1 to 13.6, FDR=0.735), but neither was within the 10 pp margin, while HKG8 was untestable.

Next, it was assessed whether Module post-IP% tracked the intronic mapping rate, where a lower intronic rate indicates a library with relatively more cytoplasmic, non-nuclear RNA. Between donors, Module post-IP% fell with the library intronic rate in M4, PVM_3, and PVM_1 (r_s_=-0.518, -0.690, and -0.577, respectively; all FDR<0.001) and in HKG8 (r_s_=-0.512, FDR=0.013). Within donor, it fell in PVM_3 (r_s_=-0.301, FDR<0.001) and PVM_1 (r_s_=-0.318, FDR=0.010), but not in M4 (r_s_=-0.114, FDR=0.562; Figure 13F). No arm differed from HKG8 (all FDR≥0.299), so the association was not specific to the Module.

Finally, whether Module post-IP% followed a technical property of the sequencing library was investigated, including the antisense rate, the NeuN-positive fraction, cDNA >400 bp, and PCR cycles. Within donor, a technical field ordered Module post-IP% only if its r_s_ with Module post-IP% had FDR<0.05 and differed from its r_s_ with the HKG8 Module post-IP%. This r_s_ also had to differ from the field’s r_s_ with sequencing depth (log median UMIs) and be at least as strong as its r_s_ with %MT, so that the association was not explained by depth or %MT.

Between donors, Module post-IP% fell with the antisense rate in M4, PVM_3, and PVM_1 (r_s_=-0.680, - 0.772, and -0.683, respectively; all FDR≤0.002), as did %MT, whereas the HKG8 Module did not (r_s_=- 0.147, FDR=0.585; Figure 13G). Within donor, the antisense rate met all four conditions in PVM_3 only (r_s_=-0.141, FDR=0.010), and in M4 it was not significant (r_s_=0.098, FDR=0.869). No other field was significant for Module post-IP% in any arm (Supplementary Figure S16). Thus, within donor, only the antisense rate ordered Module post-IP%, and only in PVM_3; no library field did so in M4 or PVM_1.

### 3.11 Battery 3: Module post-IP% shared histology with %MT, and predicted cognition beyond %MT in M4

Module post-IP% and %MT could be separated only in M4, the smallest arm. Battery 3 tested whether Module post-IP% tracked microglial activation, measured on stained tissue sections independently of the sequencing data, and whether it carried information beyond %MT. Within donor in PVM_3 (69 donors), Module post-IP% correlated in the pre-specified direction with all four IBA1 activation measures (r_s_=- 0.240 to 0.167, all FDR<0.05; Figure 14A; Supplementary Table S15), and %MT tracked them with the same signs (all FDR<0.01). The cytoplasmic share (own-ER plus ribosomal genes as a percentage of non-mitochondrial UMIs) fell with activated IBA1^+^ cells per area (r_s_=-0.153, FDR=0.008), opposite in sign to Module post-IP%. IBA1^+^ area fell with Module post-IP% in PVM_3 (r_s_=-0.229, FDR=0.001) and also in the HKG8 control (r_s_=-0.426, FDR=0.019), thus this association was not specific to the Module. In M4 (19 donors) and PVM_1 (22 donors), no IBA1 measure reached FDR<0.05 with Module post-IP%. In PVM_3, the association with activation was shared with %MT.

**Figure 14.**
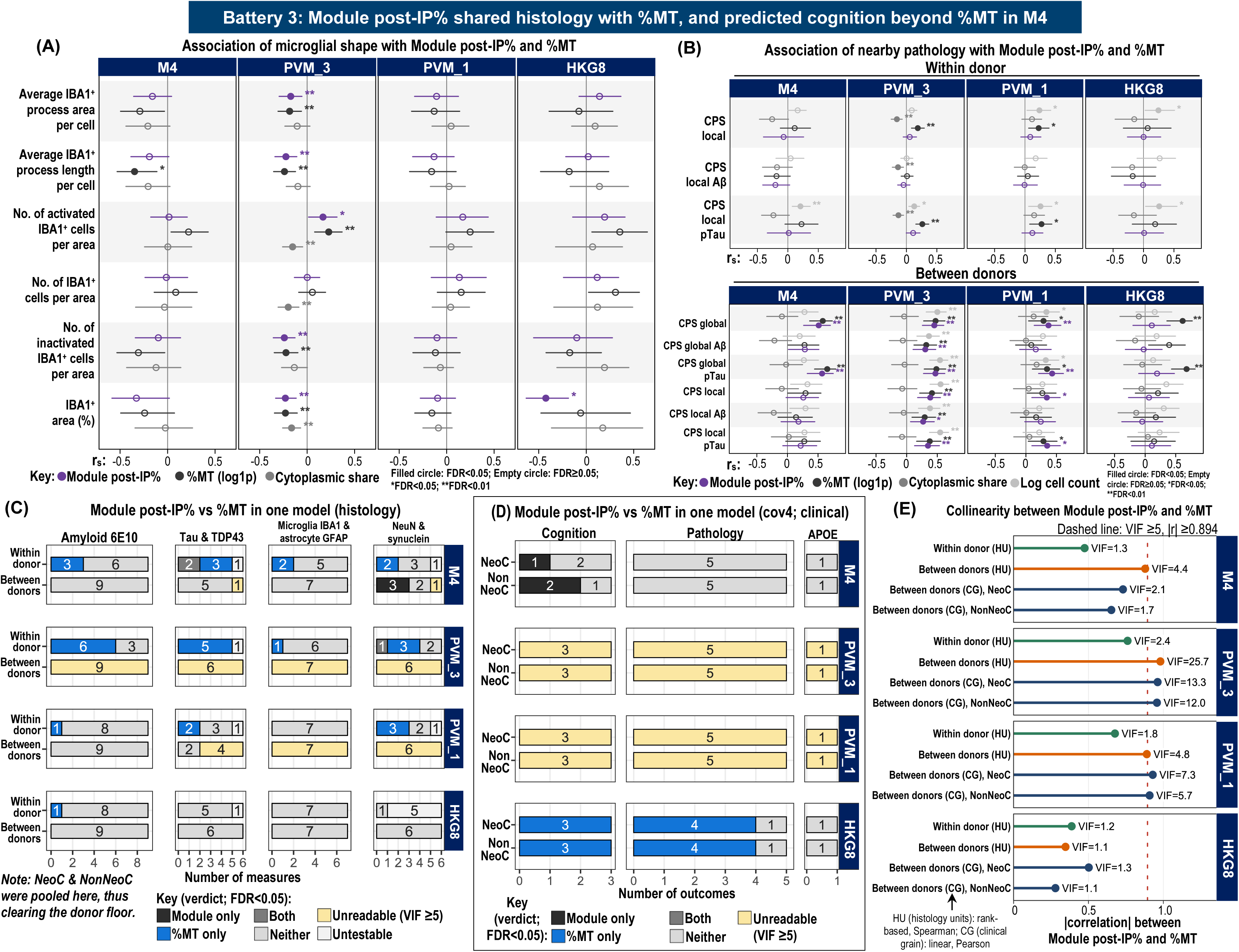
Battery 3. Units were joined to the SEA-AD neuropathology and CPS values of the same donor and region, from adjacent sections; A to C pool NeoC and NonNeoC. Intervals: donor-bootstrap 95% CIs. Filled symbols and * mark FDR<0.05; ** marks FDR<0.01. (A) Microglial shape. Within-donor r_s_ of six IBA1 measures with Module post-IP% (purple), %MT (log1p; black), and the cytoplasmic share (gray). (B) Nearby pathology. r_s_ of the overall, amyloid-beta (Aβ), and pTau CPS with the quantities of A and log cell count; upper: local scores within donor; lower: local and global scores between donors. (C) Histology, one model. Ranked stain measure regressed on ranked Module post-IP% and ranked %MT without covariates, within and between donors, for 28 measures. Bars count measures per verdict by stain family: Module only, %MT only, Both, or Neither at FDR<0.05; Untestable indicates not fitted; Unreadable indicates VIF≥5. (D) Each outcome regressed by OLS on cov4, Module post-IP%, and %MT; verdicts as in C, for cognition, pathology, and APOE ε4 dose. (E) |r| between Module post-IP% and %MT in each model frame: histology within donor (green) and between donors (orange), Spearman; clinical (blue), Pearson, per region set. Dashed line, VIF=5 (|r|=0.894).

Within donor, Module post-IP% tracked no local CPS in any arm (all FDR≥0.165; Figure 14B), whereas %MT rose with local CPS and local pTau CPS in PVM_3 and PVM_1 (r_s_=0.187-0.272, all FDR≤0.013). Between donors, Module post-IP% rose with global CPS and global pTau CPS in M4 (r_s_=0.511 and 0.566), with all six scores in PVM_3 (r_s_=0.269-0.470), with four in PVM_1 (r_s_=0.342-0.429; all FDR<0.05), and with none in HKG8. According to the pre-specified reading, it was a donor-level marker of disease stage in the three main arms; %MT showed similar between-donor associations.

When Module post-IP% and %MT were tested in the same model for each of the 28 stain measures within donor (Figure 14C), no measure in any arm was Module only. Within donor, %MT only was the most common verdict with a significant predictor, with 10 measures in M4, 15 in PVM_3, 6 in PVM_1, and 1 in HKG8. Module post-IP% therefore related to these measures only through the part it shared with %MT, apart from three Both verdicts (two pTDP43 measures in M4 and NeuN⁺ cells per area in PVM_3). Between donors, the two predictors were too collinear to separate (VIF≥5) for all 28 measures in PVM_3, 17 in PVM_1, and 2 in M4. In M4, the three synuclein measures were Module only (Module coefficient: 0.767-0.819, all FDR<0.05; 50 donors; VIF=4.39).

Summarized by stain family within donor (Supplementary Figure S17A), Module post-IP% tracked no family in M4 or HKG8. In PVM_3, it tracked the tau, TDP43, GFAP, and IBA1 families (median r_s_=-0.198 to 0.232, all FDR<0.05), and %MT tracked the same four with larger medians as well as amyloid 6E10 and synuclein. In PVM_1, Module post-IP% tracked only TDP43. The family median of |Module r_s_|-|%MT r_s_| was below zero at FDR<0.05 for tau, TDP43, and NeuN in PVM_3, and above zero for no family in any arm (Supplementary Figure S17B).

In contrast, the two predictors were separable on the clinical outcomes in M4 in both region sets (VIF≤2.39), and the verdict was Module only for MMSE in NeoC (β=-0.529, FDR=0.003; 74 donors) and for cognitive status (β=0.415, FDR=0.015; 80 donors) and MMSE (β=-0.337, FDR=0.028; 76 donors) in NonNeoC (Figure 14D). The remaining M4 outcomes (8 in NeoC and 7 in NonNeoC), including all pathology scores and APOE ε4 dose, were Neither, and the %MT coefficient was insignificant for all M4 outcomes (FDR≥0.094). All 18 comparisons on each of PVM_3 and PVM_1 were unreadable (VIF≥5). In HKG8, 14 of 18 comparisons were %MT only.

Between donors, the models with Module post-IP% and %MT entered together were unreadable in all 46 models of PVM_3 and in 35 of 46 of PVM_1 (VIF=11.7-25.7 and 5.5-10.6, respectively; Figure 14E; Supplementary Table S15). In M4 and HKG8, the two predictors were separable in the clinical models (VIF=1.7-2.4 and 1.1-1.3, respectively). Within donor, the VIF was 1.1-3.8, and all fitted models were readable.

### 3.12 Battery 4: Neither nuclear-encoded panels nor per-cell %MT reproduced the Module

Battery 4 assessed whether %MT or a nuclear-encoded mitochondrial gene panel could replace Module post-IP%. Module post-IP% or %MT and each panel score were separable in every model (VIF≤2.01). In PVM_3 and PVM_1, Module post-IP% and %MT each retained every cognition comparison with every panel (Figure 15A; Supplementary Table S16). In M4 NeoC (26-34 donors), Module post-IP% retained its MMSE association against 14 of the 15 panels and the matched reference (a null set of 300 non-MT genes outside all panels), and retained cognitive status and CASI against 4 and 7 panels. When %MT took the place of Module post-IP%, 1 of the 48 M4 NeoC cognition comparisons was %MT only, and the other 47 were Neither. In M4 NonNeoC (23-31 donors), Module post-IP% retained 8 of 48 cognition comparisons (Supplementary Figure S18), and in HKG8, it retained none. According to the pre-specified reading, no nuclear-encoded panel replaced Module post-IP%, as no panel was the only significant predictor of a cognition outcome.

**Figure 15.**
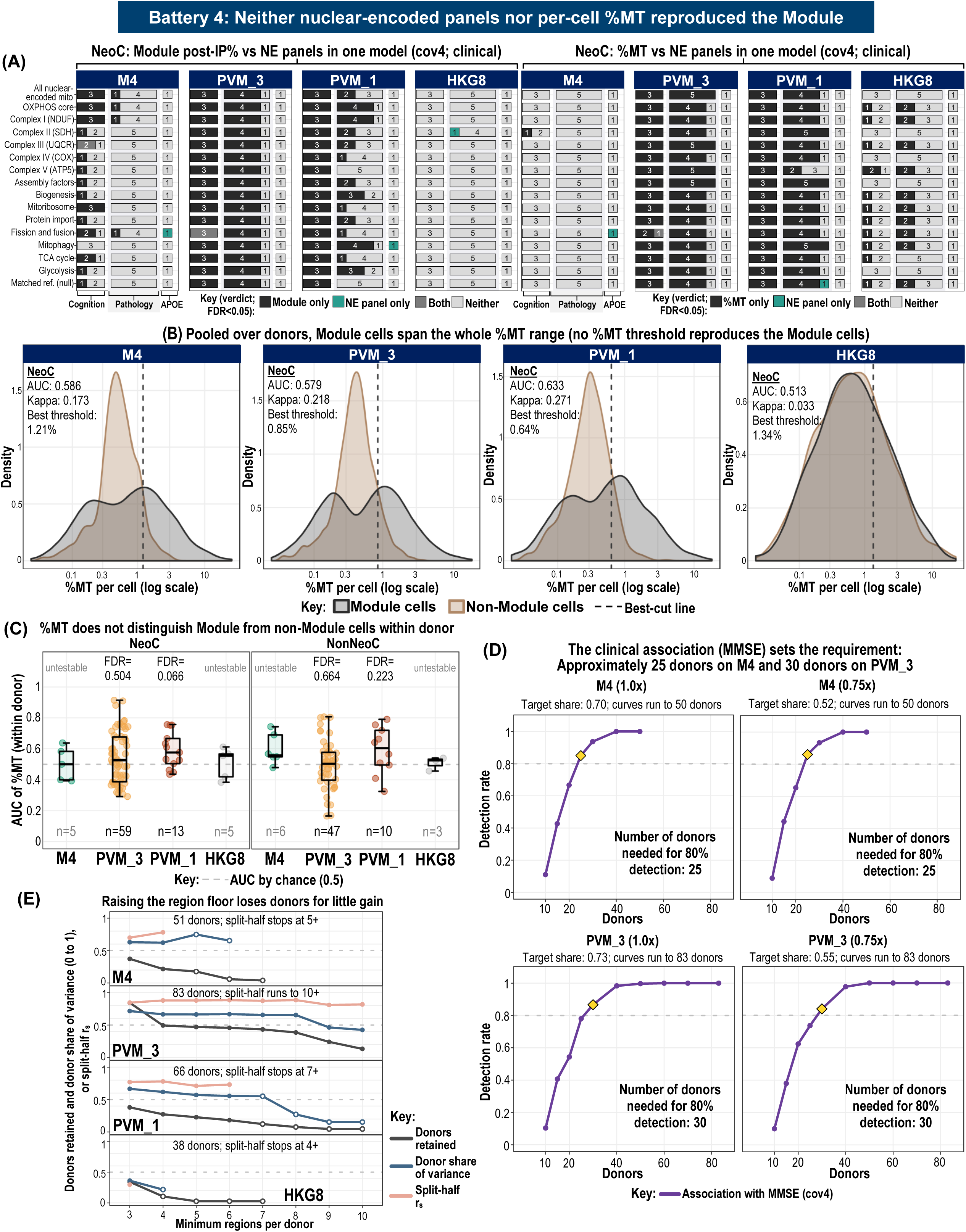
Battery 4. A and B show NeoC (NonNeoC in Supplementary Figures S18 and S19); D and E use the units of Figure 13A. (A) Module post-IP% (left) or %MT (right) with each nuclear-encoded (NE) panel in one cov4 model per outcome, as in Figure 14D: 15 NE panels (14 mitochondrial and one glycolysis) and a matched reference set. Bars count outcomes per verdict for cognition, pathology, and APOE ε4 dose: Module only (or %MT only), NE panel only, Both, or Neither. (B) Kernel densities of per-cell %MT (log scale) in Module and non-Module cells pooled over donors; dashed line: best threshold (Youden’s J); inset: AUC for Module membership and Cohen’s κ at that threshold. (C) Within-donor AUC of %MT for Module membership, in donors with ≥20 cells of each class; boxes: median and quartiles; dashed line: 0.5; untestable: fewer than 10 donors. (D) Detection rate of the MMSE association (cov4, *p*<0.05) against the number of donors subsampled in M4 and PVM_3, with donor share at 1.0x or 0.75x the observed; diamonds: first count reaching 80%. Other rules, arms, and 0.5x are in Supplementary Table S16. (E) Fraction of donors retained, donor share, and split-half r_s_ against the minimum regions per donor. Open points, fewer than 10 donors; dashed line, 0.50 split-half rule.

Pooled over donors, per-cell %MT did not recover the Module cells in M4 (AUC=0.547-0.586) or PVM_3 (0.559-0.579), and the best thresholds (%MT of 0.85-1.30%) agreed with the Module labels at κ=0.173-0.218 (Figure 15B; Supplementary Figure S19). In PVM_1, %MT recovered them only in part (AUC=0.615-0.633, κ=0.242-0.271), and in HKG8 not at all (AUC=0.513-0.520, κ≤0.044). Within donor, the median AUC did not differ from 0.5 in PVM_3 or PVM_1 (all FDR≥0.066); M4 and HKG8 were untestable, with only 3 to 6 donors (Figure 15C).

In donor subsampling, the MMSE association was detected in at least 80% of draws at 30 donors in PVM_3, 25 in M4, and 40 in PVM_1 (Figure 15D). At 0.75x the observed donor share, PVM_3 also met the donor share and split-half rules at 10 and 15 donors. In M4, the donor share rule needed 50 donors and the split-half rule never reached 80%. In HKG8, no rule reached 80% at any donor count or effect.

Sampling more brain regions per donor did not make Module post-IP% a stronger donor property in PVM_3 (Figure 15E). A region was counted for a donor only when it had at least 10 cells of the arm. In PVM_3, 70 of the 83 donors had three or more such regions, and 41 had four or more. Keeping only the 41 donors lowered the share of variation in Module post-IP% explained by donors (among the 70 donors with three or more regions) from 71.6% to 66.5%, and the split-half r_s_ changed little, from 0.846 to 0.880. Hence, three regions per donor were sufficient in PVM_3. The other arms had fewer cells per region, hence fewer donors had three or more regions: 19 of 51 in M4, 25 of 66 in PVM_1, and 13 of 38 in HKG8.

### 3.13 Battery 5: NeoC MMSE held in PVM_3 and M4 without the 11 SA donors

Battery 5 tested whether the 11 SA donors drove the results of Batteries 1 to 4. Without them, all 44 unadjusted and all 35 contamination-adjusted clinical associations kept their direction, and the Module post-IP% coefficients fell to a median of 74% and 71% of their all-donor values, respectively (Figure 16C; Supplementary Table S17).

**Figure 16.**
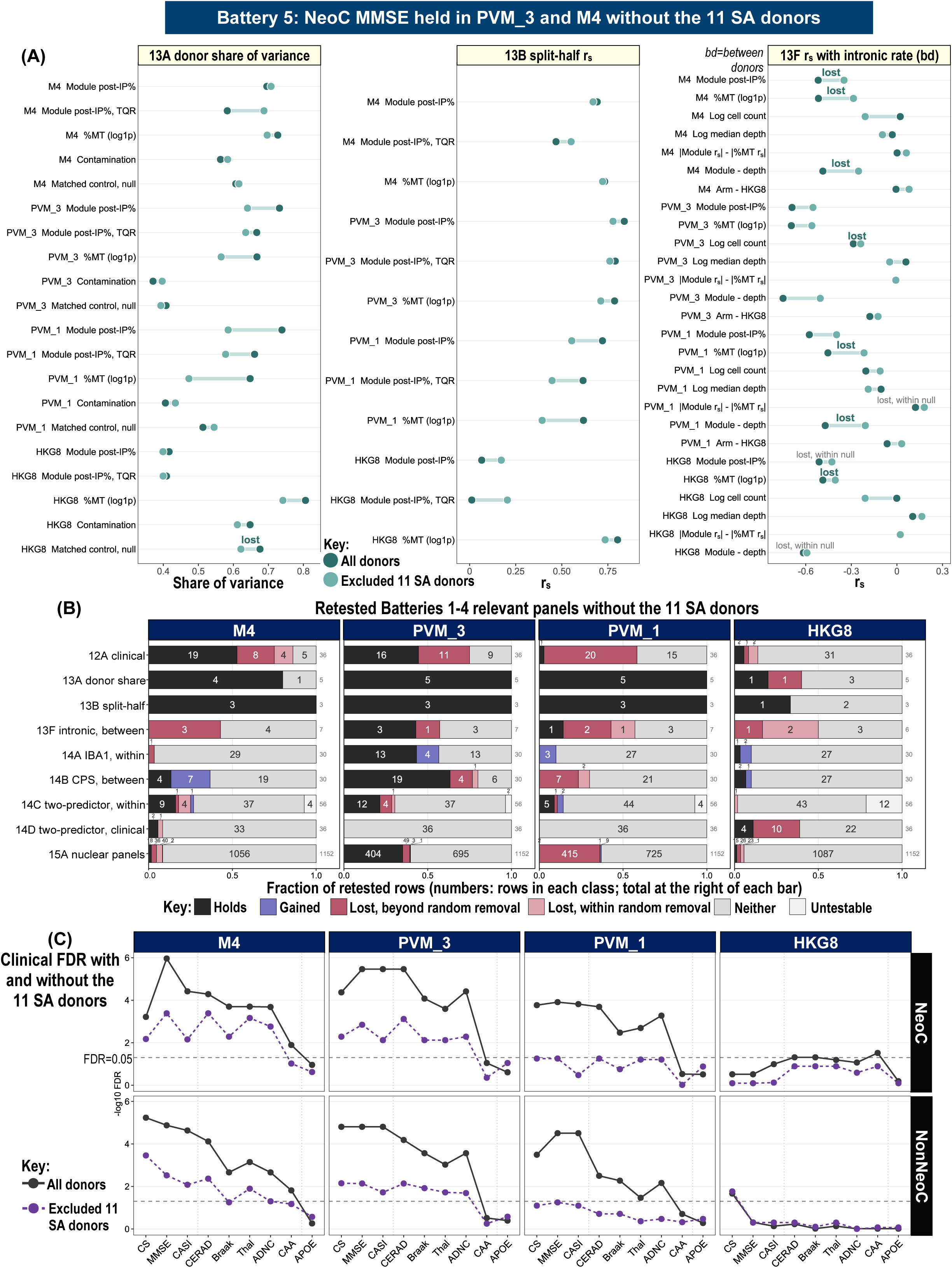
Battery 5. The SA donors, all with High ADNC, were removed from the frame of each retested statistic, with cell labels, units, Module post-IP%, %MT, and the 15A panel scores held at their all-donor values; each change was compared with 1,000 random removals of as many non-SA High-ADNC donors from the same frame. (A) Figure 13 statistics with all donors (dark) and without the SA donors (light): 13A donor share of variance (TQR: tissue quality removed; matched control: the null of the matched non-mitochondrial share), 13B split-half r_s_, and 13F between-donor r_s_ with the library intronic rate and its contrasts. Lost: significant with all donors and not without, with a move toward zero larger than in 95% of the random removals; lost, within random removal: a move within that range. (B) Classes of the retested comparisons per panel and arm. 12A: unadjusted and contamination-adjusted coefficients; 13A and 13B: every quantity; 13F and 14B: between donors; 14A and 14C: within donor; 14C, 14D, and 15A: both coefficients. Holds: significant with and without the SA donors with the same sign; Gained: significant only without them; Lost: as in A; Neither: significant in neither run; Untestable: below a floor. (C) FDR (- log_10_) of the Module post-IP% coefficient in the unadjusted 12A model (cov4, no contamination covariate) per outcome, arm, and region set, with all donors (solid) and without the SA donors (dashed); horizontal line: FDR=0.05. CS: cognitive status.

In the unadjusted model, 27 of the 44 associations remained significant, including all 14 in PVM_3 and all 6 M4 cognition associations, whereas all 14 PVM_1 associations were lost beyond random removal. In the contamination-adjusted model, 9 of the 35 remained significant (2 of 13 in PVM_3, 6 of 15 in M4, and 1 of 7 in PVM_1), including NeoC MMSE in both PVM_3 and M4 and NonNeoC MMSE in M4. In this model, 12 of the 15 pathology associations in PVM_3 and M4 were lost. Of the 7 cognition associations lost in these two arms, 4 remained below FDR=0.10: M4 CASI in NeoC and NonNeoC (FDR=0.061 and 0.073), M4 NeoC cognitive status (FDR=0.072), and PVM_3 NonNeoC MMSE (FDR=0.056).

The donor structure of Module post-IP% held in PVM_3, M4, and PVM_1 without the SA donors (Figure 16A), although the donor share of variance and split-half r_s_ fell by more than random removal in PVM_3 and PVM_1. The between-donor r_s_ with the intronic rate held in PVM_3 and PVM_1 and was lost beyond random removal in M4. Within donor, the PVM_3 associations with the IBA1 measures held (Figure 16B; Figure 14A). Between donors, the global CPS and global pTau CPS associations held in PVM_3 and M4, and all other CPS associations were lost (Figure 14B). With %MT in the same model, no new Module only verdict appeared (Figures 14C and 14D). Against the nuclear-encoded panels, 88% of the PVM_3 verdicts held, against 19% in M4 and fewer than 1% in PVM_1 (Figure 15A).

Collectively, the donor structure of Module post-IP% and its NeoC MMSE association in PVM_3 and M4 did not depend on the SA donors, although removing them weakened the clinical associations. As the SA donors had steeper memory decline, more cytosolic RNA, and lower data quality than other High-ADNC donors [19], these results may reflect either.

### 3.14 M4 against PVM_3

This section tested whether selecting the M4 cells added anything to PVM_3 (M4 is a subset of PVM_3). First, the M4 share (each donor’s M4 cells as a percentage of its PVM_3 cells) was added to the M4 clinical model (Supplementary Figure S20; Supplementary Table S18). M4 Module post-IP% retained its MMSE and CASI associations in both region sets at cov4 and +BTC (all FDR≤0.011), whereas the M4 share (significant for NeoC MMSE and CASI when tested alone) in the same models was not significant (all FDR≥0.112). Thus, the M4 association was not explained by the number of M4 cells.

Second, to test whether the M4 cells differed from other PVM_3 cells, they were compared with 10,000 random PVM_3 subsets with the same number of cells per donor and region (Supplementary Figure S21; Supplementary Table S18). The M4 cells did not differ from the random subsets in their association with cognition (all FDR≥0.106; Supplementary Figure S21A) or in how closely their Module post-IP% followed %MT (both FDR≥0.241; Supplementary Figure S21B). M4 was also the only main arm in which Module post-IP% and %MT could be separated (Figure 14D), and there it yielded three Module only verdicts. The random subsets could also be separated in every draw, and at least one in three gave the same Module only verdict for each of these comparisons (34.5-52.1% of draws; Supplementary Figure S21C). Taken together, the M4 cells did not differ from random PVM_3 cells of the same number in their cognition associations or their relation to %MT, and their Module only verdicts reflected the smaller number of cells.

### 3.15 Summary of all tests on Module post-IP%

Figure 17A summarizes the workflow, and Figure 17B shows the 15 tests of Module post-IP% for MMSE, CASI, Thal phase, and Braak stage on each arm, with NeoC and NonNeoC combined (Supplementary Table S19 also provides results of each region set separately). Tests 7 and 11 were unreadable in PVM_3 and PVM_1 (VIF≥5), thus the tallies counted the other 11 of tests 1-13 on every arm. PVM_3 held more tests than M4 in both region sets for MMSE (10 against 9 of 11), CASI (10 against 8), and Braak stage (8 against 4), and one fewer for Thal phase (6 against 7). Among the main arms, M4 lost the most models against the nuclear-encoded panels (test 12: 16 of 32 models held for MMSE and 14 of 32 models held for CASI, against 32 of 32 models held on PVM_3). PVM_1 held 7 of 11 tests in both region sets for MMSE and CASI, and HKG8 held none. On the M4-only tests, the M4 cognition association held with the M4 share in the model (test 14) and did not exceed random PVM_3 cells (test 15). Hence, PVM_3 carried the cognition association most consistently, without the need for marker-based subsetting such as M4.

**Figure 17.**
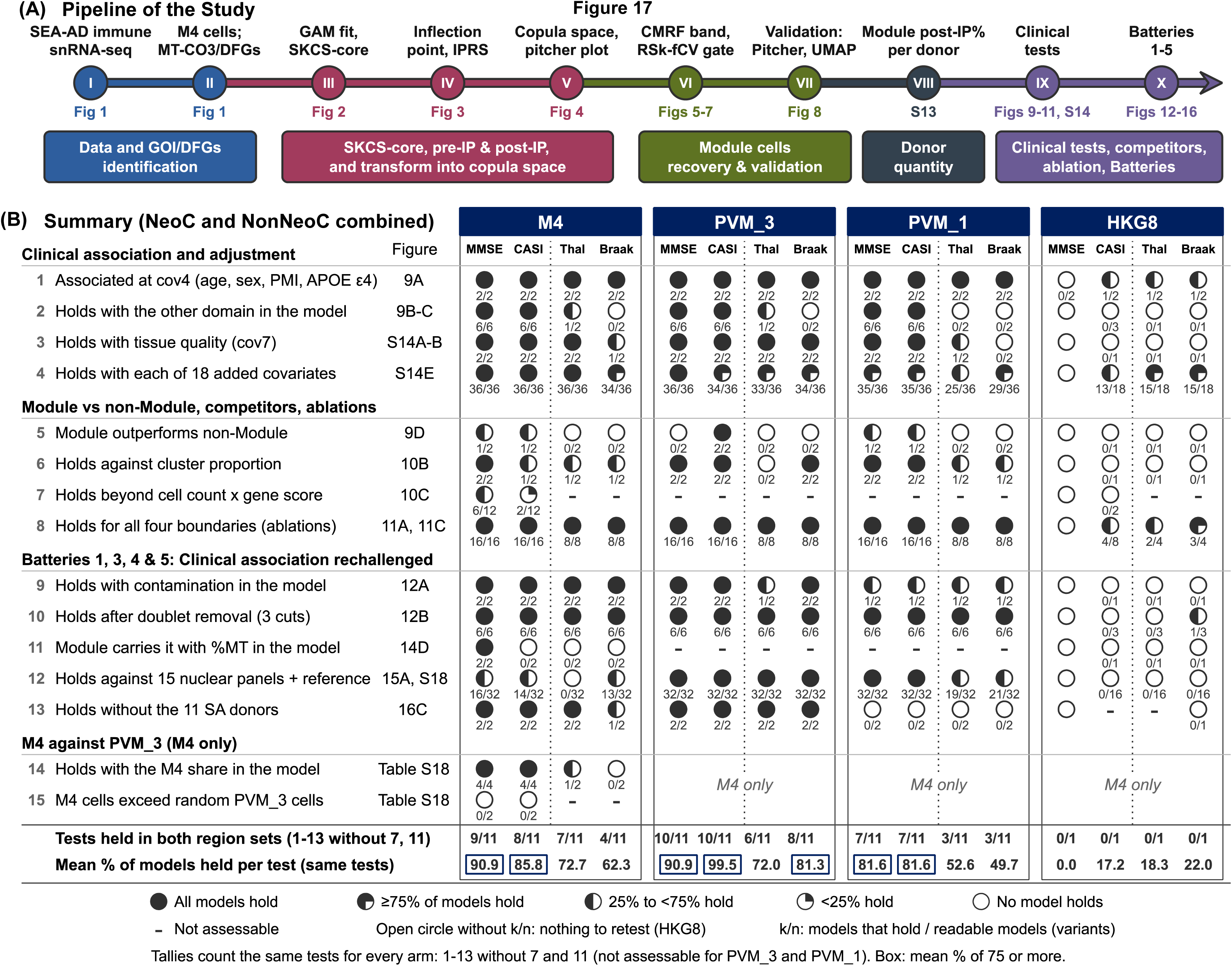
Summary of the workflow and of all tests on Module post-IP%. (A) Workflow from the SEA-AD data to the clinical tests and Batteries 1 to 5, with the figure for each step. (B) Fifteen tests for MMSE, CASI, Thal phase, and Braak stage per arm, NeoC and NonNeoC combined. Each circle is one test and outcome; k/n: models held over readable models; fill: percentage held (full, all; three-quarter, ≥75%; half, 25% to <75%; quarter, <25%; open, none). An open circle without k/n marks an outcome with no cov4 association, not retested; a hyphen marks a test that could not be assessed. Tests 14 and 15 apply to M4 only (Supplementary Figures S20 and S21). The summary lines show the tests held in both region sets and the mean percentage of models held per test for each arm and outcome. Boxes mark means of at least 75% (Supplementary Methods S15; Supplementary Table S19).

### 3.16 External validation 1: the *MT-CO3* module and its IP replicated in BA9

The *MT-CO3* module and its IP replicated in the BA9 of external dataset 1 [30] (Figure 18; Supplementary Table S20). Five of the eight DFGs passed tier 1 with *MT-CO3* (r_s_: 0.68-0.73 and τ_b_: 0.52-0.56), and two more passed tier 2, thus the DFGs replicated at the SEA-AD thresholds (Figure 18A). The r_s_ of these five were above the 0.577 expected at this depth from SEA-AD. Furthermore, cDFG rose with *MT-CO3* (DE=57.8%), but cHKG did not (DE=0.2%; Figure 18B). IPx was 3.10 (95% CI: 3.04-3.16) and IPy was 2.02, with an IPRS of 0.754 (strong tier), which passed the gate (Figure 18C). The SKCS-core held 1,052 cells, CMRF recovered 219 cells, and the Module held 1,225 cells, of which 664 were post-IP (Figures 18D and 18E). The HKG8 arm had no IP (IPRS of 0, failed tier).

**Figure 18.**
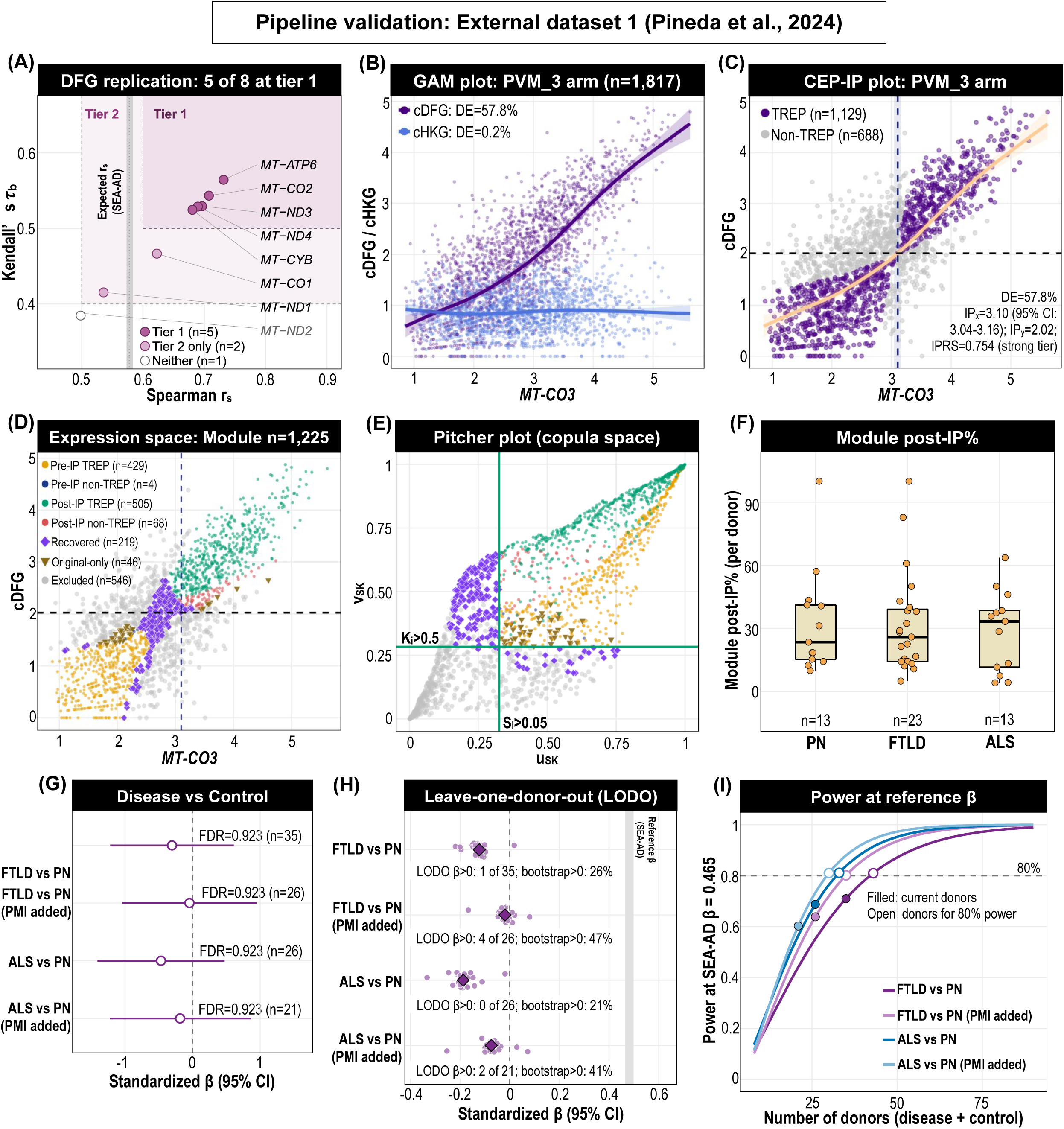
Pipeline validation in external dataset 1 (Pineda et al., 2024) [30]. BA9; panels B to E pooled the PVM_3 arm cells of all 58 donors with Micro-PVM_3 nuclei. (A) The expected r_s_ is from SEA-AD PVM_3 nuclei downsampled to 2,000 UMIs. (B) GAM plot. (C) CEP-IP plot; dashed lines: IP_x_ and IP_y_. (D) Module cells by group in expression space. (E) Pitcher plot with the SKCS-core gates. (F) Module post-IP% per donor with at least 5 Micro-PVM_3 nuclei; n: donors. (G) Standardized β (95% CI) of FTLD and ALS against PN on cov2 (age at death and sex) and on cov3 (cov2 plus PMI); one FTLD donor with no age at death is not in panels G to I. (H) β in LODO (points) and with all donors (diamonds); gray band: SEA-AD cognitive-status β. (I) Power to detect the SEA-AD NeoC β (0.465) by number of donors; open circles: donors needed for 80% power. Panels H and I are post hoc (Supplementary Table S20).

Module post-IP% had a median of 23.5% in PN, 25.9% in FTLD, and 33.3% in ALS (Figure 18F). Adjusted for cov2, Module post-IP% did not differ from PN for FTLD (β=-0.122, 95% CI: -0.488 to 0.243; 22 FTLD and 13 PN donors) or ALS (β=-0.187, 95% CI: -0.564 to 0.189; 13 ALS and 13 PN donors), nor with PMI added (β=-0.019 and -0.074; 8 PN donors), with FDR=0.923 for all four comparisons (Figure 18G). In the post hoc analyses, 1 of 35 LODO refits and 26% of the bootstrap β of the FTLD comparison were above 0 (Figure 18H). The SEA-AD NeoC β (0.465) was outside the 95% CI of all four comparisons. At this β, 80% power would need 43 donors for FTLD and 33 for ALS, against the 35 and 26 available (Figure 18I).

### 3.17 External validation 2: the *MT-CO3* module and its IP replicated in the insula

The *MT-CO3* module and its IP replicated in the insula of another snRNA-seq dataset [31] (Figure 19; Supplementary Table S21). Five of the eight DFGs passed tier 1 with *MT-CO3* (r_s_=0.76-0.79 and *τ*_b_=0.57-0.59), and the other three passed tier 2; thus, the DFGs replicated at the SEA-AD thresholds (Figure 19A). The r_s_ of these five were above the 0.630 expected at this depth from SEA-AD. Additionally, cDFG rose with *MT-CO3* (DE=64.3%), whereas cHKG did not (DE=0%; Figure 19B). IP_x_ was 3.19 (95% CI: 3.13-3.25) and IP_y_ was 2.40, with an IPRS of 0.758 (strong tier) (Figure 19C). The SKCS-core held 2,229 cells, CMRF recovered 430 cells, and the Module held 2,578 cells, where 1,341 were post-IP (Figures 19D and 19E). The HKG8 arm had no IP (IPRS of 0.200, failed tier).

**Figure 19.**
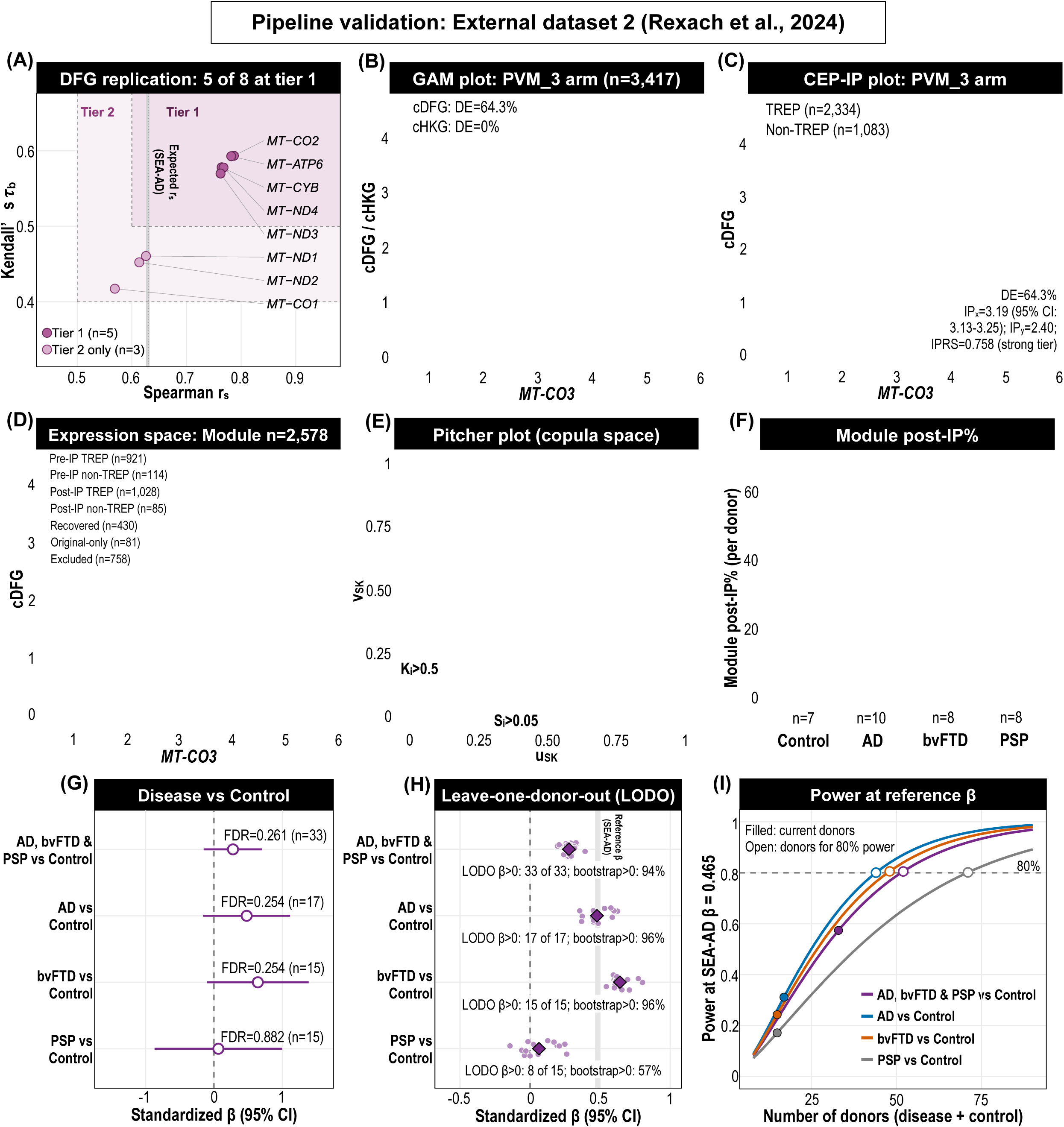
Pipeline validation in external dataset 2 (Rexach et al., 2024) [31]. Insular cortex; panels B to E pool the PVM_3 arm cells of all 37 donors. (A) r_s_ and *τ*_b_ of *MT-CO3* with each of the eight DFGs (33 donors with at least 5 Micro-PVM_3 nuclei); shaded areas: tiers 1 and 2; band: the expected r_s_ at 3,000 UMIs from SEA-AD. (B) GAM plot. (C) CEP-IP plot; dashed lines: IP_x_ and IP_y_. (D) Module cells by group in expression space. (E) Pitcher plot with the SKCS-core gates. (F) Module post-IP% per donor; n: donors. (G) Standardized β (95% CI) against controls on cov3, with FDR over the four contrasts. (H) LODO β (points), all-donor β (diamonds), and the share of bootstrap β above 0; gray band: SEA-AD cognitive-status β. (I) Power at the SEA-AD NeoC β (0.465); open circles: donors needed for 80% power. Panels H and I are post hoc analyses (Supplementary Table S21).

Module post-IP% had a median of 20.0% in controls, 28.2% in AD, 47.2% in bvFTD, and 26.8% in PSP (Figure 19F). Adjusted for cov3, the β was positive for all three diagnoses combined against controls (β=0.277, FDR=0.261), for AD (β=0.478, FDR=0.254), and for bvFTD (β=0.641, FDR=0.254), and near zero for PSP (β=0.064, FDR=0.882) (Figure 19G). In the post hoc analyses, the β of the diseases combined, AD, and bvFTD comparisons stayed positive in every LODO refit (33 of 33, 17 of 17, and 15 of 15), and 94%, 96%, and 96% of their bootstrap β were above 0, against 8 of 15 refits and 57% for PSP (Figure 19H). The SEA-AD NeoC β (0.465) was within the 95% CI of every comparison. At this β, 80% power would need 52 donors for the combined comparison and 44, 48, and 71 donors for AD, bvFTD, and PSP, against the 33, 17, 15, and 15 available, respectively (Figure 19I).

## 4. Discussion

The mitochondrial DNA (mtDNA) encodes 13 protein-coding genes, 12 of which are located on the heavy strand, while only MT-ND6, which is not part of this study’s Module DFGs, is located on the light strand [37, 38]. Heavy strand promoter transcription produces long polycistronic RNAs that are then cleaved into individual RNAs [39, 40], and heavy strand mtDNA genes are thus transcribed together. The transcription elongation factor TEFM helps mitochondrial RNA polymerase continue transcription without terminating prematurely [41, 42]. However, the polymerase often pauses after guanine-rich sequences [43], and steady-state levels of mitochondrial mRNAs are mainly controlled post-transcriptionally [39]. Hence, the heavy strand genes rise and fall together while their ratios vary, which may reflect the monotone relationship among the Module genes.

Empirical copulas were utilized in this study to define the CMRF-derived Module cells. The underlying statistics used were Spearman’s r_s_ and Kendall’s *τ*_b_ that utilize only ranks, unchanged by log transformation of normalized values. Scaling the ranks to between 0 and 1 yields the empirical copula, which separates the dependence between two genes from each gene’s own distribution. In contrast, population-level estimates assume one relationship for all samples or cells, missing heterogeneity. Copulas have modeled pairwise gene dependence in single-cell data [13, 44], and other methods estimate gene-gene association per sample [45] or per cell [46, 47]. Here, r_s_ and *τ*_b_ coefficients were split into per-cell scores, S_i_ and K_i_. This allowed CMRF to score each cell by its own contribution to the monotone relationship and gate on that score. The S_i_/K_i_ cell-level scoring was key to identifying the SKCS-core (the cells that followed the monotone relationship most closely, used to calibrate the GAM spline) and enabled the pitcher plot, in which the Recovered cells bridged the pre-IP and post-IP spines.

CMRF is similar to gating in flow cytometry, where a boundary is drawn around a cell population on a biaxial plot. Automated tools, such as flowDensity, perform biaxial gating by setting a density-based cutoff for each of the two markers [48]. Single-cell tools that provide a continuous gene-set score per cell [33, 34] or trajectory-based ordering of cells [49] are not primarily designed to group cells by pairwise gene relationships. Conversely, CMRF integrates a Gaussian kernel-based adaptive band (widened at the IP) with a cross-validated gate (RSk-fCV) to remove monotone-incoherent cells one at a time. When ablated, simpler boundaries provided the same cognition association. However, CMRF was the only boundary whose discarded cells showed no significant clinical association, and the only approach here that tested each candidate cell against the monotone fit. The cognition association of CMRF Module post-IP% therefore did not depend on the cells that CMRF discarded. The simpler boundaries yielded similar donor-level associations, but each discarded some signal-bearing cells.

Module post-IP% was associated with worse cognition (MMSE and CASI) in all three main arms and both region sets, and the association remained after cov4, tissue quality (cov7), pathology, or non-Module post-IP% adjustments. Green et al. described Mic.13 as a lipid-associated state enriched for the mouse DAM2 signature genes, and tau mediated most of its association with cognitive decline [26]. In the SEA-AD dataset, SA donors showed steeper memory decline and contained more cytosolic RNA such as transcripts from mitochondrially encoded genes [19]; without these SA donors, the cognition associations in the cov4 model held in PVM_3 and M4 but not in PVM_1. Furthermore, mitochondria can accumulate into plaques in AD brains, which co-develop with amyloid or emerge independently at early stages [50], thus plaque and tangle measures may not capture all mitochondrial pathology in AD.

Per-cell %MT is commonly used in QC to exclude nuclei with high mitochondrial content, hence a %MT cutoff may remove the Module cells. In the study by Green et al. that discovered Mic.13, nuclei with more than 10% mitochondrial RNA were removed [26]. Here, no per-cell %MT threshold reproduced the Module cells in PVM_3 or M4 (AUC≤0.586). As the Module cells had %MT below 10% (Figure 15B), a per-cell cutoff such as the 10% used by Green et al. is unlikely to remove the Module. The Module was detectable here because the mitochondrial genes were retained in the SEA-AD data. Studies that remove the mitochondrial genes or set per-cell cutoffs near 1% may not detect it.

M4 was the primary arm when the pipeline was built, and serves as the example in Figures 2 to 7 because PVM_3 holds about six times as many cells, which would crowd these cell-level scatter plots. PVM_3 was recommended only after the clinical analyses and the final comparison of M4 with PVM_3. Since the cognition associations of the M4 cells did not differ from those of random PVM_3 cells of the same number, the M4 marker-based subset did not improve on random subsets of the full supertype. Using the full supertype also removes the marker cutoff, an analyst choice that other studies would otherwise have to reproduce, and keeps the population tied to the published SEA-AD taxonomy. PVM_3 is therefore the recommended arm for further use of the framework. In PVM_3 and PVM_1, Module post-IP% and %MT were too collinear to be separated in the clinical models, hence whether the Module adds clinical information beyond %MT could not be tested in these supertypes.

This study has several limitations as follows:

1. The cognition associations were derived from a single cohort. In two external datasets, the *MT-CO3* module and its IP replicated, but the association of Module post-IP% with diagnosis was not confirmed. In Rexach et al. [31], the only external validation dataset with AD donors, the AD β against controls (0.478) was not significant with 7 controls but it was between the SEA-AD NeoC and NonNeoC cognitive-status β (0.465 and 0.499). The Gerrits et al. [24] dataset, used here for the technical checks of Battery 1, lacks cognition data and has 18 donors, below the 25-donor floor used in this study. Subsampling showed that the MMSE association required approximately 25-40 donors (Figure 15D). Within this cohort, fewer donors had CASI scores (68-69) than MMSE scores (79-80), which lowered the power for CASI.
2. DecontX was run in reduced mode, estimated from the expression profiles of eight immune supertypes. Empty-droplet correction requires the raw sequencing data, which are under controlled access on the AD Knowledge Portal (Synapse ID: syn26223298), and was not performed in this study.
3. Mitochondrial genes are transcribed together [39], and ambient RNA is rich in their transcripts (Figure 12E). Here, ambient RNA contributed to the mitochondrial co-expression, but modestly. In the external Gerrits dataset, nuclei in the top ambient-index tertile showed higher co-expression than those in the bottom tertile (Δr_s_=0.060), yet r_s_ rose from 0.267 to 0.366 once the ambient index and depth were removed. Observations in the SEA-AD dataset agreed, as PVM_3’s *MT-CO3*/DFG co-expression r_s_ fell to within Gerrits’s r_s_ IQR when matched to Gerrits’s lower sequencing depth and mitochondrial fraction (Battery 1). Moreover, in SEA-AD, module r_s_ did not rise with the ambient index or DecontX contamination, Module post-IP cells in PVM_3 were less contaminated than Module pre-IP cells, and the Module’s cognition associations held after DecontX adjustment for all three cognition outcomes in both region sets of PVM_3 and M4. Two further lines of evidence were independent of DecontX: i) ACTB, a highly expressed transcript that may contribute to ambient RNA, did not co-vary with *MT-CO3* (Figure 1G); ii) the offset share of mitochondrial UMIs (the depth-independent part expected of ambient RNA) did not differ between Module post-IP and pre-IP cells in PVM_3 (Figure 13E). However, a residual ambient contribution to Module post-IP% cannot be excluded until an empty-droplet correction such as CellBender [51] is applied to the SEA-AD raw data.
4. Module post-IP% shares variation with simpler gene scores (AUCell, UCell, and AddModuleScore) and %MT, hence their separate contributions to the clinical outcomes could be tested only on M4. Module post-IP% retained the M4 NeoC cognition associations against the gene scores, which may also reflect the subset effect of M4 (Supplementary Figure S21). In contrast, the gene scores carried some pathology associations that Module post-IP% did not. Within donor, no histology measure was associated with Module post-IP% alone (Module only) once %MT was held fixed, and adding the donor-mean %MT or sequencing saturation removed some cognition associations. Nonetheless, the gene scores and %MT provide each cell with a continuous value and require an added cutoff to define cell populations, whereas CMRF assigns each cell to a population (e.g., Module post-IP or non-Module) for downstream cell-level comparisons.
5. Despite the Batteries, the source of the Module’s mitochondrial transcripts remains unresolved. In isolated nuclei, mitochondrial transcripts may come from cytoplasm and mitochondria retained around the nucleus, from ambient RNA in the droplet [52], or from a true increase in cellular mitochondrial content (e.g., activated microglia) [53]. The Batteries favored retained cytoplasm. Post-IP Module cells carried more ER and ribosomal RNA and less nuclear-retained NEAT1 RNA than pre-IP Module cells (Supplementary Figure S15), and these cytoplasmic RNAs are depleted in microglial nuclei compared with whole cells [54]. Ambient RNA and lower nuclear RNA were likely minor contributors, and tissue quality may contribute in part, although the cognition associations held at cov7. Free mitochondria attaching to nuclei were not tested. Whether these microglia contain more mitochondria or retain more cytoplasm cannot be resolved with these data.
6. Experimental data are required for validation and to determine the origin of the mitochondrial transcripts:

i. For validation, untargeted spatial transcriptomics that captures mitochondrial transcripts is required to locate Module cells in tissue, but targeted spatial data cannot be used for this purpose, e.g., SEA-AD’s released MERFISH data of the MTG (140-gene panel) [19] and of the MEC and HIP (433 genes, neurotypical tissue) contain no mitochondrial gene.
ii. To test the origin, nuclei could be stained for attached mitochondria (e.g., TOMM20) and sorted before sequencing, as sorting removes extra-nuclear ambient RNA [55], or whole-cell and single-nucleus RNA-seq could be compared on tissue from the same donors [54]. These experiments are challenging: fresh human brain tissue is scarce, PMI and RNA quality vary [15, 19], and isolation protocols differ [56].
7. The study pipeline has several procedures, which may be demanding to run in full. To make this less challenging, the full code is publicly available, together with a container that fixes the software versions, the logs of the original runs, and the processed data needed to rerun the R analyses without the SEA-AD h5ad. In the container on a desktop computer (Intel Core i9-14900KF, 64 GB RAM, Fedora 44), the R analyses ran in about 2.5 hours, and the whole pipeline starting from the SEA-AD h5ad completed in about five hours. The reruns, including an independent test on Windows 11, reproduced the original findings (Section 2.10), and an R package of CEP-IP with CMRF (cepip) has been submitted to the Comprehensive R Archive Network (CRAN) so that the pipeline can be rerun on other datasets.

## 5. Conclusion

In conclusion, Module post-IP% tracked cognition in SEA-AD microglia, and no per-cell %MT cutoff reproduced the Module cells. Routine handling of mitochondrial reads in AD snRNA-seq studies, such as excluding the mitochondrial genes or applying %MT cutoffs close to 1%, may therefore remove a signal linked to cognition. Before such a cutoff is applied or the mitochondrial genes are excluded, mitochondrial reads can be checked with the released pipeline, its container, or the R package cepip, with and without the intended %MT cutoff. The origin of these mitochondrial transcripts remains unknown. Experiments on nuclei preparations and paired whole-cell data are needed to determine their origin, and external cohorts are needed to test whether the association with cognition replicates.

## Data and Code Availability

1. Source data

i. The SEA-AD immune snRNA-seq dataset (Multiregion release of 22 June 2026), with the quantitative neuropathology and CPS workbooks: https://brain-map.org/consortia/sea-ad/our-data
ii. Gerrits et al. dataset: GEO (GSE148822)
iii. External dataset 1 (Pineda et al.): SRA (PRJNA1073234), with the donor metadata on Synapse (syn51105515)
iv. External dataset 2 (Rexach et al.): CELLxGENE (dataset aa87f914-07e0-48aa-b915-8de906c95baf)
v. Inputs of the external datasets not available elsewhere (the Cell Ranger count matrices of external dataset 1 and the MapMyCells outputs of both): folder external_validation/ of the Hugging Face dataset (link in item 2 below).
2. Processed data

i. The data needed to rerun the R analyses from Part 9 without the SEA-AD h5ad (Part 2 donor clinical tables, Part 7 DecontX estimates, and Part 8 count matrices of the four arms; 14.9 GB), under the Allen Institute Terms of Use: https://huggingface.co/datasets/kahkengwong/CEP-IP_with_CMRF_in_SEA-AD
ii. This study used publicly released SEA-AD data and was conducted independently of the Allen Institute and the SEA-AD consortium.
3. Code

i. GitHub, under the MIT License: https://github.com/kahkengwong/CEP-IP_with_CMRF_in_SEA-AD
ii. Full repository code and the container images (SEA-AD and the two external datasets) on Zenodo: https://doi.org/10.5281/zenodo.23653156
iii. Contents: 23 Python scripts (Parts 1 to 8 and the Part 12 extraction step), 43 R scripts (Parts 9 to 13 and the Figure 17 summary), three shared R settings files, and the two settings files and two pre-specified reading files of the Batteries 1-5.
iv. Logs: each script writes a log of its inputs, settings, and checks. The logs of the original runs (194 from R and 21 from Python) and of the container reruns are also deposited.
4. Software environment and reproducibility

i. Versions: R 4.6.1, Bioconductor 3.23, and all 280 R packages in renv.lock, and the Python 3.13.14 package versions in requirements-python.txt
ii. Container: the Containerfile builds a Podman or Docker container with all packages at these versions and the scripts. Inside it, seaad-run all reruns the whole pipeline and generates the figures and supplementary tables, and seaad-run hf reruns R Parts 9 to 11 from the Hugging Face data.
iii. Tests: the author ran seaad-run all on Fedora 44, and an independent tester ran seaad-run hf on Windows 11. Both reruns reproduced the clinical associations (Section 2.10; Supplementary Methods A3).
iv. Instructions: the commands are in Supplementary Methods A3, and step-by-step instructions are in README_container.md in the GitHub repository.
5. R package cepip

i. The methods of this study (CEP-IP, SKCS, and CMRF), on GitHub under the MIT License: https://github.com/kahkengwong/cepip (submitted to the Comprehensive R Archive Network, CRAN)
ii. Example data (the PVM_3 and HKG8 cells, the donor table, and the DEG matrices): https://huggingface.co/datasets/kahkengwong/cepip_example
iii. With these data, the package script reproduce_seaad.R reruns the PVM_3 and HKG8 arms and checks 698 published values, and all of them agree.
6. Supplementary files

i. Supplementary Figures (one PDF file) and Supplementary Tables (21 Excel workbooks) on Zenodo: https://doi.org/10.5281/zenodo.23719390

## CRediT authorship contribution statement

Kah Keng Wong: Conceptualization, Data curation, Investigation, Formal analysis, Visualization, Validation, Software, Resources, Supervision, Writing - original draft, Writing - review & editing.

## Acknowledgements

The author thanks the School of Medical Sciences and the Research Creativity and Management Office (RCMO), Universiti Sains Malaysia, for their support. The author thanks Nik Mohd Asri Nik Amirah Auni for independently testing the reproducibility of the pipeline and the R package cepip. The main analyses used the SEA-AD data, and the author thanks the brain donors and their families and the SEA-AD consortium (RRID:SCR_023110; funded by the National Institute on Aging, U19AG060909) for making them publicly available. The author also thanks the authors and donors of the external datasets of Gerrits et al., Pineda et al., and Rexach et al.

## Supplementary figure legends

The Supplementary Figures (one PDF file) and the Supplementary Tables (Excel workbooks) are deposited on Zenodo (https://doi.org/10.5281/zenodo.23719390).

**Supplementary Figure S1.**
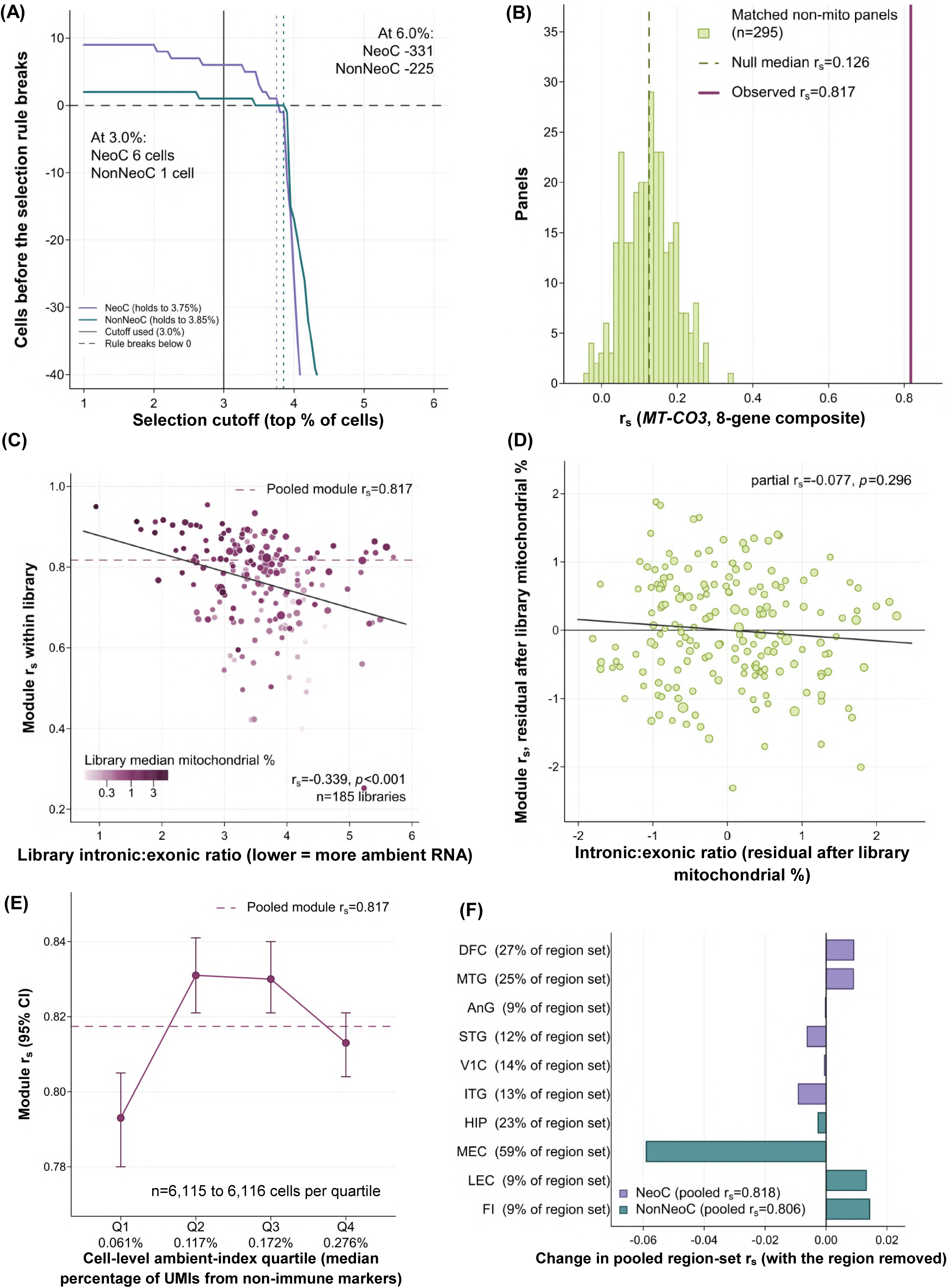
Technical controls behind Figure 1. (A) Selection-rule margin against the M4 cutoff (Supplementary Methods S1); dotted lines: limits of 3.75% (NeoC) and 3.85% (NonNeoC); solid line: 3.0% cutoff used. (B) Module r_s_ of 295 abundance-matched random non-mitochondrial panels (histogram; dashed line, median; solid line, observed). (C) Per-library module r_s_ against the intronic:exonic ratio (lower, more ambient RNA); point size: analyzed cells; color: median %MT; dashed line: pooled r_s_. (D) As C, after partialing out library %MT; solid lines: linear fit. (E) Module r_s_ by ambient-index quartile, with 95% CIs. (F) Change in pooled module r_s_ when each region is removed. Values are in Supplementary Tables S1 and S5.

**Supplementary Figure S2.**
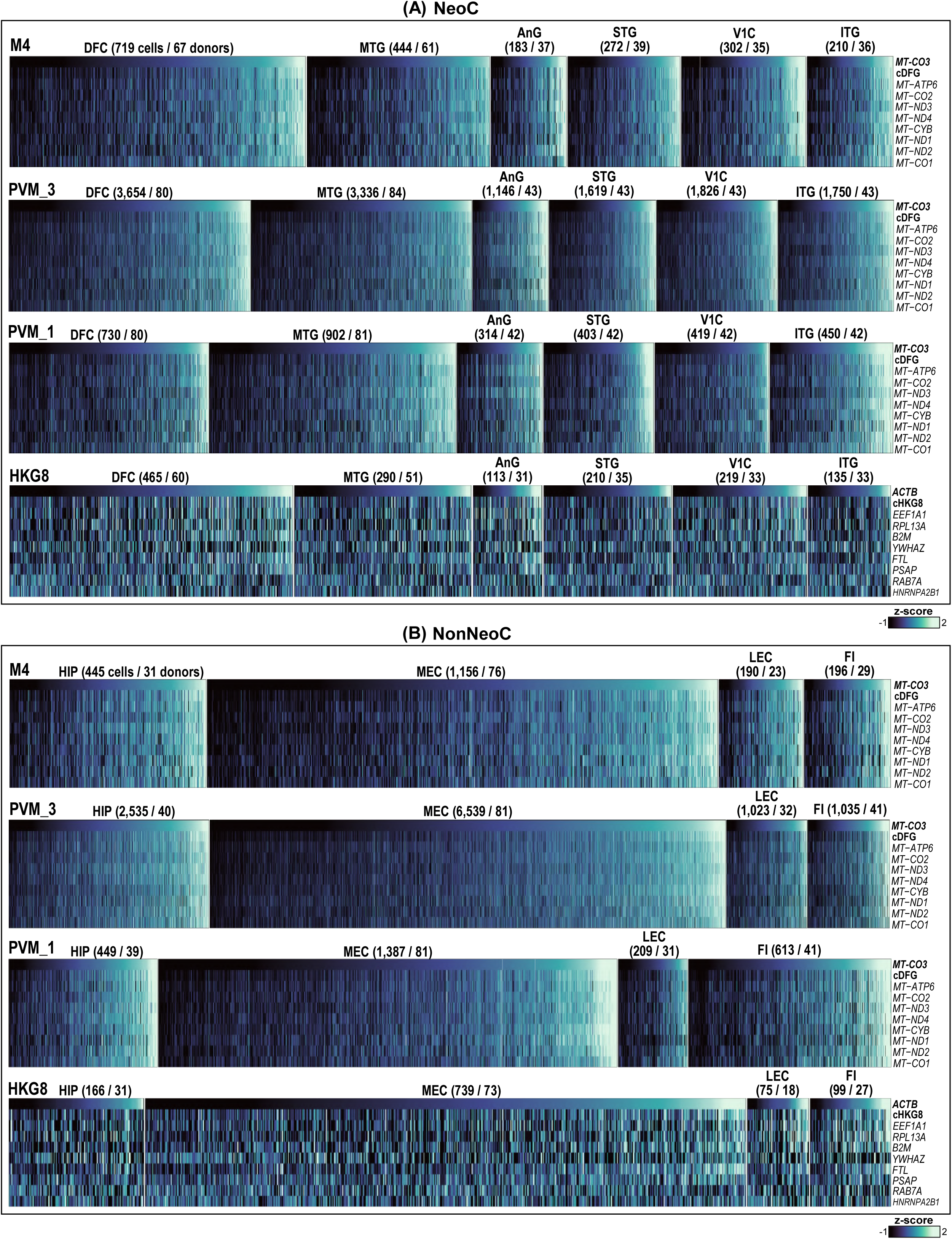
Heatmaps of M4, PVM_3, PVM_1, and HKG8 in NeoC (A) and NonNeoC (B). Cells are ordered by ascending GOI expression (*MT-CO3*; *ACTB* for HKG8) within each brain region; column labels give cells and donors per region. Values are per-gene z-scores over all cells of the region set, capped at -1 to +2 (Supplementary Table S7). cHKG8 is the composite of the eight housekeeping genes; the other arms’ housekeeping blocks are in Supplementary Figure S3.

**Supplementary Figure S3.**
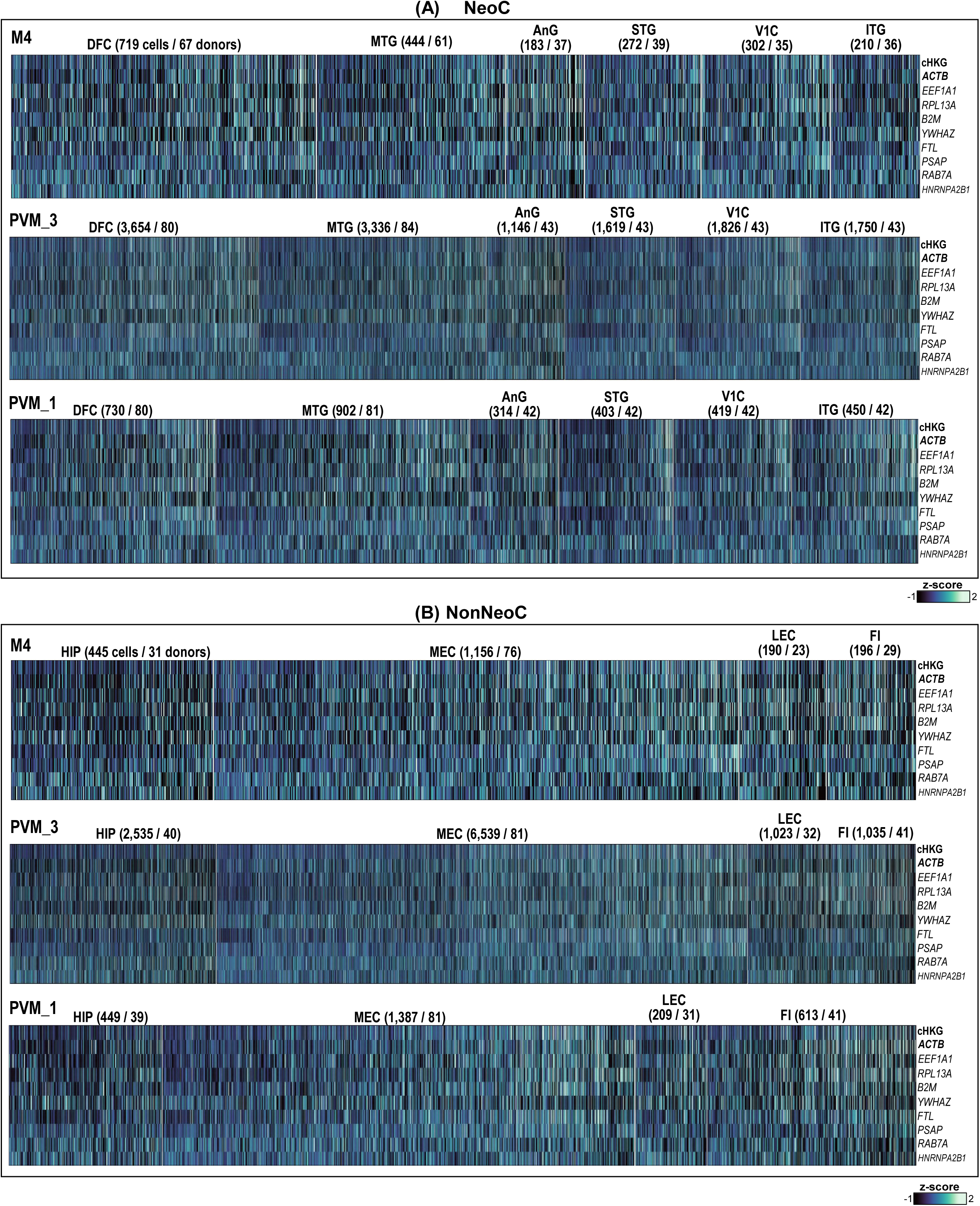
Heatmap of the housekeeping genes in M4, PVM_3, and PVM_1 in NeoC (A) and NonNeoC (B). Same cells and column order as Supplementary Figure S2 (ascending *MT-CO3* expression in each brain region). cHKG denotes the composite of the nine housekeeping genes, including *ACTB*. Values are in Supplementary Table S7.

**Supplementary Figure S4.**
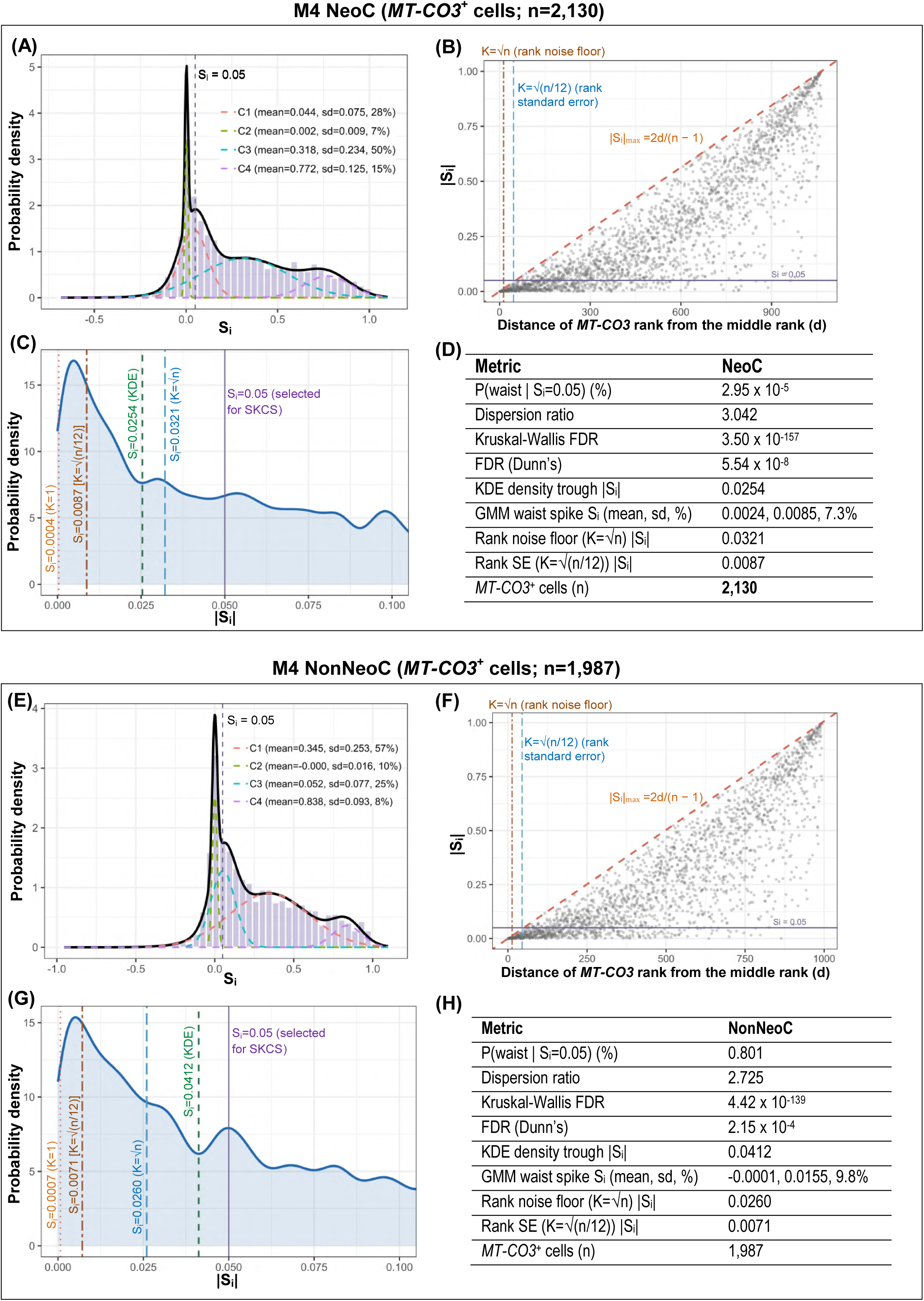
Justification of the S_i_>0.05 threshold in M4 *MT-CO3*^+^ cells, NeoC (A to D) and NonNeoC (E to H). (A and E) Distribution of S_i_ with a four-component Gaussian mixture model (GMM; C2, the peak at zero); the dashed line marks S_i_=0.05. (B and F) |S_i_| versus d, the distance of each cell’s *MT-CO3* rank from the middle rank, with the bound |*S_i_*| ≤ 2*d*⁄(*n* − 1) (dashed diagonal), vertical lines at K=√n and K=√(n/12), and S_i_=0.05 (horizontal line). (C and G) Probability density of |S_i_| between 0 and 0.1, with the largest |S_i_| of cells within K=1, √(n/12), and √n ranks of the middle rank, the trough of the kernel density estimate (KDE), and the selected S_i_=0.05. (D and H) Summary values: the posterior probability that a cell at S_i_=0.05 belongs to C2 (P(waist)); the dispersion ratio of cells with S_i_ in [0.04, 0.06) against the peak. Values are in Supplementary Table S7.

**Supplementary Figure S5.**
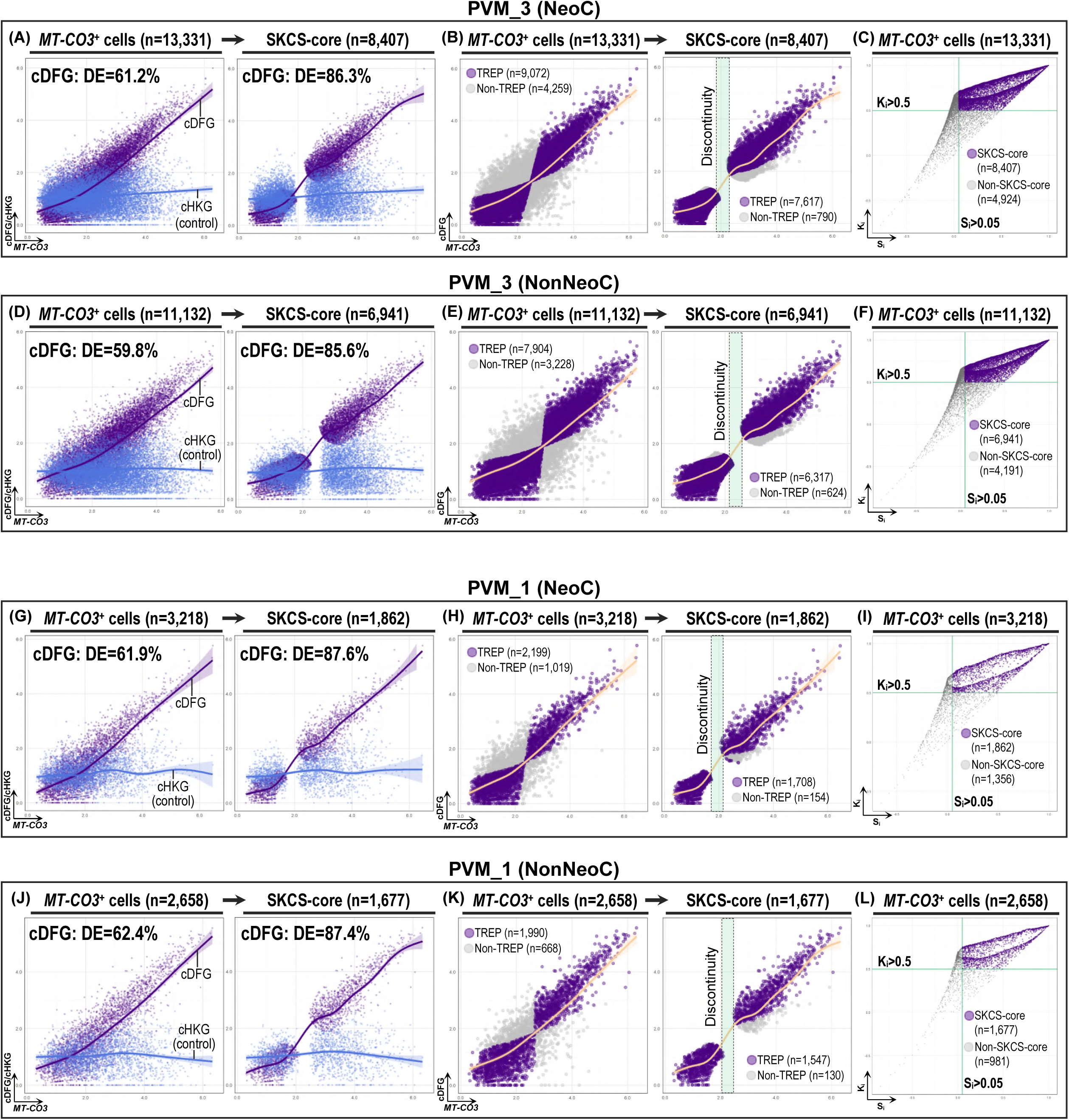
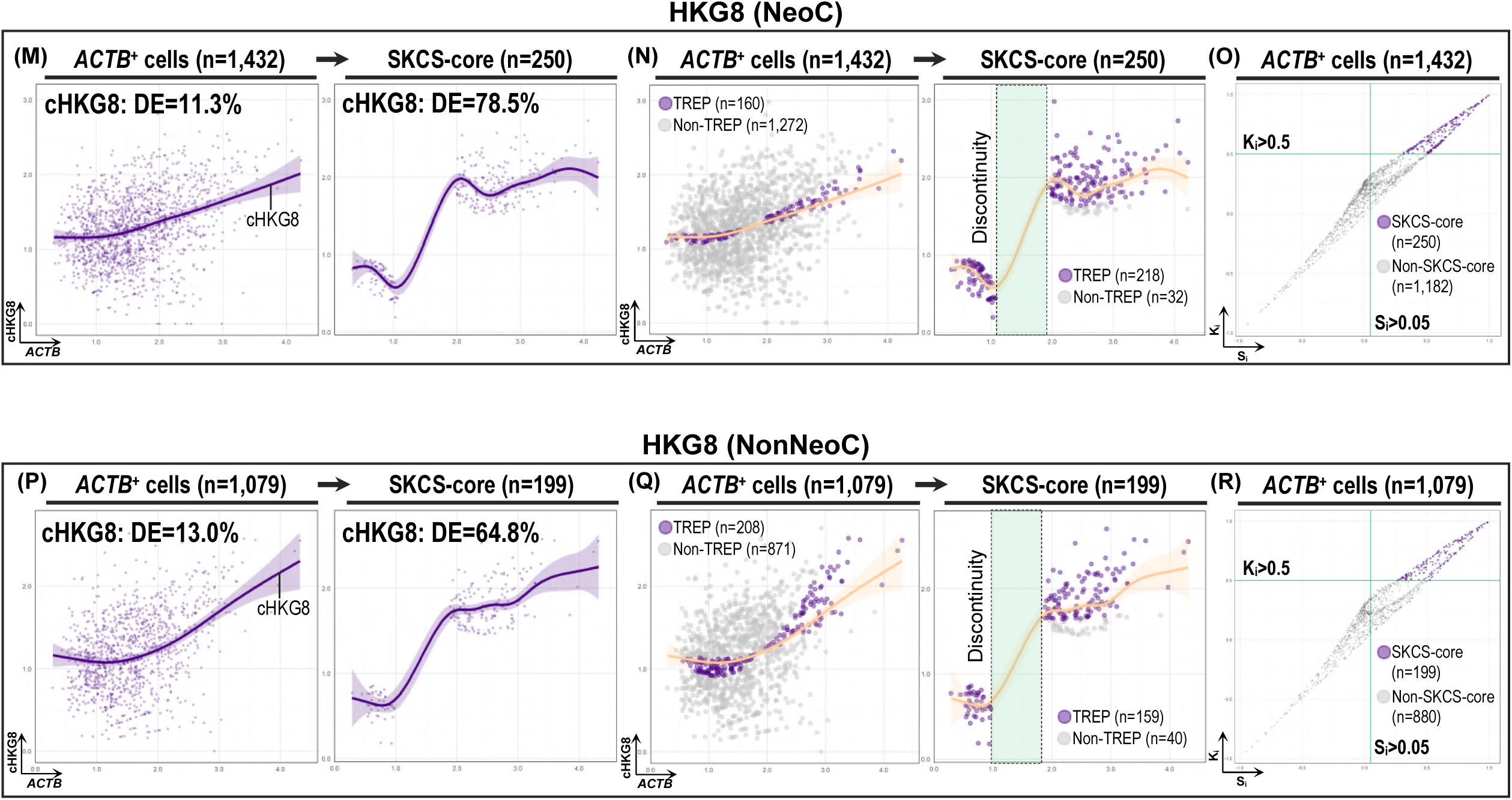
GAM, CEP-IP, and pitcher plots of PVM_3, PVM_1, and HKG8 NeoC and NonNeoC cells, as in Figure 2. (A to F) PVM_3, (G to L) PVM_1, and (M to R) HKG8: GAM plots (A, D, G, J, M, P), CEP-IP plots (B, E, H, K, N, Q), and pitcher plots (C, F, I, L, O, R) of GOI^+^ cells and SKCS-core cells. For HKG8, the GOI is *ACTB* and the composite is cHKG8 (the eight housekeeping genes), hence no separate housekeeping control is shown. Values are in Supplementary Table S7.

**Supplementary Figure S6.**
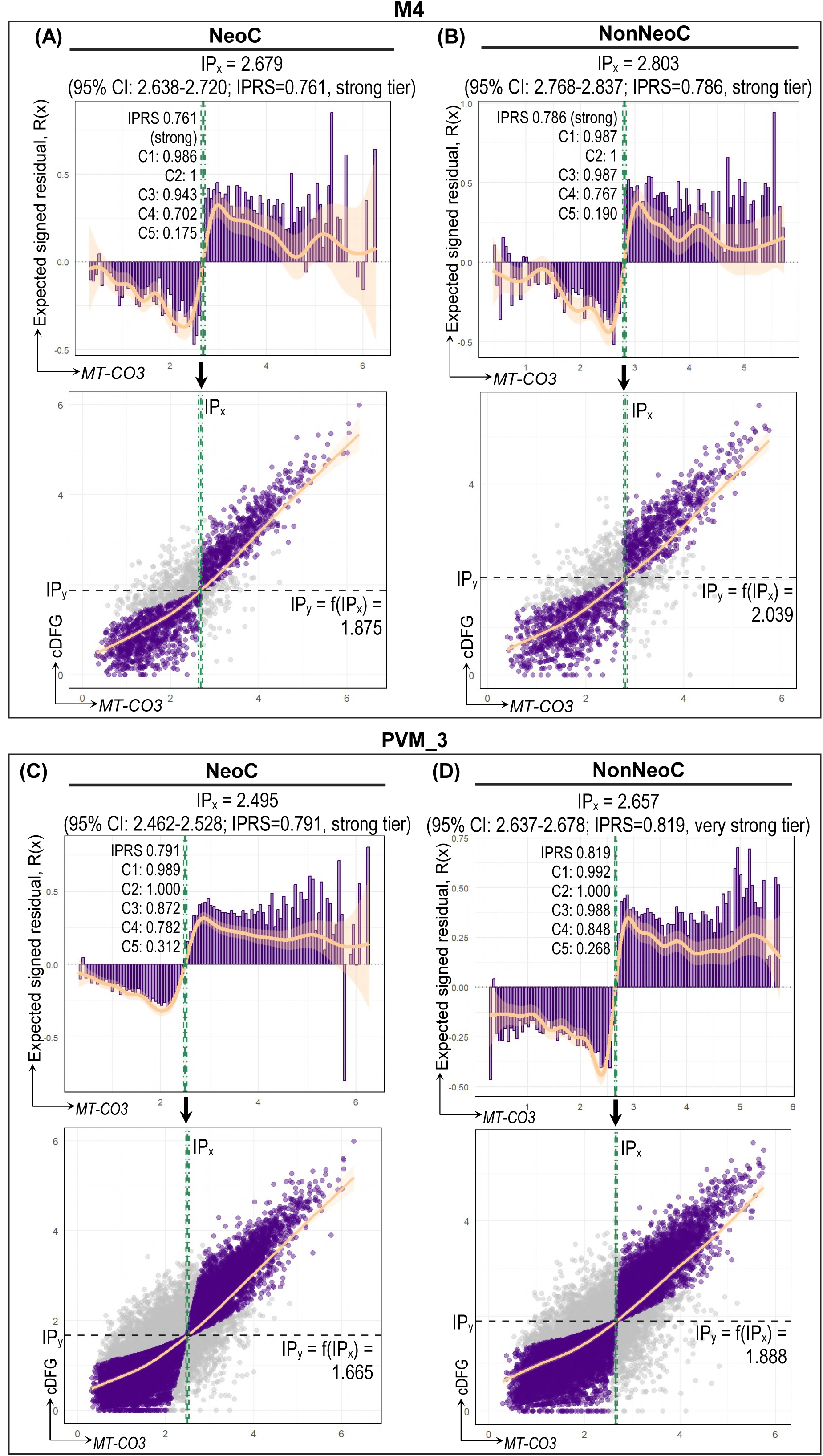
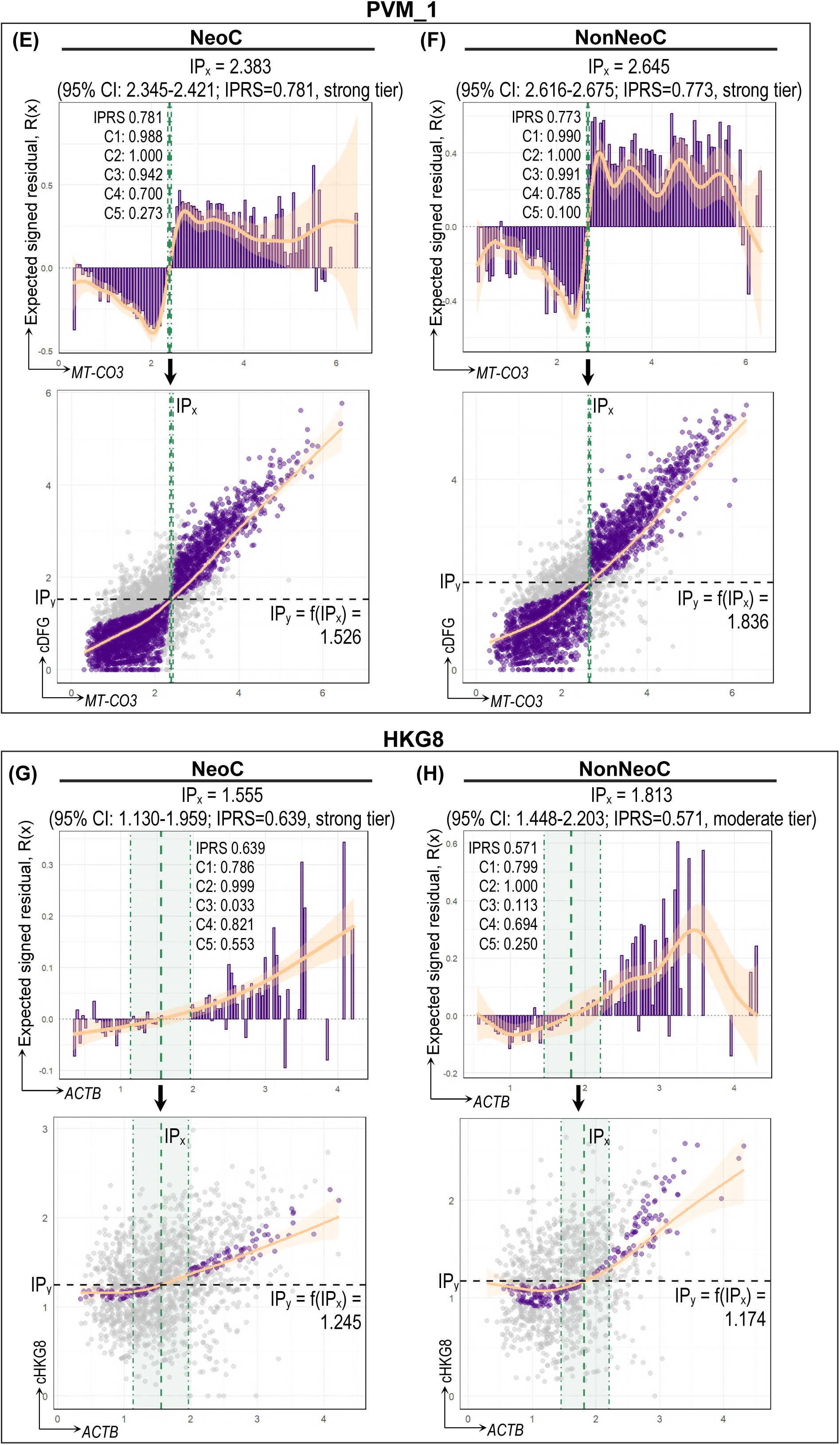
IP per arm and region set (Supplementary Table S8): M4 (A, B), PVM_3 (C, D), PVM_1 (E, F), and HKG8 (G, H), NeoC first in each pair. Top: binned mean TREP residual from the GAM fit (bars) and the residual GAM R(x) with 95% band; green dashed line: IP_x_ (the first negative-to-positive crossing of R(x)) with its 95% CI (dot-dashed). Bottom: composite against GOI (TREP cells purple), with the GAM fit (orange) and IP_y_ (black dashed line).

**Supplementary Figure S7.**
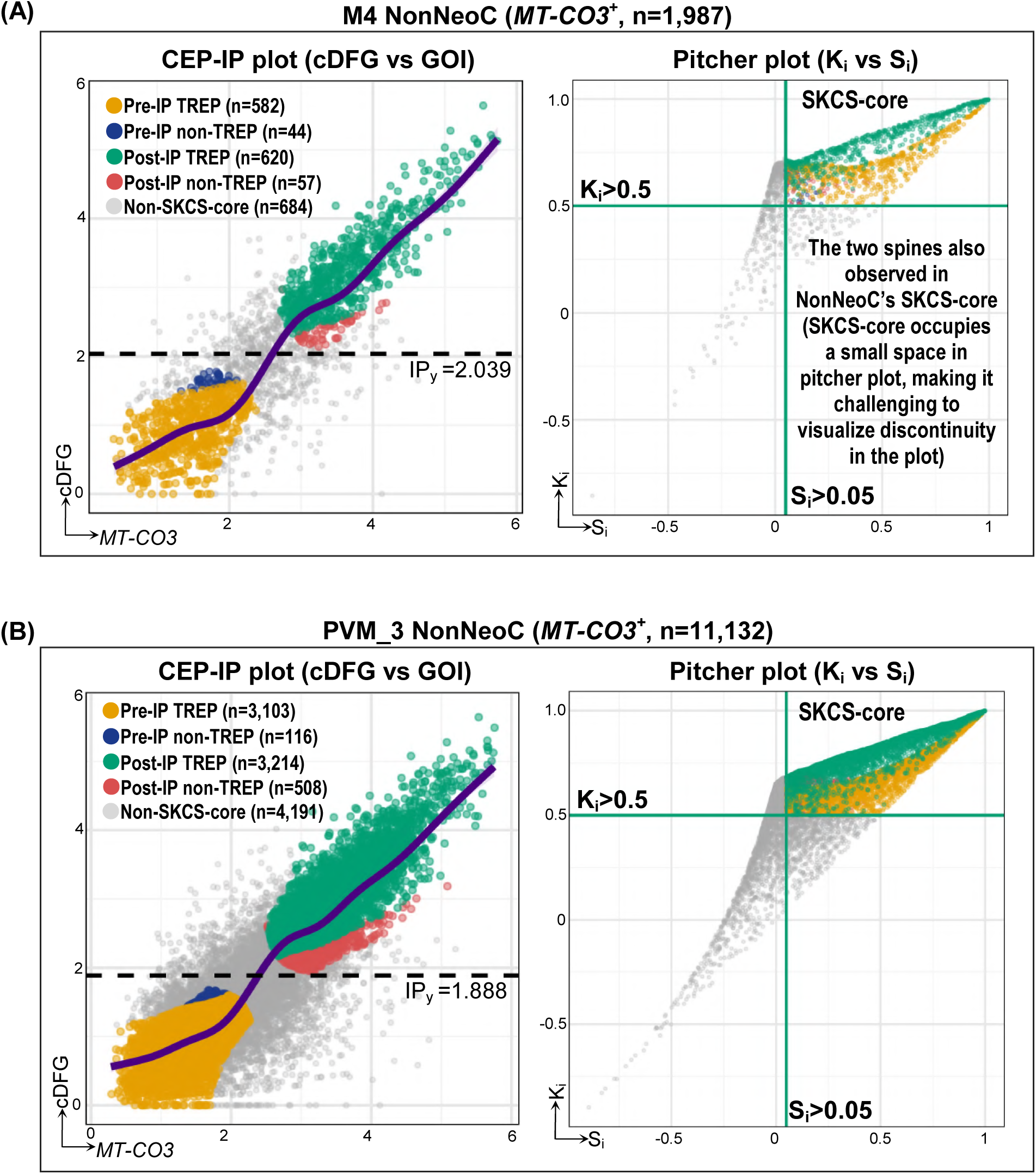
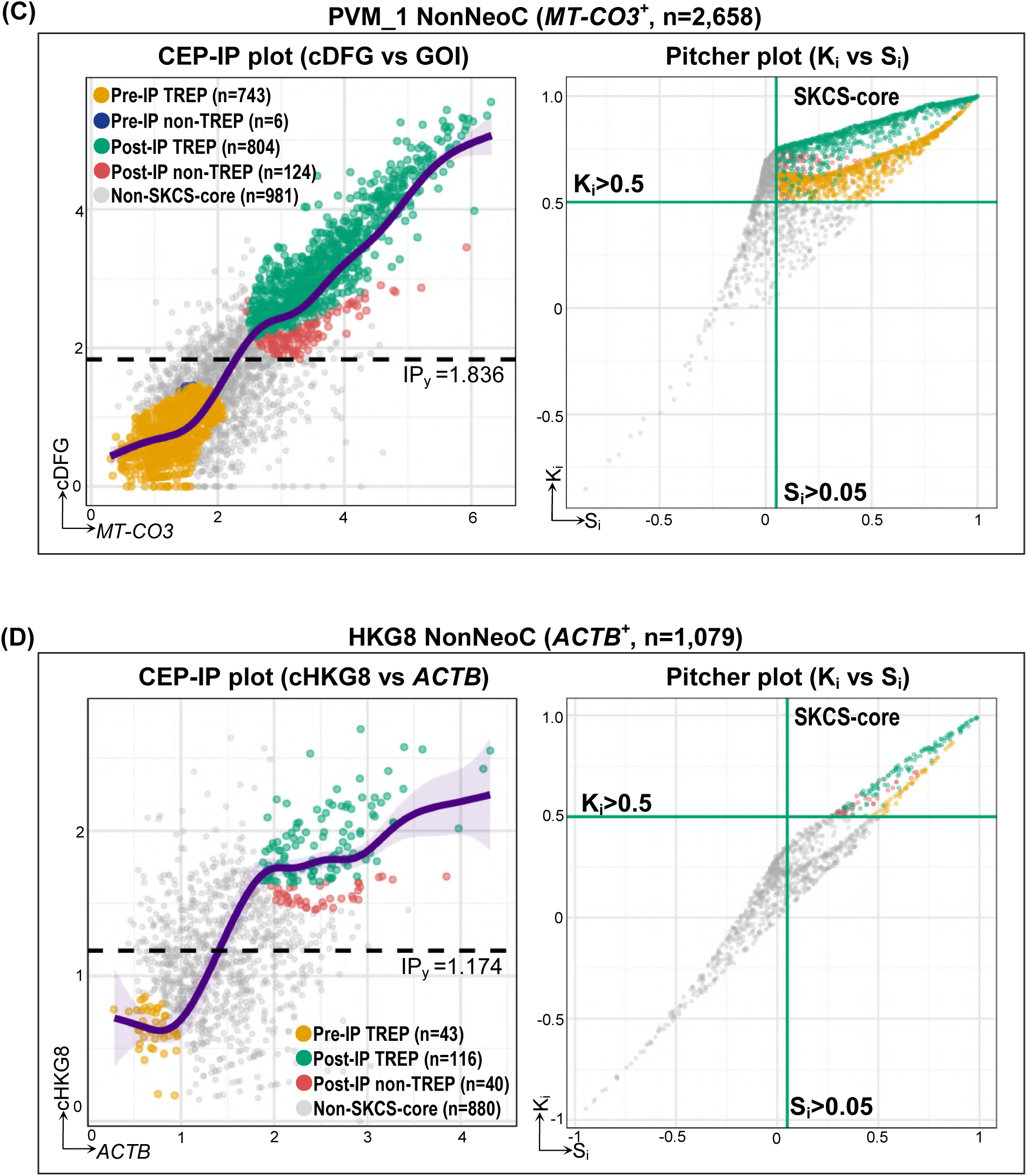
Distribution of M4 (A), PVM_3 (B), and PVM_1 (C) NonNeoC *MT-CO3*⁺ cells, and HKG8 (D) NonNeoC *ACTB*⁺ (control group) cells, according to the CEP-IP plot (left panel) and the pitcher plot (right panel). Details as in Figure 3. HKG8 has no pre-IP non-TREP cells in either region set. Values are in Supplementary Table S7.

**Supplementary Figure S8.**
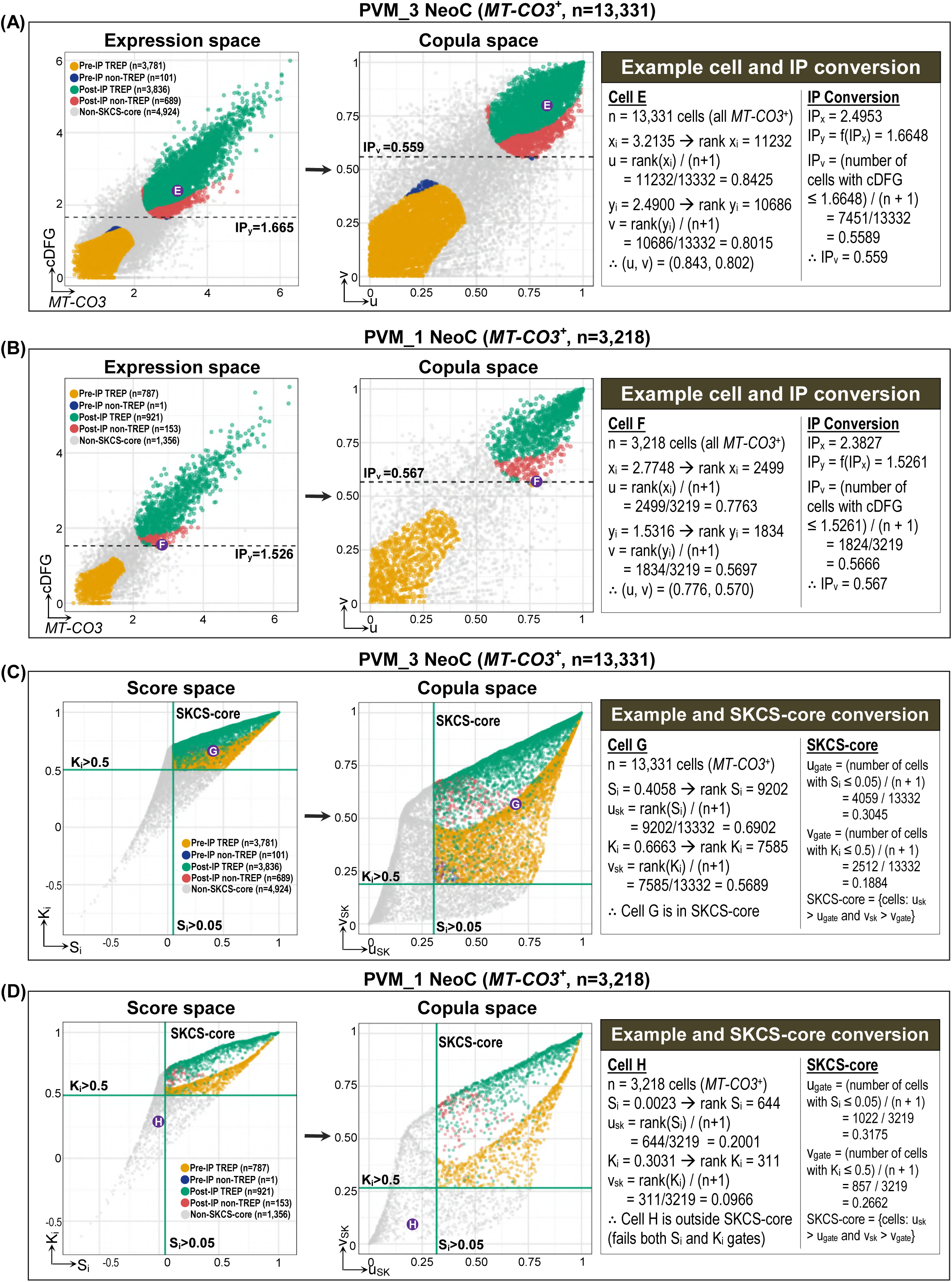
Distribution of PVM_3 (A, C) and PVM_1 (B, D) NeoC *MT-CO3*^+^ cells in expression space and score space, and in the copula space of each. Details as in Figure 4. (A, B) Expression space and its copula space, with cells E and F in the insets. (C, D) Score space and its copula space, with cells G and H in the insets; cell H lies outside the SKCS-core and fails both gates. Values are in Supplementary Tables S7 and S9.

**Supplementary Figure S9.**
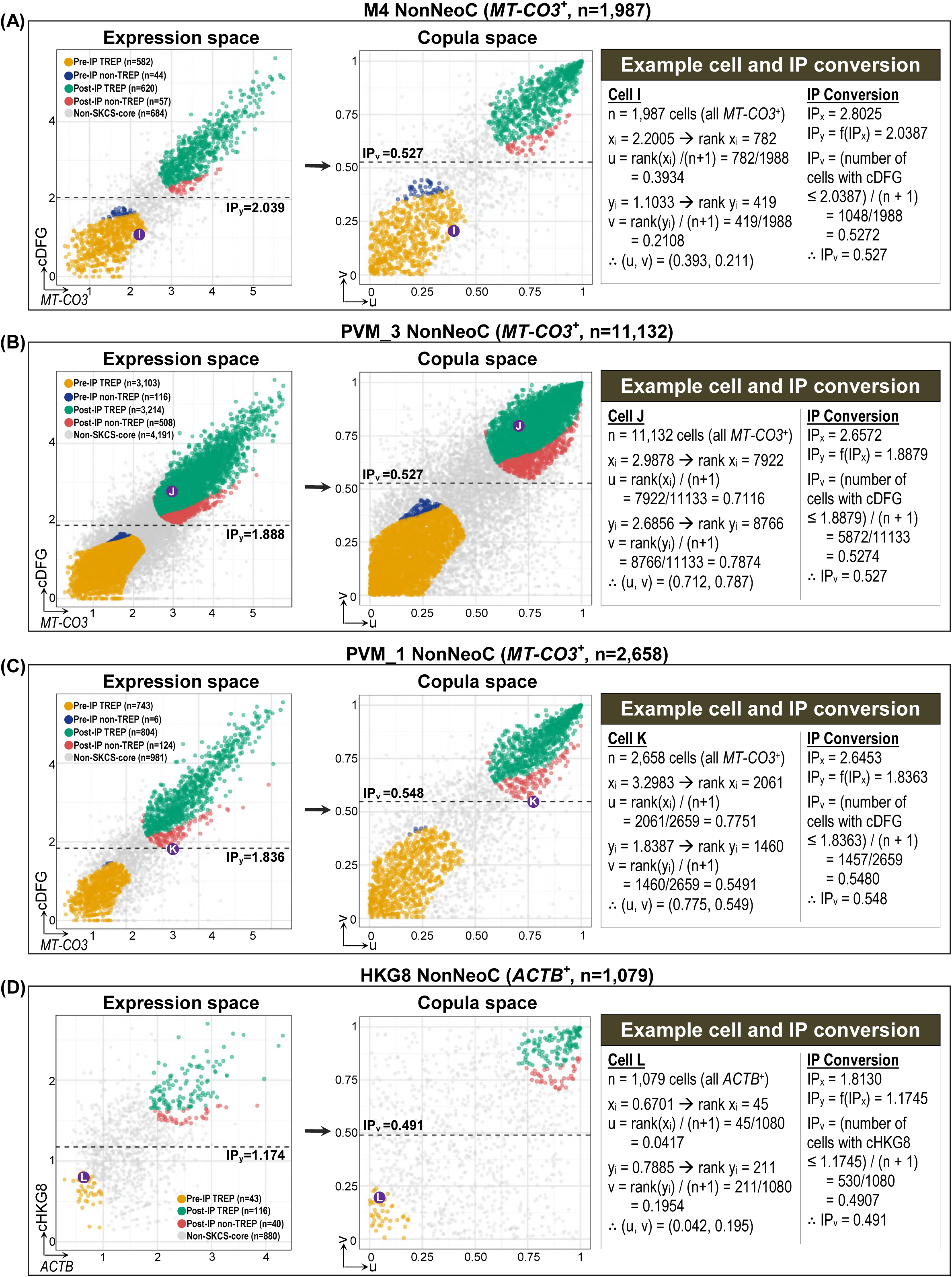
Distribution of M4 (A), PVM_3 (B), and PVM_1 (C) NonNeoC *MT-CO3*⁺ cells, and HKG8 (D) NonNeoC *ACTB*⁺ cells in expression space (cDFG or cHKG8 versus GOI; left) and copula space (ranks of the same two axes divided by n + 1; right). The inset converts one circled cell (I to L) and the IP into copula space; dashed line: IP_v_. Colors as in Figure 3. Values are in Supplementary Tables S7 and S9.

**Supplementary Figure S10.**
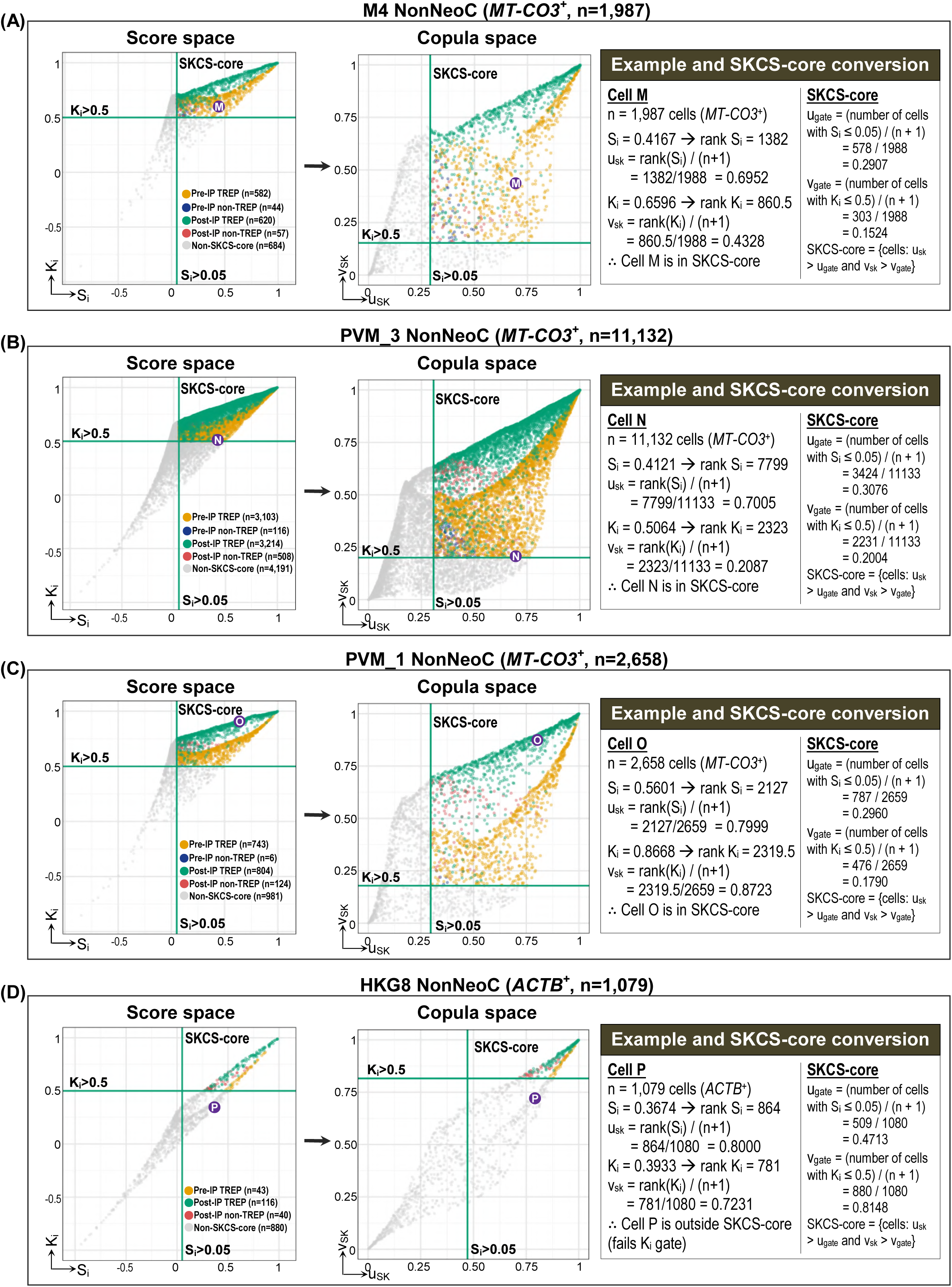
Distribution of M4 (A), PVM_3 (B), and PVM_1 (C) NonNeoC *MT-CO3*⁺ cells, and HKG8 (D) NonNeoC *ACTB*⁺ cells in score space (K_i_ versus S_i_; left) and copula space (ranks of the same two axes divided by n + 1; right). The inset converts one circled cell (M to P) and the SKCS thresholds (S_i_=0.05 and K_i_=0.5) into u_gate_ and v_gate_; SKCS-core cells have u_SK_>u_gate_ and v_SK_>v_gate_, except three cells (Supplementary Methods S6). Colors as in Figure 3; S_i_ as in Figure 4C and 4D. Values are in Supplementary Tables S7 and S9.

**Supplementary Figure S11.**
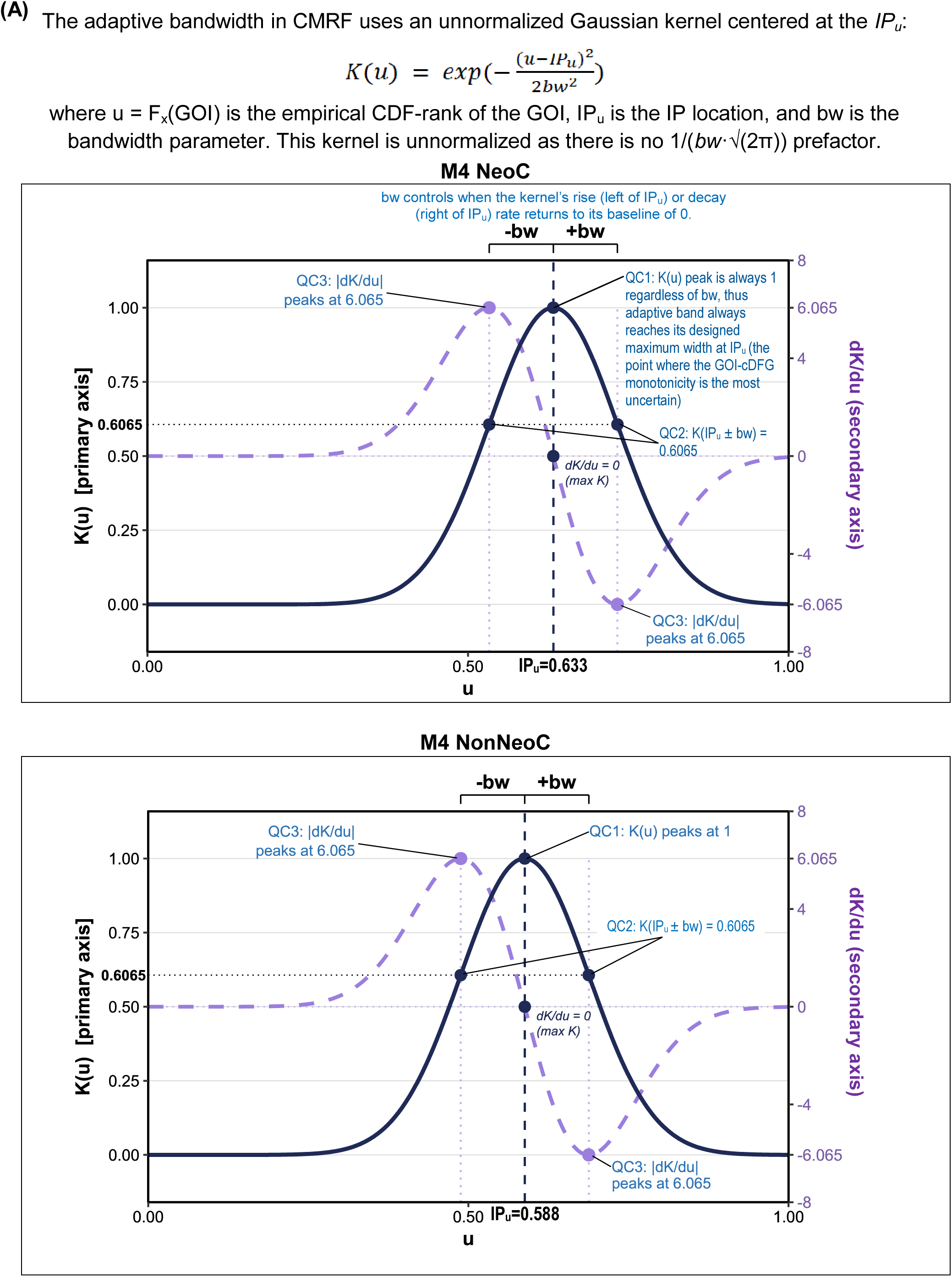
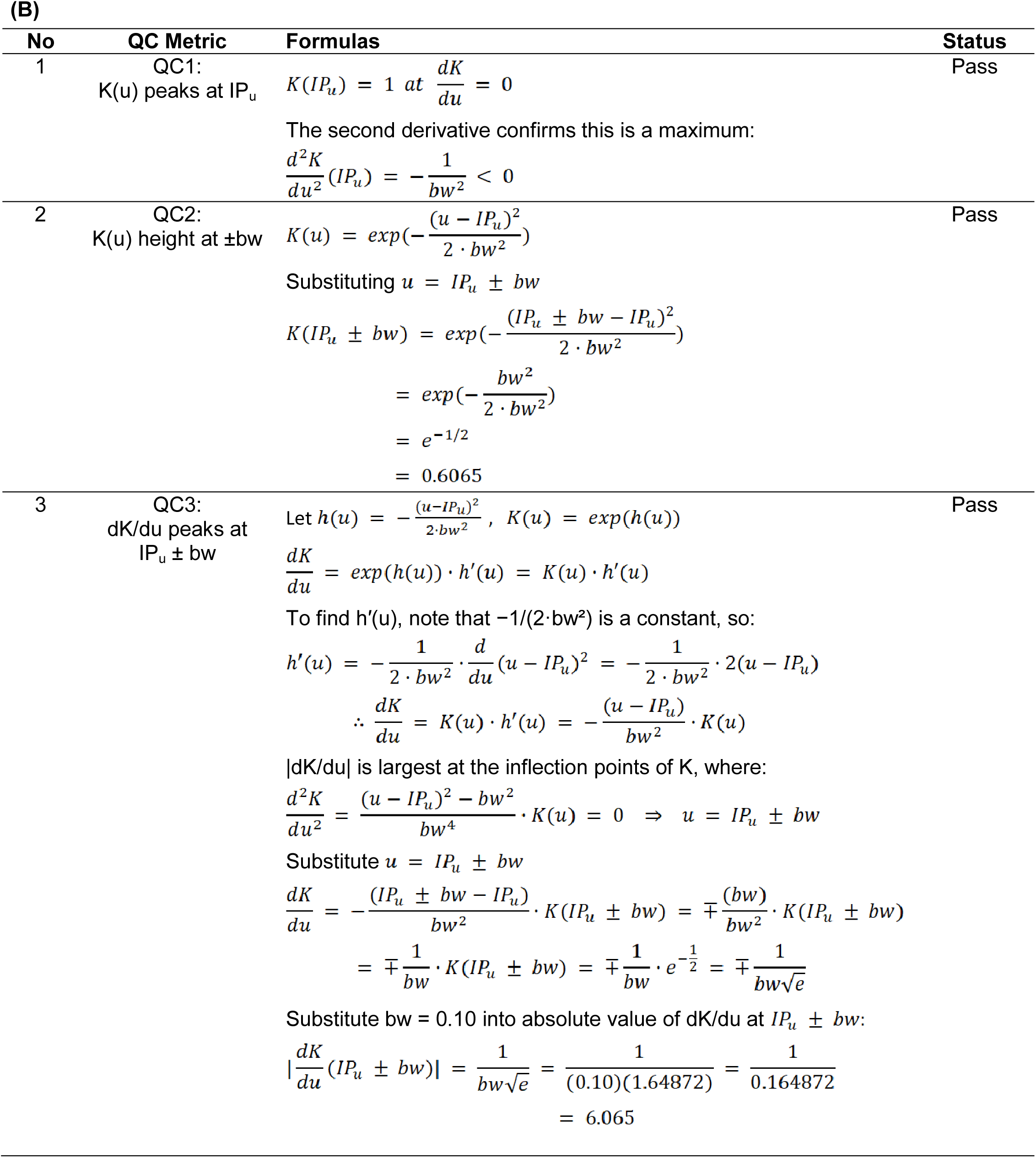
Kernel of the CMRF adaptive band. (A) The Gaussian kernel K(u) sets how much wider δ_adapt_ is than δ_base_ at each u. QC1 checks that the band is widest at the IP: K reaches its peak of 1 at IP_u_, hence δ_adapt_ reaches δ_max_ there. QC2 checks the width one bandwidth away: K(IP_u_ ± bw) = exp(-1/2) = 0.6065, thus 60.65% of the extra width remains. QC3 checks where the band narrows fastest: |dK/du| is largest at IP_u_ ± bw, at 1/(bw√e) = 6.065. (B) Derivations of QC1 to QC3.

**Supplementary Figure S12.**
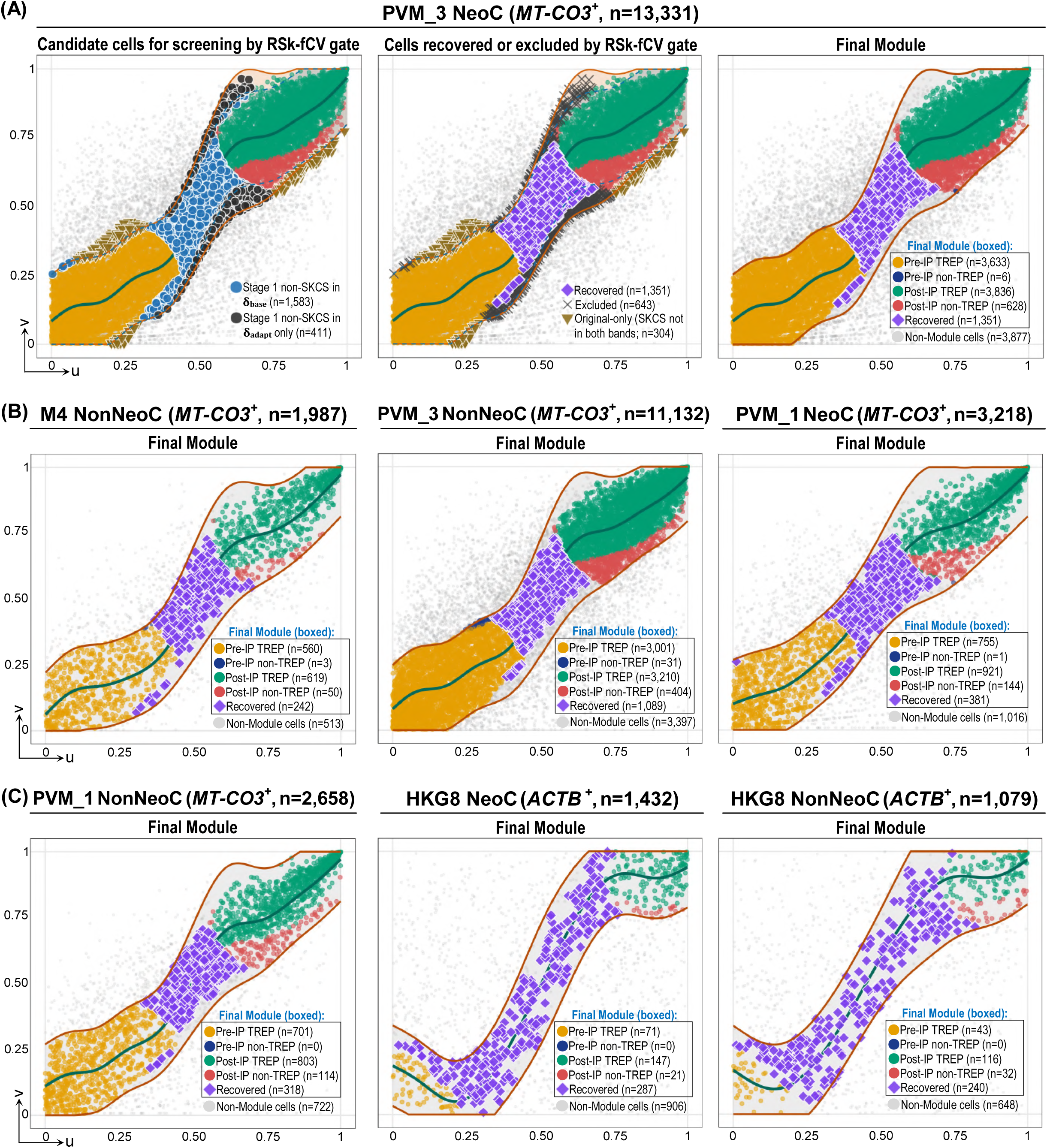
CMRF in the other arms and region sets (as in Figure 6). (A) PVM_3 NeoC: Stage 1 candidates, RSk-fCV outcome, and final module. (B, C) Final modules of the other combinations. In HKG8, the spline (dashed) is not monotone, and Recovered cells outnumber OriRef cells (Supplementary Table S10).

**Supplementary Figure S13.**
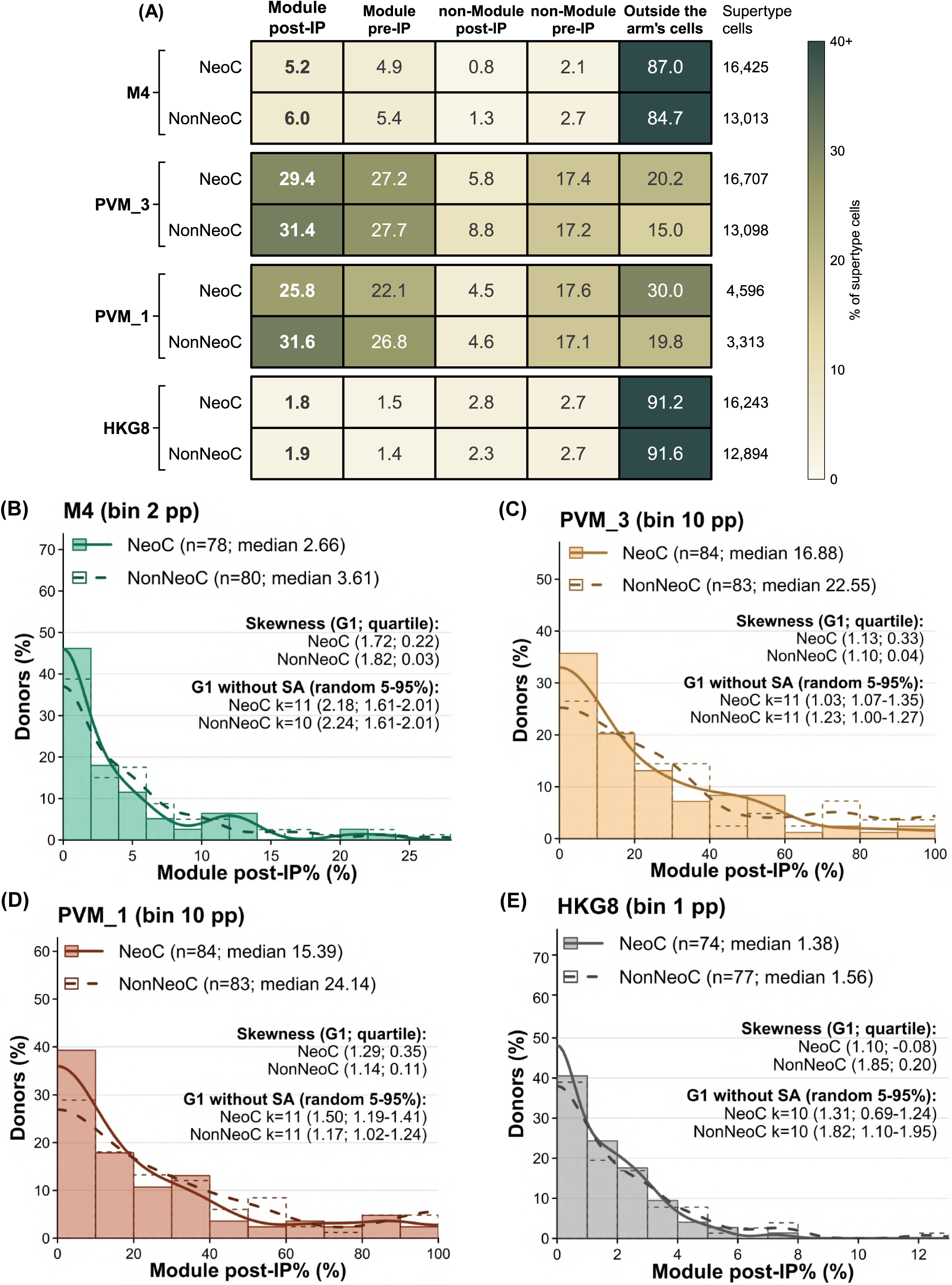
Module post-IP% and the cell groups it was drawn from. (A) Cell groups as a percentage of each arm’s supertype cells, pooled over donors: Module post-IP, Module pre-IP, non-Module post-IP, non-Module pre-IP, and supertype cells outside the arm, summing to 100%. (B to E) Module post-IP% over donors for M4, PVM_3, PVM_1, and HKG8 (NeoC filled, NonNeoC outlined), with kernel densities. Skewness: G1 (above 1: strongly right-skewed) and quartile skewness (near 0: symmetric). Each panel also yields G1 without the k SA donors, and the range of G1 (5th-95th percentile) when k other High-ADNC donors were removed at random 1,000 times.

**Supplementary Figure S14.**
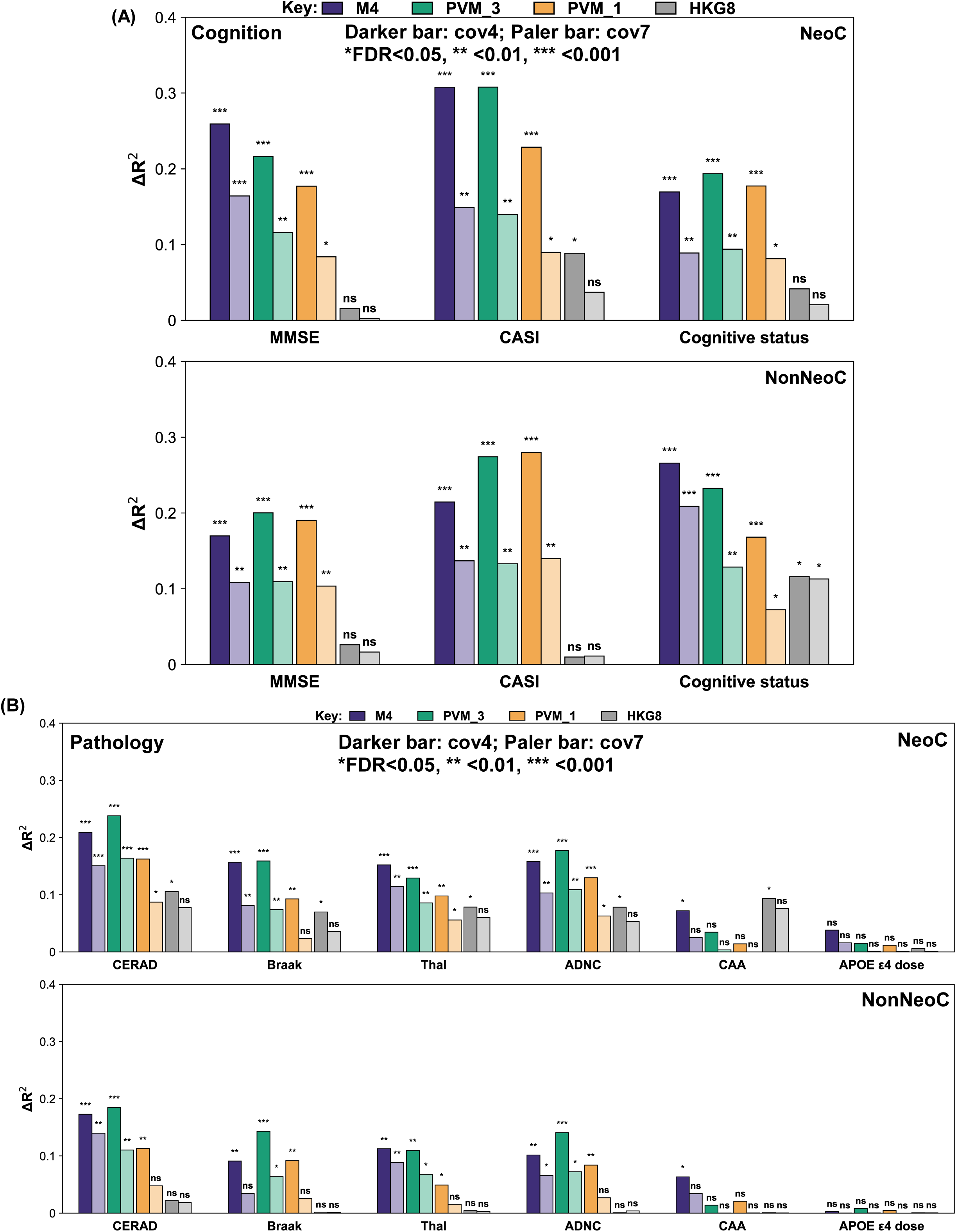
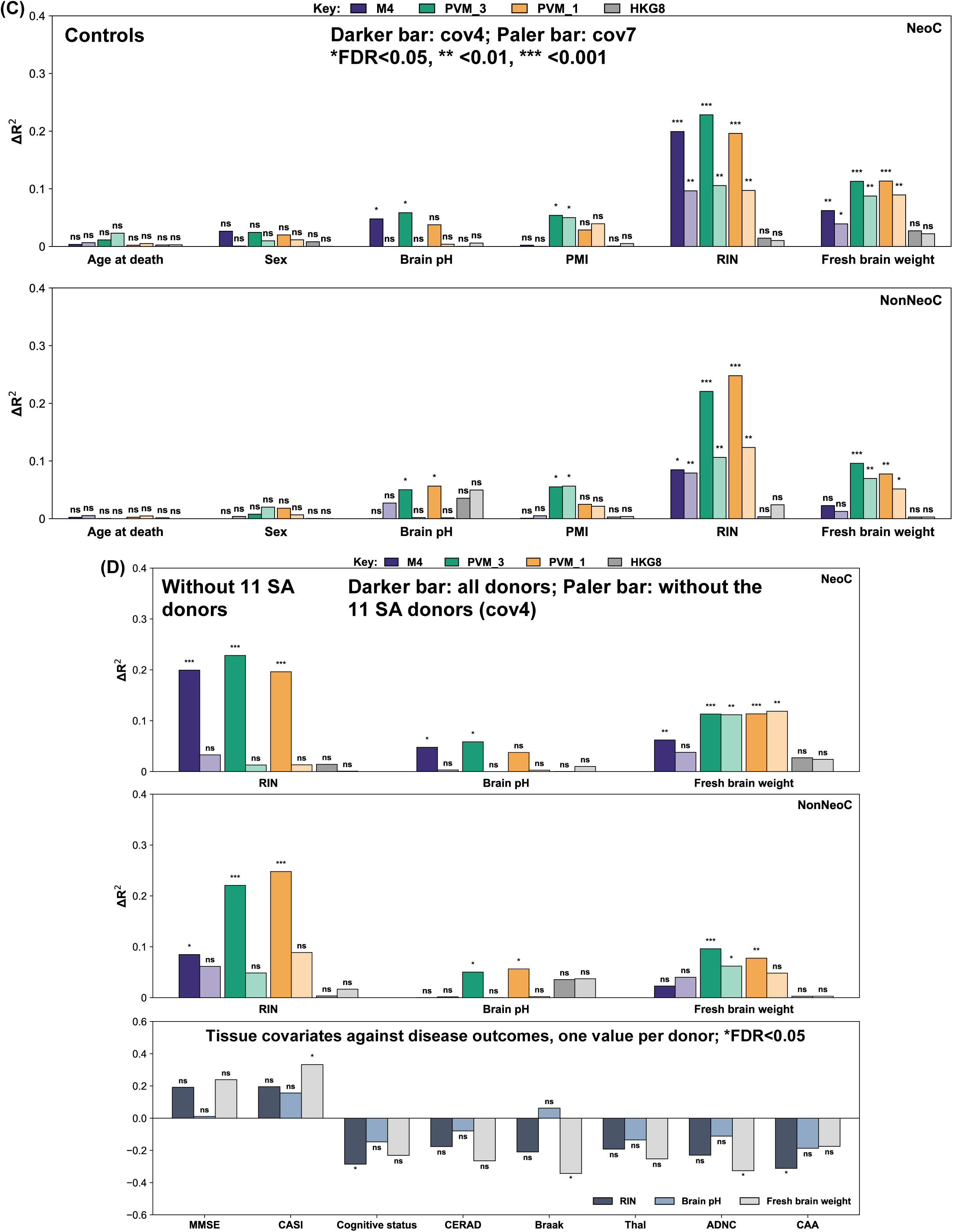
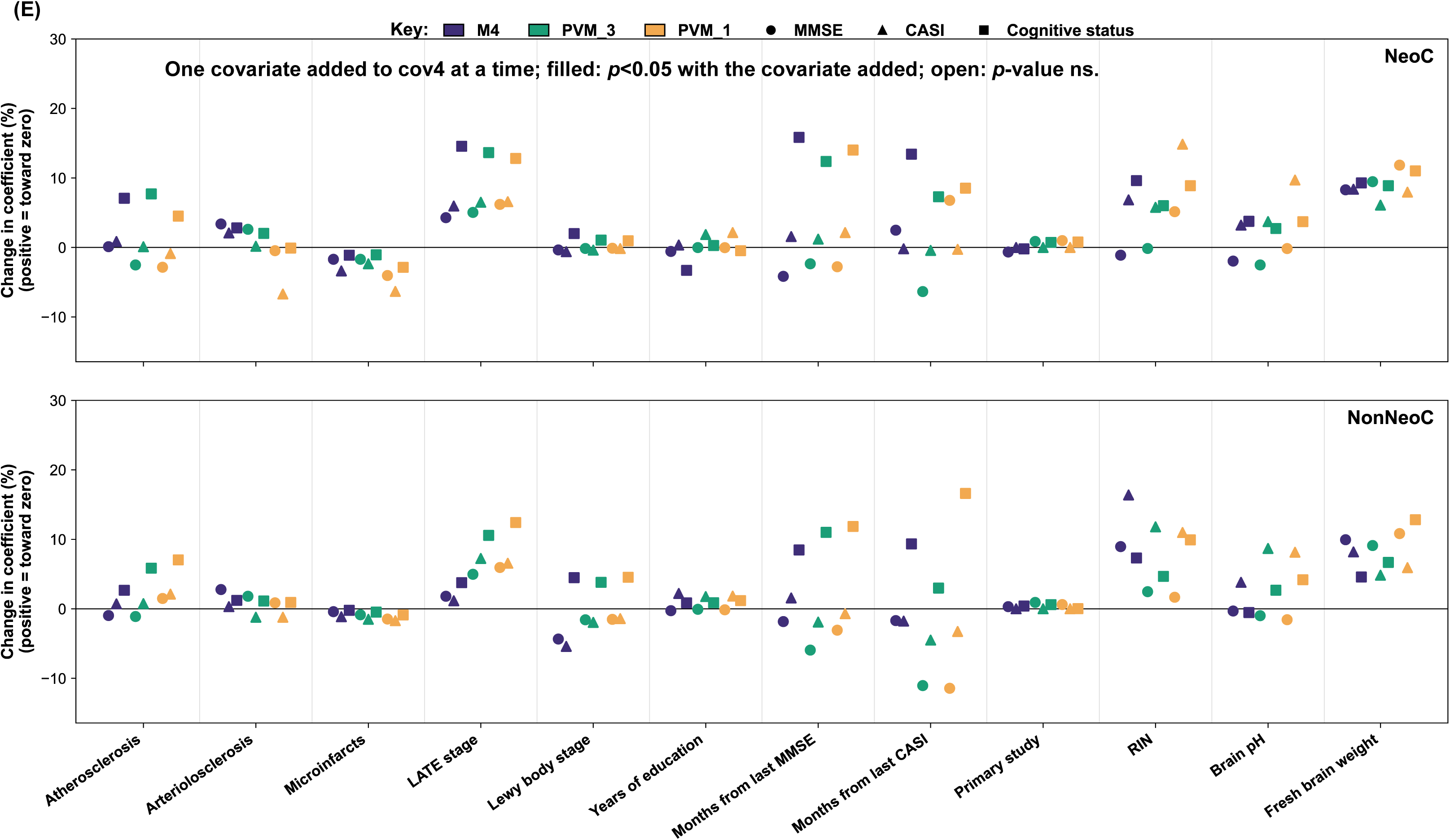
Module post-IP% against tissue quality and added covariates. (A to C) ΔR^2^ of Module post-IP% at cov4 (dark) and cov7 (pale; cov4 plus RIN, brain pH, and fresh brain weight) for cognition (A), pathology and APOE ε4 dose (B), and negative-control outcomes (C). (D) Top, ΔR^2^ of Module post-IP% on these three measures with and without the 11 SA donors (dark and pale). Bottom, their r_s_ with eight disease outcomes. (E) Change in the cognition coefficient (positive, toward zero) when each of 12 covariates was added to cov4; filled symbols indicate *p*<0.05 with the covariate added. In A to D, * marks FDR<0.05, ** FDR<0.01, *** FDR<0.001 (Supplementary Table S12).

**Supplementary Figure S15.**
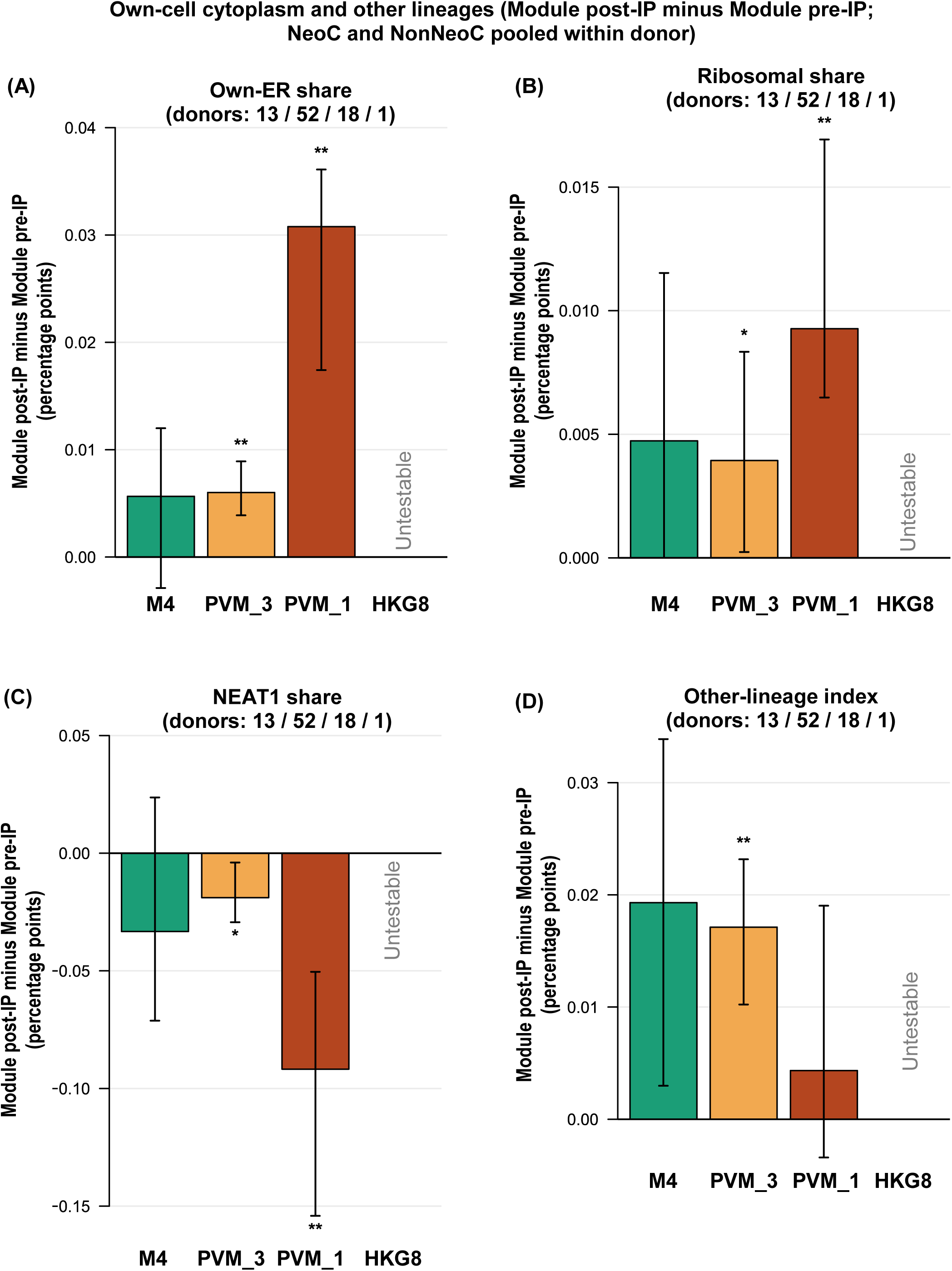
Own-cell cytoplasm and other lineages. Module post-IP minus Module pre-IP cells of the same donor (NeoC and NonNeoC pooled): (A) own-endoplasmic reticulum (ER), (B) ribosomal, and (C) *NEAT1* shares of non-mitochondrial UMIs; (D) other-lineage index, the percentage of total UMIs on five non-microglial marker panels. Bars are the median over donors of the within-donor difference, in percentage points, with 95% CIs and FDR; * FDR<0.05, ** FDR<0.01. Donors with at least 20 cells in both groups are given in each panel title (M4, PVM_3, PVM_1, HKG8); HKG8 is below the 10-donor floor. Values are in Supplementary Table S13.

**Supplementary Figure S16.**
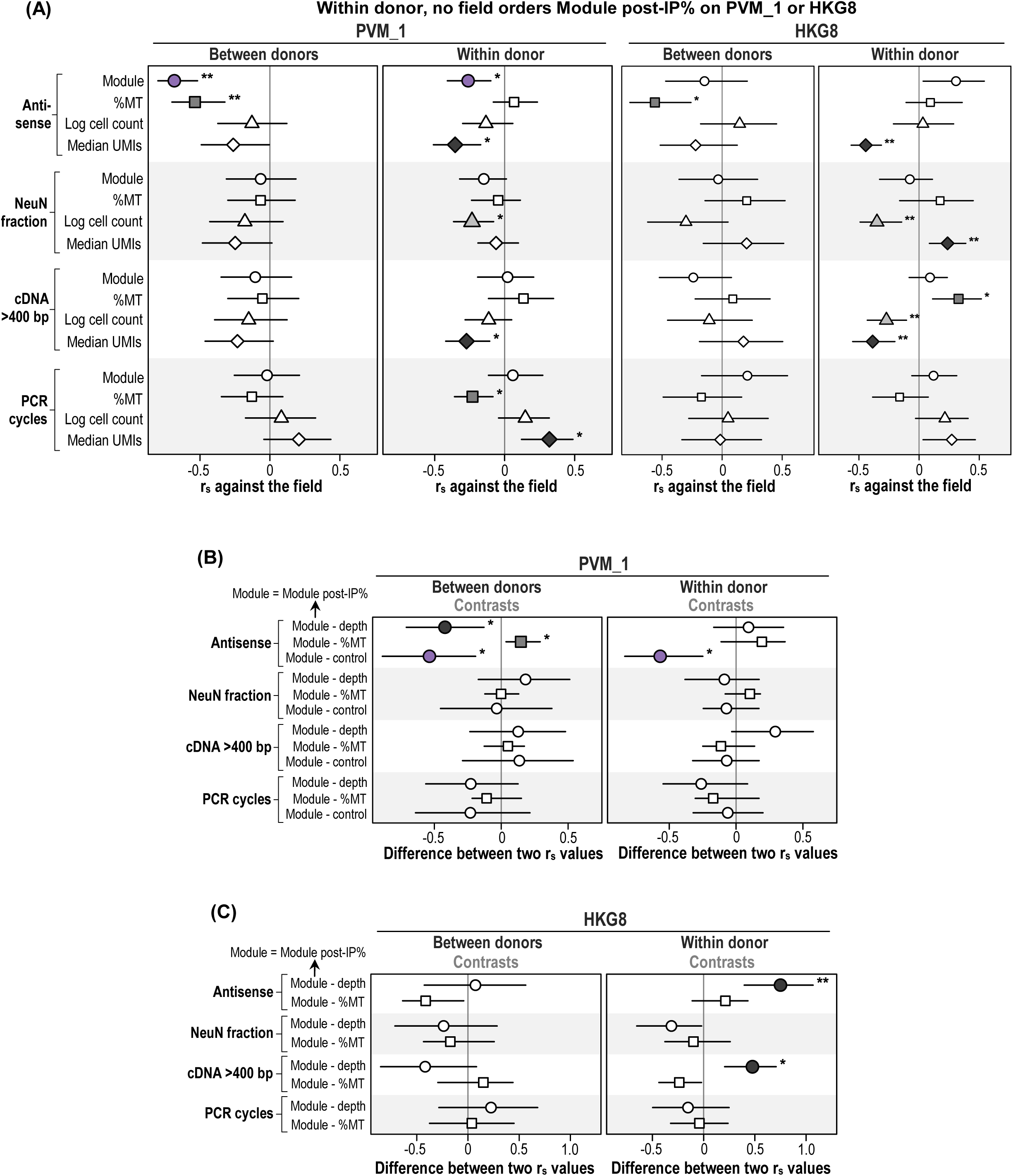
Library fields in PVM_1 and HKG8 (as in Figure 13G). (A) r_s_ of Module post-IP%, %MT, log cell count, and log median UMIs with each library field in PVM_1 and HKG8, between donors (donor means) and within donor (units centered on the donor mean, in donors with at least three units). (B) Differences between two r_s_ values on the same donor resamples in PVM_1: Module minus depth, |Module r_s_| minus |%MT r_s_|, and Module minus control (PVM_1 minus HKG8). (C) As panel B for HKG8, which is the control and thus has no Module minus control contrast. Filled symbols and * mark FDR<0.05, ** FDR<0.01. Values are in Supplementary Table S14.

**Supplementary Figure S17.**
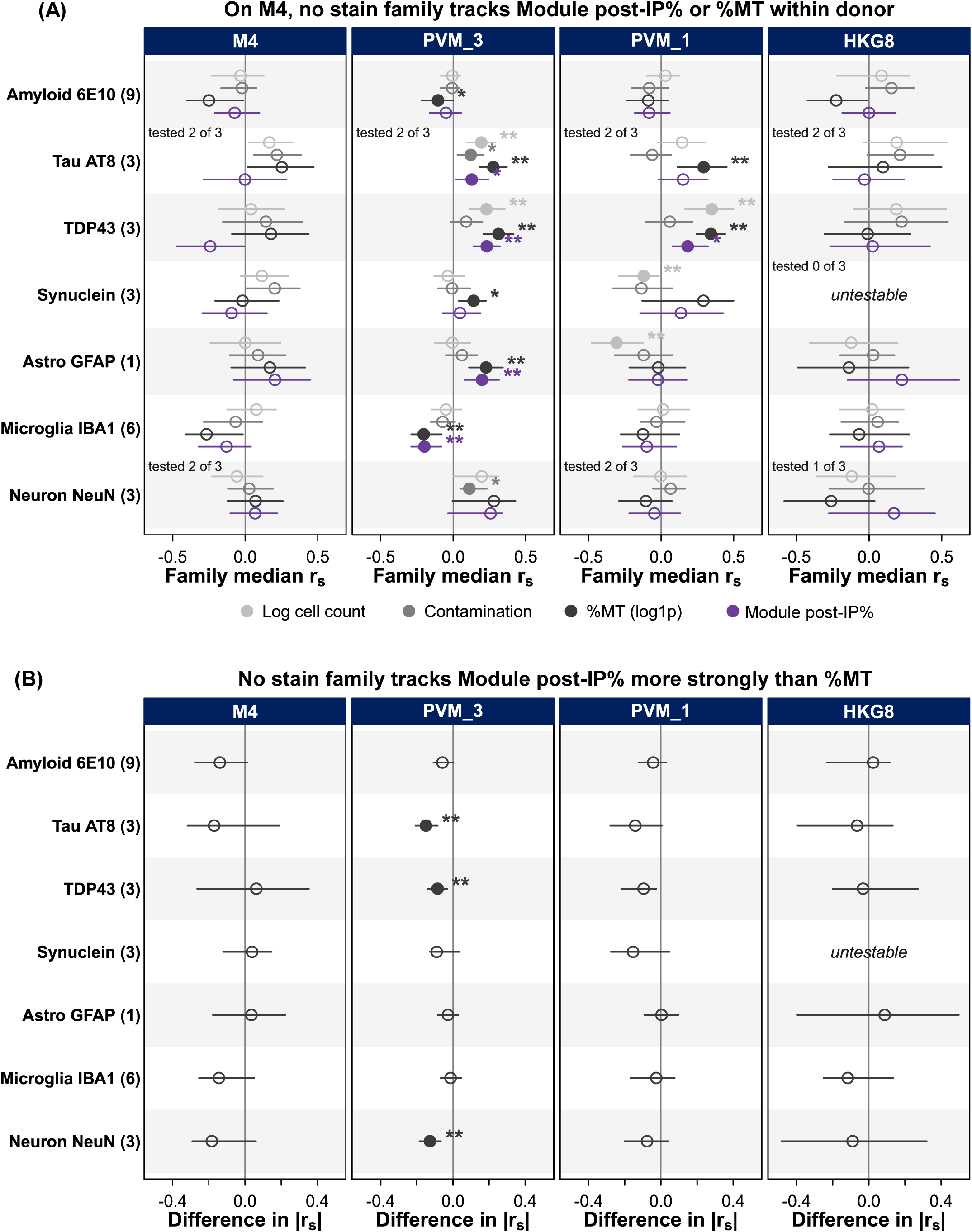
Stain families and each quantity within donor. Units and levels as in Figure 14A; each family label gives its number of measures. (A) Family median of the within-donor r_s_ of its measures with log cell count (light gray), DecontX contamination (gray), %MT (log1p; black), and Module post-IP% (purple). “Tested n of m”: measures above the floor (10 donors with at least three units). (B) Family median, over measures tested for both, of |Module r_s_| minus |%MT r_s_|; negative values mean %MT tracks the family more strongly. BH within arm (and quantity in panel A) over the seven families. Filled symbols and * mark FDR<0.05, ** FDR<0.01. Values are in Supplementary Table S15.

**Supplementary Figure S18.**
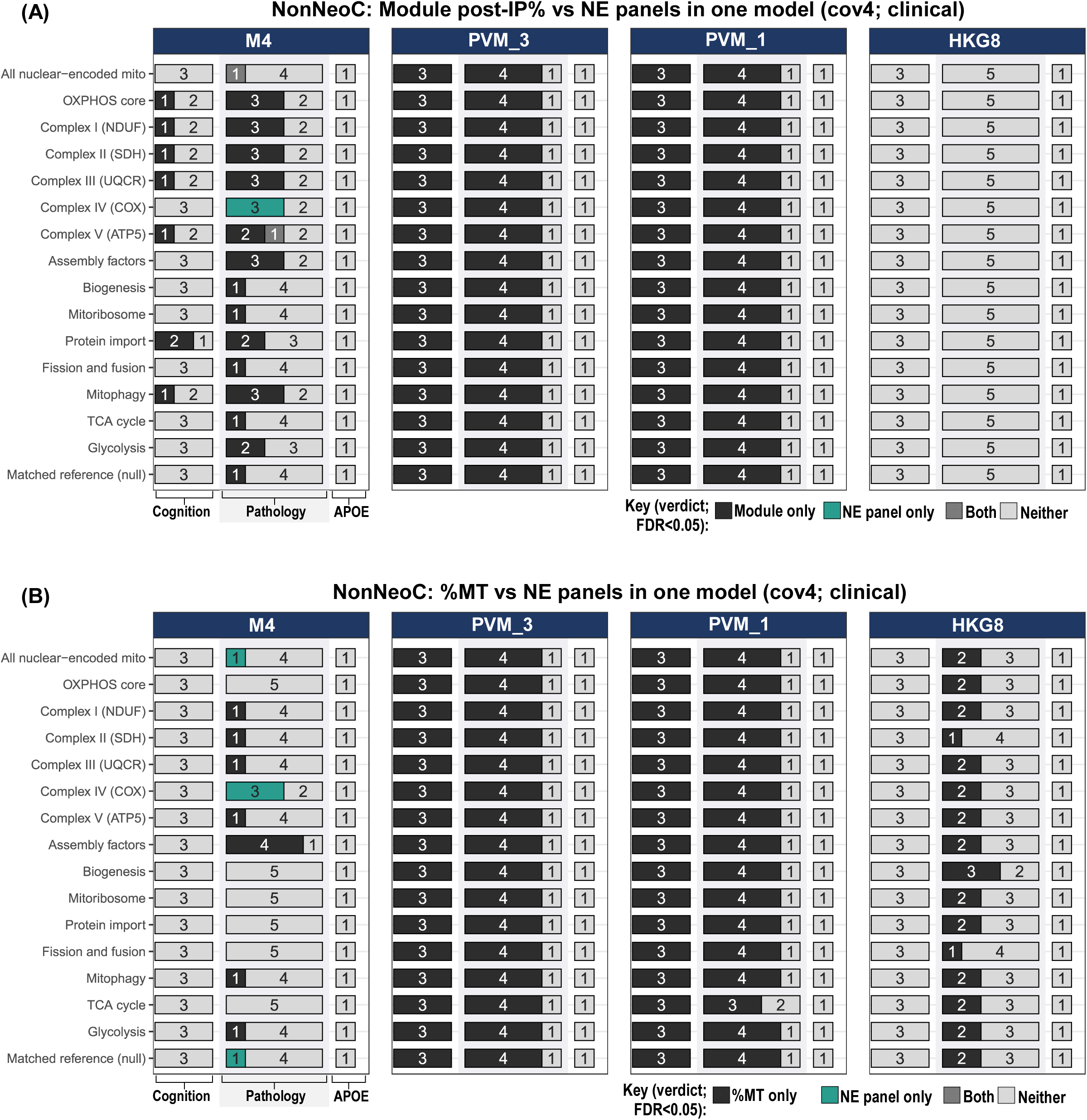
Nuclear-encoded panels on NonNeoC. As Figure 15A, for NonNeoC: (A) Module post-IP% and (B) %MT, each with every nuclear-encoded (NE) panel and the matched reference in one cov4 OLS model per outcome. Bars count the outcomes with each verdict in each group; verdicts as in Figure 15A. Values are in Supplementary Table S16.

**Supplementary Figure S19.**
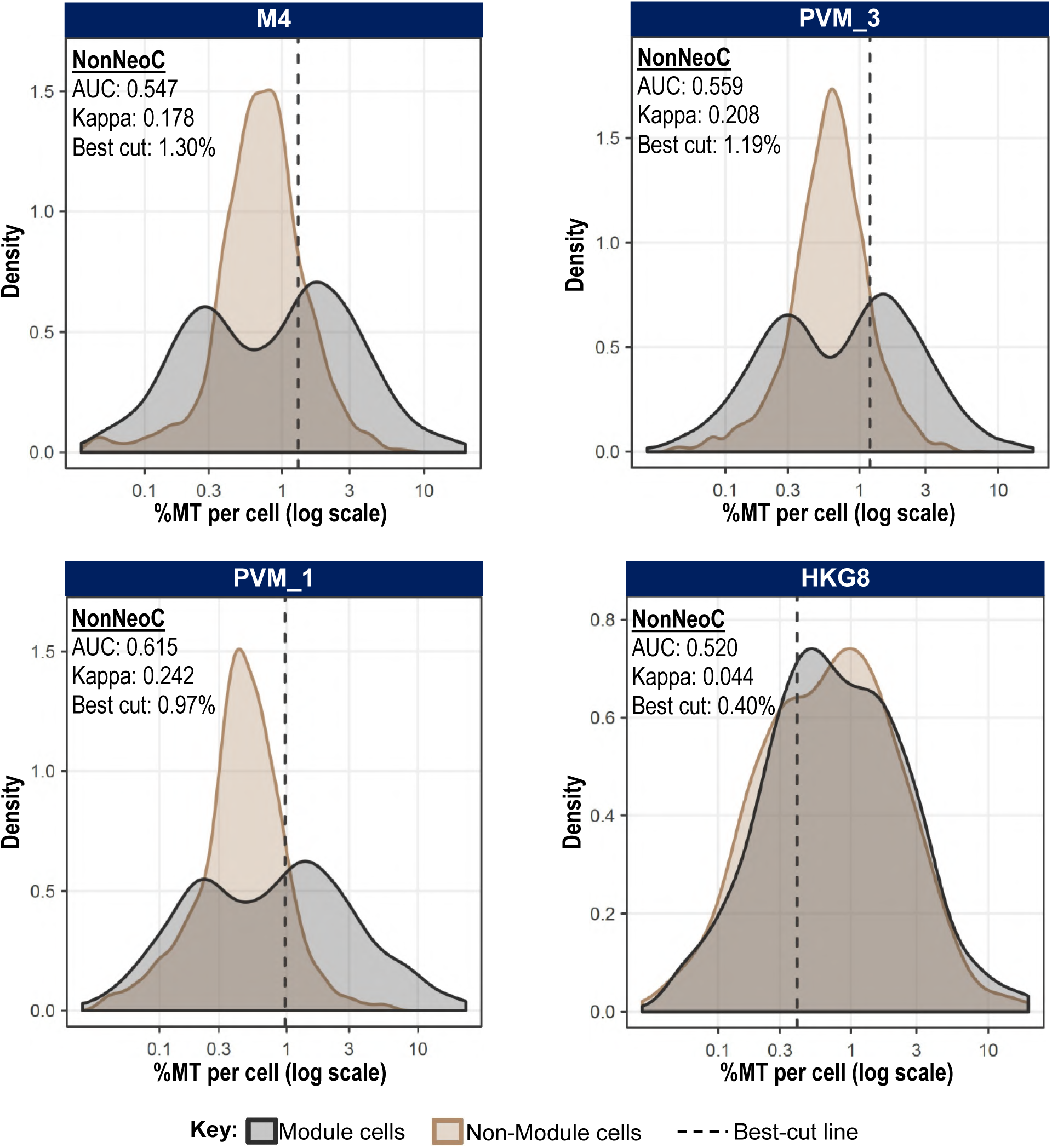
Per-cell %MT of Module and non-Module cells on NonNeoC (as in Figure 15B). Kernel densities pooled over donors, the best threshold (dashed line: Youden’s J), the AUC, and Cohen’s κ at that cut. Values are in Supplementary Table S16.

**Supplementary Figure S20.**
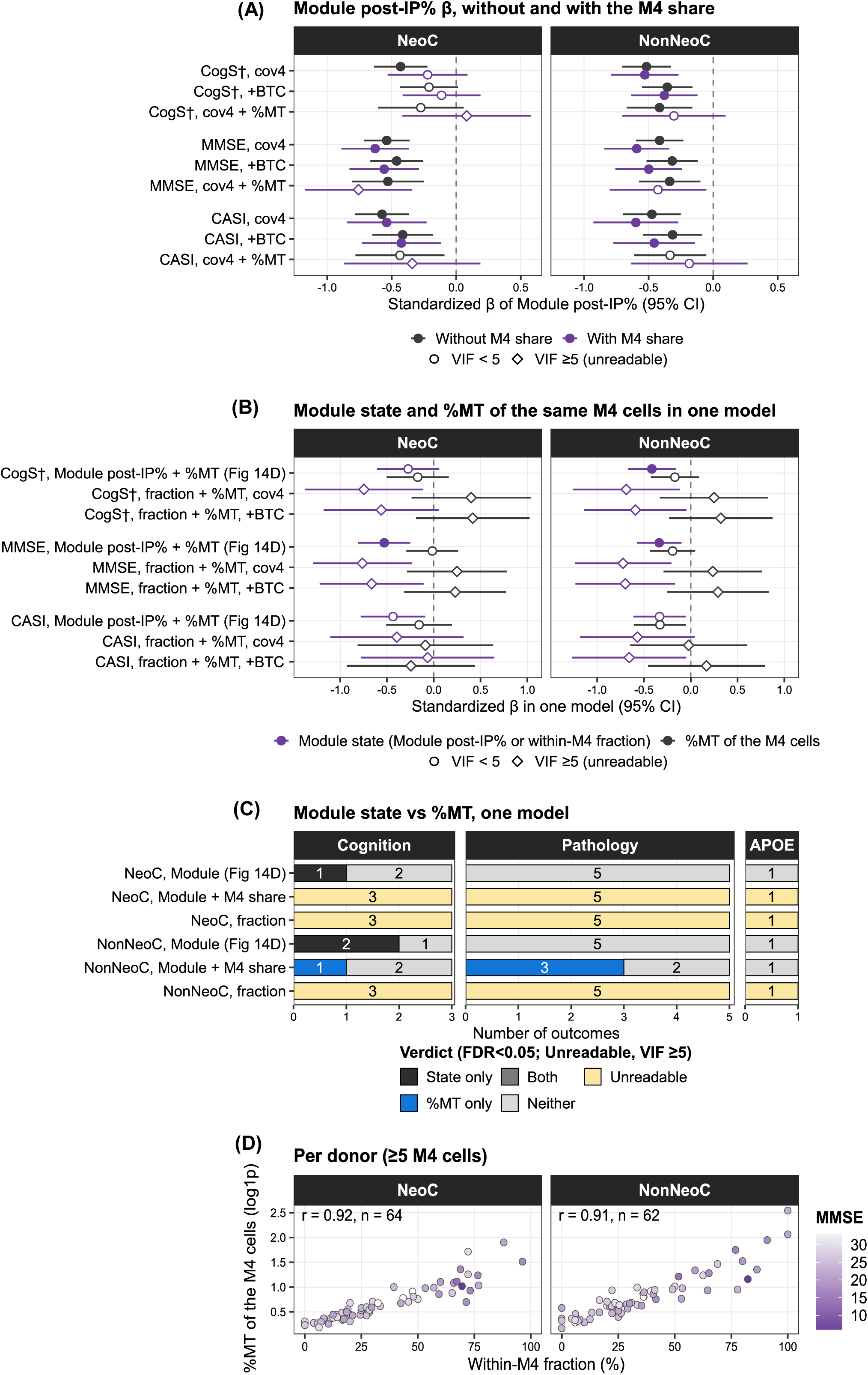
M4 share and Module state against %MT in M4. %MT as in Figure 14D; unreadable: VIF≥5; ^†^sign-reversed, hence a lower β means worse cognition. (A) β of Module post-IP% on cognition without (dark) and with (violet) the M4 share, at cov4, +BTC, and cov4 + %MT; bars: 95% CIs; filled symbols mark FDR<0.05; diamonds: unreadable. (B) Module post-IP% or the within-M4 fraction with %MT in one model. (C) Verdicts for B and for the Figure 14D model with the M4 share; State only, Module post-IP% or fraction only. (D) Within-M4 fraction against %MT per donor, shaded by MMSE (Supplementary Table S18).

**Supplementary Figure S21.**
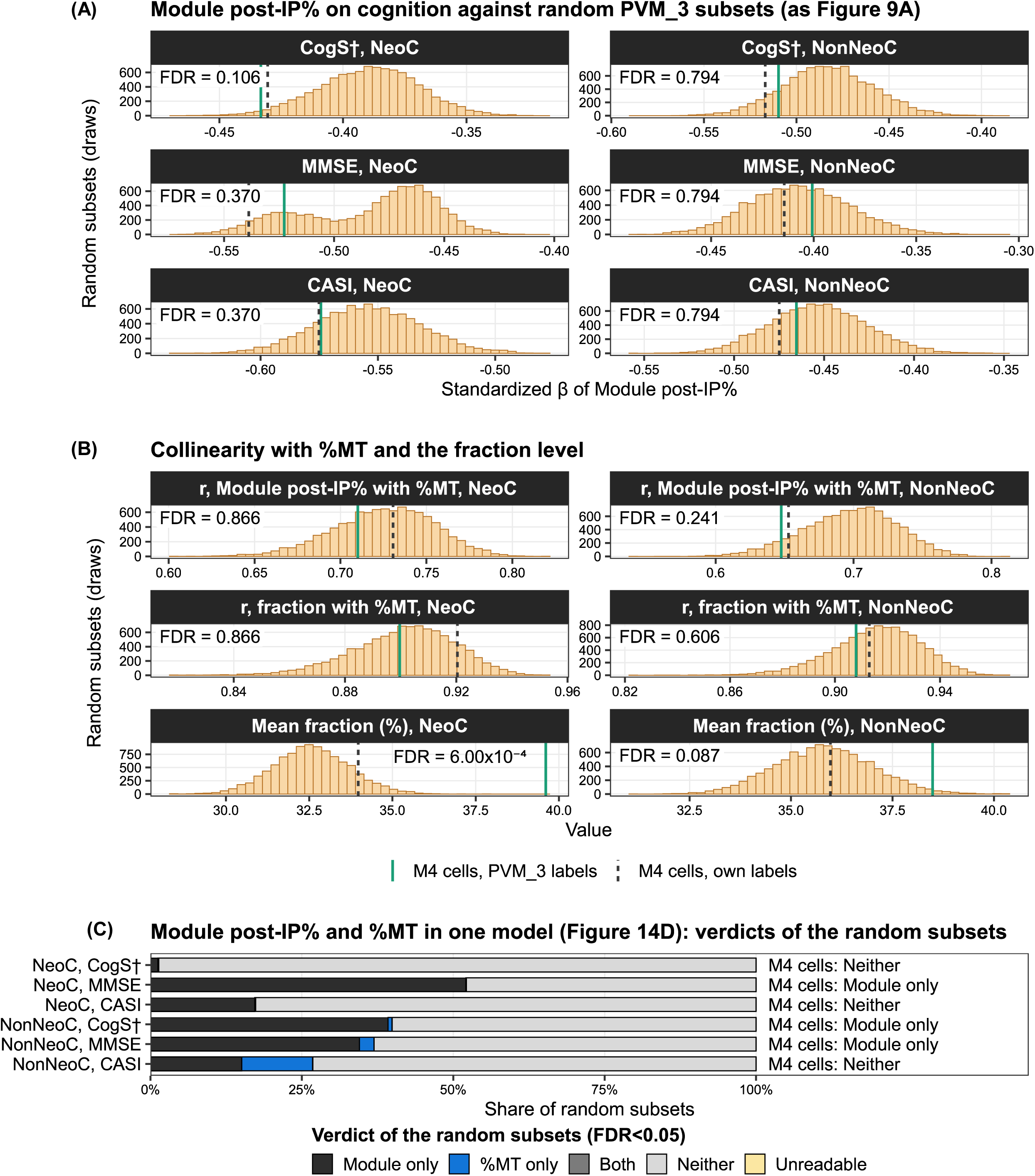
M4 cells against random PVM_3 cells of the same number. Per donor and region, random PVM_3 cells matched the M4 cell number (10,000 draws per region set), with PVM_3 labels and cov4 models. Histograms, random subsets; solid green line, M4 cells; dashed line, M4 cells under their own labels (not tested). FDR, from two-sided empirical *p*-values. (A) β of Module post-IP% on cognition; ^†^higher β means worse cognition. (B) Pearson r of Module post-IP% and the fraction with %MT, and mean fraction. (C) Share of random subsets per Figure 14D verdict, and the M4 verdict (Supplementary Table S18).

## Supplementary table legends

**Supplementary Table S1.** Cell selection: selection-rule verdicts at 101 cutoffs, the cutoff searches and sweep, region-set matching, scorer comparison, and M4 composition by supertype with donor-bootstrap 95% CIs (Figure 1A and 1B; Supplementary Figure S1A).

**Supplementary Table S2.** Mic.13 validation: differential expression of M4-selected against non-selected PVM_3 cells, recovery of the held-out Mic.13 markers, and per-gene Mic.13 detail.

**Supplementary Table S3.** Choice of *MT-CO3*: the 13 mitochondrial genes compared on nine screen measures, their composite rank with leave-one-measure-out ranks, and the DFG partners of every gene in the screens (Figure 1C, 1D, and 1F).

**Supplementary Table S4.** GOI-DFG screens: populations, subsample sizes, DFG counts, and top genes of the regional screens (Figure 1F), and the pooled screens of each supertype with the rank of MT-CO3 (Figure 1E).

**Supplementary Table S5.** Pre-checks of the immune object: the mitochondrial gene inventory, module r_s_ within donor and by library, ambient-index quartile, region, and supertype, the abundance-matched null, break-location screens, pathology fits, and checks of the clinical variables (Figure 1E and 1G; Supplementary Figure S1B to S1F).

**Supplementary Table S6.** DecontX ambient-RNA removal: the contamination estimate of every immune cell, summarized by supertype, region, and region set, with module and reference gene expression and module r_s_ before and after decontamination.

**Supplementary Table S7.** Cells of the four arms, GAM fits, the S_i_ floor, and the pitcher plots: every GOI-positive cell with expression (log1p CP10K), scores, labels, and copula coordinates, plus GAM fits, the M4 S_i_ floor, and pooled r_s_, *τ*_b_, and SKCS-core (Figures 1H and 2 to 4).

**Supplementary Table S8.** Inflection points. (A) IP detection for each arm and region set: IP_x_ with its 95% CI, IP_y_, the test behind C2, the IPRS components, IPRS, and tier (Supplementary Figure S6; Figure 3; Supplementary Figure S7). (B) R(x) and its 95% band at every tenth point of the 2,000-point grid (Supplementary Figure S6). Cell-level values, including the pre-IP and post-IP groups, are in Supplementary Table S7, sheet A.

**Supplementary Table S9.** Copula coordinates: the IP and the S_i_ and K_i_ gates converted to copula coordinates, and the worked cells A to P with their values, ranks, and copula coordinates (Figure 4; Supplementary Figures S8 to S10).

**Supplementary Table S10.** CMRF and validation of the Recovered cells: every GOI-positive cell with its band, CMRF class, and coordinates, the band and gate settings, the M4 NeoC gate trajectory, and the statistical tests (Figures 5 to 8; Supplementary Figures S11 and S12).

**Supplementary Table S11.** Differential expression between Module and non-Module cells, and between post-IP and pre-IP cells: paired pseudobulk DEGs per donor (FDR<0.05 and |log_2_FC|≥0.25) in each arm and region set, their test against HKG8, the 14 non-mitochondrial post-IP genes, and UMI checks.

**Supplementary Table S12.** Clinical association, benchmark against competitors, and boundary ablation: the donor frames, the clinical, competitor, and boundary models with their statistics, the boundary scorecard, and the tissue-quality and covariate checks (Figures 9 to 11; Supplementary Figure S14).

**Supplementary Table S13.** Battery 1, contamination and doublets: the clinical models with contamination added or doublets removed, within-donor contamination by group and tertile, the Gerrits ambient-RNA analyses, depth thinning, and the Module post-IP minus pre-IP differences (Figure 12; Supplementary Figure S15).

**Supplementary Table S14.** Battery 2, measurement artifacts: the donor share of variance, split-half r_s_, CMRF label agreement, within-donor co-expression, offset share, and the library-field correlations, with the values of every donor-region unit (Figure 13; Supplementary Figure S16).

**Supplementary Table S15.** Battery 3, histology and clinical outcomes against %MT: correlations with the IBA1 measures and CPS scores, the one-model verdicts for the stain measures and clinical outcomes, the collinearity of Module post-IP% with %MT, and the stain-family summaries (Figure 14; Supplementary Figure S17).

**Supplementary Table S16.** Battery 4, nuclear-encoded panels, %MT thresholds, and donor requirement: Module post-IP% or %MT with each nuclear-encoded panel, the AUC of %MT for Module membership, and the donor and region subsampling (Figure 15; Supplementary Figures S18 and S19).

**Supplementary Table S17.** Battery 5, removal of the 11 SA donors: the Battery 1 to 4 statistics and verdicts with all donors and without the 11, their LODO changes, and the random-removal draws (Figure 16).

**Supplementary Table S18.** M4 against PVM_3: the M4 share models, verdicts, bootstrap, and correlations, the random-subset readouts with their empirical *p*-values, the donor values, and the pre-specified readings of both analyses (Supplementary Figures S20 and S21).

**Supplementary Table S19.** Summary of all tests: the test definitions and rules, the verdict of every test, arm, and region set, the tallies and boxed values of Figure 17B, every model read by the tests, and the build checks (Figure 17).

