## Supplementary Methods for "CEP-IP with CMRF in SEA-AD Microglia: A Mitochondrial Co-expression Module Linked to Cognition"

### Supplementary Methods (Figures 1-19 and Supplementary Figures S1-S21)

Supplementary Methods are organized in four parts. Settings not provided here are in the deposited scripts (Supplementary Methods A3). The sections are as follows:

**Main analysis (Figures 1 to 11)**
**A1. Data and conventions**
S1. Cell selection (Figure 1A and 1B; Supplementary Figure S1A)
S2. Choice of supertypes and the GOI-DFG screens (Figure 1C to 1F)
S3. Gene-level and technical controls, and the four arms (Figure 1G and 1H; Supplementary Figure S1B to S1F)
S4. Cell-level scores and the SKCS-core (Figure 2; Supplementary Figures S2 to S5)
S5. Inflection point (IP) and the pre-IP and post-IP groups (Figure 3; Supplementary Figures S6 and S7)
S6. Copula coordinates (Figure 4; Supplementary Figures S8 to S10)
S7. CMRF and validation of the Recovered cells (Figures 5 to 8; Supplementary Figures S11 to S13)
S8. Differential expression between Module and non-Module cells (Supplementary Table S11)
S9. Clinical association, benchmark against competitors, and boundary ablation (Figures 9 to 11; Supplementary Table S12)

**Batteries 1-5 and summary of all tests (Figures 12 to 17)**
**A2. Settings shared by the Batteries**
S10. Battery 1 (Figure 12 and Supplementary Figure S15)
S11. Battery 2 (Figure 13 and Supplementary Figure S16)
S12. Battery 3 (Figure 14 and Supplementary Figure S17)
S13. Battery 4 (Figure 15 and Supplementary Figures S18 and S19)
S14. Battery 5 (Figure 16)
S15. M4 against PVM_3 and the summary of all tests (Figure 17; Supplementary Figures S20 and S21)

**External validation (Figures 18 and 19)**
S16. External validation in two independent datasets (Figures 18 and 19; Supplementary Tables S20 and S21)

**Code availability**
**A3. Code availability**

#### Supplementary Methods A1. Data and conventions

All analyses used the SEA-AD immune single-nucleus RNA-seq multiregion release of 2026-06-22 (240,651 nuclei, 36,601 genes, 84 donors, and 10 regions) [1, 2]. Expression was the released log1p counts per 10,000 (CP10K). As in the main text, nuclei are referred to as cells, and supertype names drop the "Micro-" prefix and the "-SEAAD" suffix for brevity. The two region sets were analyzed separately unless indicated otherwise: NeoC (134,088 nuclei) and NonNeoC (106,563 nuclei).

Ranks were computed with averaged ties. Generalized additive models (GAMs) used a thin-plate regression spline (TPRS) with a shrinkage penalty and were fitted by restricted maximum likelihood (REML) with $\gamma=1.5$. The donor bootstrap resampled donors with all their cells or units and yielded percentile 95% confidence intervals (CIs). Multiple testing was corrected by the Benjamini-Hochberg (BH) procedure within the families stated for each analysis. Unless stated otherwise, two groups were compared by a t-test (paired t-test for paired data) if both passed the Shapiro-Wilk test, and otherwise by a Mann-Whitney U test. Each supplementary table was built by a script that checked its values against the analysis outputs.

#### Supplementary Methods S1. Cell selection (Figure 1A and 1B; Supplementary Figure S1A)

Mic.13-like cells were selected in each region set with 133 Mic.13 marker genes (85 upregulated, 48 downregulated; no mitochondrial gene) and their published fold changes and adjusted *p*-values [3].

Six candidate scorers were compared, i.e., M1-M6. Initially, three scores were computed per cell: M1 (each marker weighted by its fold change; FC), M2 (by its FC x [$-\log_{10}$ FDR]), and M3 (by the rank of its FDR). M4-M6 were intersection scorers that selected only the cells in the top 3.0% (or the cutoff tested) of two of these scores, where M4 required both M1 and M2 (i.e., M1 ∩ M2), M5 required both M1 and M3 (i.e., M1 ∩ M3), and M6 required both M2 and M3 (i.e., M2 ∩ M3).
These six scorers were compared on the composition of the selected cells and on the donor-level clinical correlations ($r_s$) of the selected PVM_3 cells (Supplementary Table S1). M3 failed the selection rule in both region sets. Over five cutoffs (2.5%-5%) in both region sets, M4 had the highest average rank of the six scorers in FDR-significant clinical associations and the highest mean $r_s$ of the three intersection scorers. M4 was therefore chosen as the scorer. For validation on the 68 Mic.13 markers outside the scoring panel, M4-selected and non-selected PVM_3 cells were compared per region (FDR<0.05, $|\log_2 FC| \geq 0.25$, and expression in ≥15% of selected cells). Recovery of these markers was tested via hypergeometric test (Supplementary Table S2).

**Supplementary Methods S2. Choice of supertypes and the GOI-DFG screens (Figure 1C to 1F)**
cDFG was the mean log1p CP10K of the eight DFGs. Module $r_s$ was the Spearman correlation of *MT-CO3* with cDFG over cells with *MT-CO3*>0 (24,463 of 29,805 PVM_3 cells), as the ranks of a gene with many zeros are dominated by ties. In a GOI-DFG screen, a gene passed the GOI gate when it was expressed in ≥25% of cells with a mean ≥0.3 among expressing cells. Genes expressed in ≥10 cells were candidate partners, and a partner was a DFG when it reached $r_s \geq 0.6$ and $\tau_b \geq 0.5$ with the GOI with the same sign.

The pooled GOI-DFG screen used all cells of each supertype in the region set from donors with ≥5 cells, and subsampled to 2,500 cells with equal numbers per region. The regional screen used M4 cells of each region, and the Module DFGs were the DFG partners of *MT-CO3* in ≥8 of the 10 regions. *MT-CO3* was compared with the other 12 mitochondrial genes by its mean rank over nine expression and screen measures (Supplementary Table S3). PVM_1 met the supertype criteria of Section 2.2 (Figure 1E) and became the supertype control, with the donor floor of these criteria adopted from the power estimate of Figure 15D.

**Supplementary Methods S3. Gene-level and technical controls, and the four arms (Figure 1G and 1H; Supplementary Figure S1B to S1F)**
Over the PVM_3 cells with *MT-CO3*>0, the $r_s$ of each of the 12 other mitochondrial genes with *MT-CO3* was computed. These $r_s$ values were then regressed on the genes' $\log_{10}$ mean counts to yield an abundance trend, and *ACTB* was placed against this trend by its scaled residual (Figure 1G). In M4 cells, cDFG and cHKG (the mean of nine housekeeping genes) were each fitted on *MT-CO3* by GAM, and their deviance explained (DE) was compared (Figure 1H). Module $r_s$ was also compared with 295 random nine-gene panels matched on abundance and detection (Supplementary Figure S1B).

Module $r_s$ was correlated with the intronic:exonic ratio per library (185 libraries), directly and after partialing out the library median %MT (Supplementary Figure S1C and S1D). Per nucleus, cells were divided into four groups by their ambient index and module $r_s$ was computed within each group (Supplementary Figure S1E). In leave-one-region-out, module $r_s$ was recomputed over the cells of the remaining regions of the region set, once for each region left out (Supplementary Figure S1F).

DecontX [4] was run in reduced mode without empty droplets, because empty-droplet correction requires the raw Cell Ranger outputs (controlled access) and was not performed in this study. Contamination was estimated per library batch in 897 of 905 batches. A change in module $r_s$ of <0.05, set before the run, was interpreted as the module surviving decontamination (Supplementary Table S6).

Four arms were exported for the R analyses: M4 (2,130 NeoC and 1,987 NonNeoC cells), PVM_3 (13,331 and 11,132), PVM_1 (3,218 and 2,658), and HKG8 (1,432 and 1,079). Each export held the expression data, the metadata, and a settings record that every R script checked before running.

**Supplementary Methods S4. Cell-level scores and the SKCS-core (Figure 2; Supplementary Figures S2 to S5)**

In each arm, cells with GOI>0 were analyzed per region and pooled per region set, and the inflection point (IP) and later steps used the pooled sets. The heatmaps (Supplementary Figures S2 and S3) order cells by region and increasing GOI, with per-gene z-scores capped at -1 and 2. The composite was fitted on the GOI by a GAM with a linear term, a TPRS, and the donor random intercept. The basis dimension k was chosen from 3 to 10 by the lowest penalized residual sum of squares.

The explanatory power (EP) of each cell i was $EP_i = 1 - (y_i - f(x_i))^2/(y_i - \bar{y})^2$, the share of its deviation from the mean $\bar{y}$ that the population-level GAM fit f (donor term excluded) explained. Cells were ranked by EP, and the top-ranked explanatory power (TREP) cells were the top DE%, round(n x $DE_{pop}$) cells, with $DE_{pop}$ the weighted DE of the same fit [5]. All cells and the SKCS-core were fitted and labeled separately. The housekeeping pitcher plots (Figure 2B and 2D) used HKG8 cells expressing *ACTB*, *YWHAZ*, or *RAB7A*, with that gene as the GOI and the mean of the other eight housekeeping genes as the composite.

**Supplementary Methods S5. Inflection point (IP) and the pre-IP and post-IP groups (Figure 3; Supplementary Figures S6 and S7)**

The IP was located on all cells with GOI>0, as the SKCS-core has few cells near the IP. The TREP residuals (donor term excluded) were smoothed over the GOI by a residual GAM, R(x), with a donor random intercept and k = min(20, distinct GOI values - 1), and evaluated on 2,000 GOI values. $IP_x$ was the first negative-to-positive crossing of R(x). Its 95% CI was bounded by the nearest crossings of R(x)±1.96 SE, and the $IP_y$ bounds are the fitted composite values at the $IP_x$ bounds. The IP reliability score (IPRS) was the mean of five components, with tiers of ≥0.80 (very strong), ≥0.60 (strong), ≥0.40 (moderate), ≥0.20 (weak), and <0.20 (failed) [5].

**Supplementary Methods S6. Copula coordinates (Figure 4; Supplementary Figures S8 to S10)**

Thresholds were converted to copula coordinates as the number of cells at or below the threshold divided by (n + 1): $IP_y$ became $IP_v$, $IP_x$ became $IP_u$, $S_i$=0.05 became $u_{gate}$, and $K_i$=0.5 became $v_{gate}$. $IP_u$ is not drawn in Figure 4 but is the center of the CMRF band.

In the copula panels, $S_i = 4(u - 0.5)(v - 0.5)$, which equals the $S_i$ of Supplementary Methods S4 multiplied by $((n - 1)/(n + 1))^2$ (0.996 to 0.9997). The gate counts and the worked example cells used this form, on which three SKCS-core cells lay at or below $u_{gate}$ and retained their labels. The example cells (A to P) are for illustration only (Supplementary Tables S7 and S9).

**Supplementary Methods S7. CMRF and validation of the Recovered cells (Figures 5 to 8; Supplementary Figures S11 to S13)**

CMRF was implemented on all $GOI^+$ cells in the copula coordinates (u, v). In Stage 1, g(u) used the donor weights and a basis size of 3 to 10, with the donor random intercept and without a monotone constraint. The band was widened around the IP as $\delta_{adapt}(u) = \delta_{base}[1 + \kappa \exp(-(u - IP_u)^2/(2bw^2))]$, with κ=0.50 and bw=0.10 (Supplementary Figure S11). Cells with $|v - g(u)| \leq \delta_{adapt}(u)$ were admitted, and non-SKCS cells outside the band were Excluded.

In Stage 2, folds were assigned at random without blocking by donor, and the gate closed on ΔRMSE ≤BV in all eight datasets. In Stage 3, candidates removed in Stage 2 joined Excluded. Permutation tests assessed the dependence of u and v among Recovered cells (Hoeffding's D and distance correlation) and the ordered increase of |v - g(u)| from OriRef to Recovered, Original-only, and Excluded cells (Jonckheere-Terpstra test).

For Figure 8, the TREP groups of the refitted GAM held OriRef cells only. The bridge distance of a cell in ($u_{SK}$, $v_{SK}$) was the larger of its distances to the nearest pre-IP and post-IP TREP cell. Gate failure was the share of Recovered cells with $S_i \leq 0.05$ or $K_i \leq 0.5$. Monocle3 was run on the raw counts of each arm's $GOI^+$ cells (50 principal components; UMAP). Position was the projection on the axis from the pre-IP to the post-IP TREP centroid, scaled from 0 to 1. Neighbor enrichment was the share of a reference TREP group among the 50 nearest neighbors of a Recovered cell, divided by its expected share, and kBET tested the neighbor composition of 500 sampled cells.

Module post-IP% over donors was illustrated by G1 skewness and the quartile skewness (Supplementary Figure S13). G1 was also recomputed without the 11 severely affected (SA) donors, all with High ADNC, and compared with 1,000 random removals of as many other High-ADNC donors. Module post-IP% was right-skewed in every arm and region set (G1=1.1-1.9) and remained so without the SA donors, and removing the 11 SA donors lowered the skew more than random removal only in PVM_3 NeoC.

**Supplementary Methods S8. Differential expression between Module and non-Module cells (Supplementary Table S11)**

Module cells (OriRef and Recovered) and non-Module cells (Original-only and Excluded) were compared by paired pseudobulk in four comparisons per arm and region set: Module against non-Module on each side of the IP and post-IP against pre-IP within each group. Donors with ≥3 cells per group were paired, each comparison needed ≥5 paired donors, and genes detected in ≥5% of paired donors were tested. The per-donor $\log_2$FC was tested by a two-sided Wilcoxon signed-rank test with BH correction over genes within each comparison. Genes with FDR<0.05 and $|\log_2 FC| \geq 0.25$ were considered significant, and mitochondrial genes were not included as distinguishing genes.

Genes significant in PVM_3 passed when the per-donor $\log_2$FC of both M4 and PVM_1 differed from that of HKG8 at FDR<0.05 in the PVM_3 direction. A comparison was resolution-limited when it had too few paired donors for any gene to pass correction. Module membership was compared between arms on shared cells by Cohen's κ.

To test whether the passed genes reflected RNA made by the cell or carried in the nucleus, post-IP and pre-IP cells of each donor (≥3 cells per side) were compared on total and non-mitochondrial UMIs per nucleus. They were also compared on the amount and share of three panels: the passed genes that increased, 11 short neuron-specific genes, and six microglial genes (*CSF1R, P2RY12, CX3CR1, DOCK8, ARHGAP15*, and *INPP5D*). The donor $\log_2$ ratios of post-IP to pre-IP were tested against 0 by the signed-rank test (FDR over arms and region sets).

**Supplementary Methods S9. Clinical association, benchmark against competitors, and boundary ablation (Figures 9 to 11; Supplementary Table S12)**

***Clinical association***

All models were fitted at the donor level (84 NeoC and 83 NonNeoC donors), and no earlier step used a clinical variable. The denominator of each donor percentage included cells outside the arm and those with GOI=0, hence donors without cells of the arm entered with 0.

The clinical outcomes used the last MMSE and CASI scores, and the six negative-control outcomes were age at death, sex, brain pH, postmortem interval (PMI), RNA integrity number (RIN), and fresh brain weight. In each ordinary least squares (OLS) model, cov4 excluded any covariate that was the outcome, and 95% CIs were β±1.96 SE. The added blocks were +ADNC, +BTC (Braak, Thal, and CERAD), +ALL (+BTC + ADNC; sensitivity only), and +MMSE + CASI. In Figure 9D, β(Module) - β(non-Module) was tested by a two-sided t-test. For Supplementary Figure S14, cov4 was extended by the tissue quality covariates (cov7). Moreover, each of 18 covariates was added to cov4 one at a time, and the change in the Module post-IP% coefficient was recorded (Supplementary Table S12).

Two out-of-range values were kept as recorded: an MMSE of 33 (donor H20.33.002; cognitively normal, CASI 93) and a brain pH of 4.5 (donor H21.33.034; RIN 2.9). Setting the MMSE to missing or 30 changed no MMSE result, and the pH donor had negligible influence (Cook's distance ≤0.020).

***Benchmark against competitors***

Five competitors were computed per donor from all cells of the arm's supertype, on the arm's GOI and its eight partners and without CMRF input. Cluster proportion (CP) was the supertype's percentage of the donor's Micro-PVM cells, and AUCell, UCell, and AddModuleScore were computed per cell and averaged per donor. The changepoint split (CS) was the donor percentage of cells with the GOI above the breakpoint of a segmented regression of the partner mean on the GOI.

In Figure 10C, a nested F-test compared outcome ~ covariates + AUCell + CP + Module post-IP% with the same model without Module post-IP%. Figure 10D repeated this in M4 with each of AUCell, UCell, and AddModuleScore; the FDRs and labels in Figure 10D are from the same model without CP (Supplementary Table S12).

***Boundary ablation***
In Figure 11, three simpler boundaries were built in the copula plane: a band around an OLS line of v on u, a band around a TPRS (fixed spline), and the same spline with a half-width estimated in 20 bins of u (local spline). Each boundary held about as many cells as the CMRF Module, hence the boundaries differed only in shape. Cells outside a boundary were its discarded cells, tested by the discarded-cell term of outcome ~ cov4 + module% + discarded%.

**Supplementary Methods A2. Settings shared by the Batteries**
A unit was one donor in one region in one arm, and it needed at least 10 labeled cells (Section 2.1). There were 125 units from 51 donors in M4, 450 from 83 donors in PVM_3, 201 from 66 donors in PVM_1, and 80 from 38 donors in HKG8, yielding 856 units from 83 donors in total. For within-donor tests (in donors with at least 3 units), each unit's value was centered on its donor's mean before $r_s$ was computed. For between-donor tests, each donor's mean over its units was used.

%MT was the mean share of UMIs from the 13 mitochondrial protein-coding (MT) genes over the arm's labeled cells, log1p-transformed. %MT does not use the housekeeping genes, so in HKG8 the %MT results describe the mitochondrial share of those cells and are not related to the HKG8 Module. Contamination was the DecontX estimate per nucleus.

A test needed ≥10 donors, and in Batteries 1, 2, and 4 a donor entered a within-donor comparison with ≥20 cells per group. Bootstraps used B=10,000 resamples. Clinical outcomes were interpreted by the OLS *p*-value. The BH families and the reading of every panel were fixed in pre-specified reading files (deposited on GitHub) before each run.

Two readings were changed after the results were observed: 1) For the offset share (Battery 2), FDR replaced the uncorrected 95% CI, and M4 then did not pass (FDR=0.124); 2) For the donor requirement (Battery 4), only the clinical association rule was used, as the M4 split-half rule never reached 80% detection at 0.75x the observed effect.

**Supplementary Methods S10. Battery 1 (Figure 12 and Supplementary Figure S15)**
Each model of Figure 9A was refitted with a contamination covariate or after removing doublets (Figure 12A and 12B). The covariate was the donor mean DecontX contamination of the arm's non-Module cells (≥5 cells), and doublets were Module post-IP nuclei with a SEA-AD doublet score ≥0.15, 0.20, or 0.30. Retention was the adjusted $\Delta R^2$ divided by the unadjusted $\Delta R^2$. Each model was then classed by whether it kept, lost, or gained significance after adjustment.

At the cell level, cells from both region sets were pooled within each donor. For Figure 12D, each donor's cells were split into DecontX tertiles, and the tertiles were then pooled across donors. Module post-IP and pre-IP cells were also compared on three shares of their non-mitochondrial UMIs: own endoplasmic reticulum (ER; *CALR, CANX, HSPA5, PDIA3, P4HB, SEC61A1, SEC61B, RPN1, DDOST*, and *SSR4*), ribosomal (*RPLP0, RPS18*, and *RPS6*), and *NEAT1*. They were also compared on an other-lineage index from five non-microglial marker panels, without markers that were post-IP DEGs (*SYT1* and *MEG3* in NeoC; Supplementary Figure S15).

The Gerrits et al. [6] dataset (36 samples from 18 donors) was used for technical checks only. Cells were called by `DropletUtils::emptyDrops`. The ambient index of a nucleus was the share of its UMIs that fell on the marker panels of other lineages. The ambient pool was the set of barcodes with 1-100 UMIs. Its mitochondrial fraction was compared with that of the called nuclei (Figure 12E). The $r_s$ of *MT-CO3* with each DFG was computed without adjustment and as a partial correlation adjusted for the ambient index and log depth (Figure 12F). It was also computed within the ambient-index tertiles of each sample (Figure 12G).

Finally, the 24,463 nuclei of the SEA-AD PVM_3 arm (median 5,650 UMIs) were thinned to lower depths by binomial sampling, with 20 repeats per depth. The target depths ranged from 8,000 down to 500 UMIs and included 1,504 UMIs, the median depth of the Gerrits data. Each depth was run with and without also thinning the MT genes to the Gerrits mitochondrial fraction of 0.12%. The median $r_s$ over the eight DFGs was then compared with the per-sample $r_s$ in the Gerrits data. *ACTB* and cHKG8 served as controls (Figure 12H).

**Supplementary Methods S11. Battery 2 (Figure 13 and Supplementary Figure S16)**
The donor share of variance (between-donor over total sum of squares) was computed for Module post-IP%, its residual after tissue quality was removed (TQR), %MT, unit mean contamination, and a matched non-mitochondrial gene share, and was compared with a null of 10,000 within-region shuffles of donor labels (Figure 13A). For split-half $r_s$, each donor's units were split at random into two halves, and the mean of each half was computed. The $r_s$ between the two half means was then computed over donors. This was repeated 200 times, and the median $r_s$ was reported (Figure 13B).

The matched non-mitochondrial share was a control for %MT. For each of the 13 MT genes, the 30 non-mitochondrial genes most similar to it in mean count and detection rate were listed. In each of 200 draws, one gene was picked at random from each list, yielding a set of 13 matched genes. The share of UMIs on these genes was then averaged over the 200 draws. The drawn genes are listed in `02_matched_sets.csv` in the GitHub repository.

Cohen's κ was computed only in regions with ≥20 shared cells (Figure 13C). The $r_s$ of *MT-CO3* with cDFG was computed within each donor that had ≥20 PVM_3 arm cells. It was computed without adjustment and as a partial correlation adjusted for log total UMIs (Figure 13D). A median within-donor $r_s$ of ≥0.75 meant that the association held within donors. A median of 0.40 to <0.75 meant that it held partly within donors. A median of <0.40 meant that it arose mostly between donors.

The offset share was the part of a nucleus's mitochondrial UMIs that stayed roughly constant as sequencing depth changed. Ambient RNA is expected to behave this way, as each droplet picks up a similar amount of it regardless of the nucleus's sequencing depth. Mitochondrial UMIs from the nucleus itself are expected to rise with depth instead. For each donor, mitochondrial UMIs were regressed on total UMIs by OLS, separately in Module post-IP and pre-IP cells. The intercept (a) is the fixed part, and the slope (b) times depth is the depth-dependent part. The offset share was median a/(median a + median b x reference depth), with the median total UMIs of the Module cells as the reference depth (Figure 13E). The post-IP and pre-IP offset shares were compared by paired donor bootstrap and interpreted as equivalent when the 90% interval of their difference was within ±10 pp with FDR≥0.05.

Finally, the $r_s$ of Module post-IP% with five library fields was computed within and between donors (Figure 13F and 13G; Supplementary Figure S16). The fields were the intronic and antisense mapping rates, the NeuN-positive fraction, the percentage of cDNA longer than 400 bp, and PCR cycles. Within donor, a library field was interpreted as ordering the Module only when four conditions all held:

1) The $r_s$ of Module post-IP% with the field reached FDR<0.05.
2) The Module $r_s$ differed from the $r_s$ of sequencing depth with the same field. The 95% CI of the difference excluded 0.
3) The Module $r_s$ differed from the $r_s$ of the HKG8 control with the same field. The 95% CI of the difference excluded 0.
4) The Module $r_s$ was not clearly weaker than the %MT $r_s$ with the same field. The upper bound of |Module $r_s$| minus |%MT $r_s$| was ≥0.

**Supplementary Methods S12. Battery 3 (Figure 14 and Supplementary Figure S17)**
Each unit was one donor in one brain region. The units were matched by donor and region to two SEA-AD resources. The first was the quantitative neuropathology of the gray matter, and the second was the CPS. The 28 stain measures fell into seven families: amyloid 6E10, tau AT8, TDP43, synuclein, astrocyte GFAP, microglia IBA1, and neuron NeuN. The amyloid family included two

measures of 6E10 and IBA1 colocalization. The CPS had three local and three global scores, each for overall, Aβ, and pTau. Local scores varied by region, while each donor had only one value for each global score. In total, 822 of the 856 units had a match. For the histology analyses, the two region sets were pooled within each donor.

The within-donor $r_s$ with IBA1 was computed only for donors with ≥3 units. For two IBA1 measures, $IBA1^+$ cells per area and $IBA1^+$ area (%), no expected direction was pre-specified, hence they were tested two-sided (Figure 14A). The CPS were also correlated with the log cell count and the contamination level of each unit (Figure 14B).

Module post-IP% and %MT were then entered together in one model for each comparison. For histology, the measure and both predictors were converted to ranks and standardized. The ranked measure was then regressed on ranked Module post-IP% and ranked %MT, with no covariates (Figure 14C). Each histology model was fitted at two levels, within donors and between donors. For the within-donor fit, each variable was centered on its donor mean before ranking.

For the clinical outcomes, the model of Figure 9 was fitted separately for each region set, using the donors of Figure 12A (Figure 14D). A comparison was considered unreadable when the VIF between the two predictors was ≥5, which corresponds to $|r|$ ≥0.894 between them. A comparison was considered untestable when its model could not be fitted.

**Supplementary Methods S13. Battery 4 (Figure 15 and Supplementary Figures S18 and S19)**
Fifteen panels of nuclear-encoded mitochondrial and metabolic genes were scored in each cell. A reference panel of 300 matched genes was scored the same way. Each panel used only the genes detected in ≥10% of the arm's cells, and a panel needed at least three such genes to be scored. A cell's panel score was the mean log1p CP10K of these genes. For donors with ≥20 labeled cells, the donor score was the median of the scores of those cells. No VIF reached 5 in the Figure 9 models with Module post-IP% or %MT (Figure 15A and Supplementary Figure S18).

For each arm and region set, the area under the curve (AUC) measured how well per-cell %MT separated Module cells from non-Module cells. It was computed over the pooled labeled cells (36,967 cells). The best %MT threshold was chosen by Youden's J, and Cohen's κ was computed at that threshold. An AUC below 0.60 meant that %MT did not recover the Module. An AUC of 0.60 to <0.80 meant that it recovered the Module partly. An AUC of ≥0.80 meant that the Module was close to a simple %MT threshold (Figure 15B and Supplementary Figure S19). The AUC was also computed within each donor with ≥20 Module and ≥20 non-Module cells. The donor median AUC was tested against 0.5 by donor bootstrap (Figure 15C).

The number of donors needed (Figure 15D) was estimated by subsampling. For each donor count from 10 to 83, 300 random sets of donors were drawn without replacement. The donor share of Module post-IP% was also scaled down to 1.0x, 0.75x, or 0.5x of its observed value, to test smaller effects. In each draw, MMSE was regressed on cov4 and the donor mean Module post-IP% by OLS. The donor requirement was the smallest donor count at which this model reached $p$<0.05 in ≥80% of draws. Two other criteria were also scored at each donor count. The first was a donor share above the 95th percentile of its shuffle null. The second was a split-half $r_s$ of ≥0.50.

**Supplementary Methods S14. Battery 5 (Figure 16)**
Battery 5 was added after Batteries 1 to 4 were complete, and all 84 donors remain primary. The nine exposed panels (Figures 12A, 13A, 13B, 13F, 14A, 14B, 14C, 14D, and 15A) were recomputed without the SA donors of each frame, the donor set from which a statistic is computed (8 to 11 SA donors per frame). The Figure 12A and 14D models used the donors of Battery 1.

For each comparison, the change was the estimate without the SA donors minus the estimate with all donors. The change was multiplied by the sign of the all-donor estimate, thus a negative change meant a weaker effect. Each change was compared with the changes from 1,000 random removals of as many non-SA High-ADNC donors as SA donors in the frame, from the same donor set. Each significant comparison was then placed in one of four classes:

1) Held. It stayed significant with the same sign.
2) Reversed. It stayed significant with the opposite sign.
3) Lost beyond random removal. It lost significance, and its change was below the 5th percentile of the random changes.
4) Lost within random removal. It lost significance, but its change was not below that percentile.

A panel held when none of its comparisons was reversed or lost beyond random removal (Supplementary Table S17).

Supplementary Table S17 also provides the Figure 14C within-donor, 14D, and 15A verdicts with and without the 11 SA donors, and leave-one-donor-out estimates for Figures 13A, 13B, and 13F.

**Supplementary Methods S15. M4 against PVM_3 and the summary of all tests (Figure 17; Supplementary Figures S20 and S21)**

The two analyses below used the same donor sets as Figure 9, the same cell labels as the Batteries, and the same %MT as Figure 14D. The M4 share models used a donor bootstrap with B=10,000 (Supplementary Figure S20).

In the random-subset analysis (Supplementary Figure S21), random sets of PVM_3 arm cells were drawn without replacement 10,000 times. This comparison was valid because, in the cells that both arms selected, the Module post-IP labels of the PVM_3 arm agreed with those of the M4 arm (Cohen's $\kappa$=0.884 in NeoC and 0.939 in NonNeoC). The M4 value was compared with the values from the 10,000 draws by a two-sided empirical *p*-value.

Figure 17B tallies tests 1 to 13 except tests 7 and 11, for which PVM_3 and PVM_1 had no readable model (VIF≥5; Supplementary Table S19). Each test is shown as the models held in both region sets over those read in both, and as the mean percentage of models held, boxed at ≥75%. The pathology tests used Thal phase and Braak stage, the amyloid and tau measures of the NIA-AA ABC score [7], chosen beforehand. For pathology, test 2 added MMSE and CASI to the model, and test 4 was read by the unadjusted *p*-value (Supplementary Table S12). Test 8 used the module-only models at cov4 (Supplementary Table S12).

Tests 9 to 11 were interpreted without the Battery 1 entry condition. Comparisons that were not significant on the Battery 1 donors were therefore still assessed, and they showed as not holding. Two examples are the HKG8 NeoC CASI score (FDR=0.102) and Thal phase (FDR=0.065). In Figure 17B, k and n are summed over the region sets. Supplementary Table S19 provides each region set separately. A hyphen marks a test that could not be assessed for one of these reasons:

1) The VIF was ≥5.
2) There were too few donors.
3) The result was inconclusive.
4) The retested model had no all-donor association.
5) The test was test 7 for a pathology outcome.

**Supplementary Methods S16. External validation in two independent datasets (Figures 18 and 19; Supplementary Tables S20 and S21)**

External dataset 1 comprised the prefrontal cortex (BA9) snRNA-seq data of Pineda et al. [8] (batch 210526): 63 donors (23 ALS, 24 FTLD, and 16 pathologically normal [PN]) with one library each. Raw reads (SRA PRJNA1073234) were counted with Cell Ranger 10.1.0 (GRCh38-2020-A, intronic reads included), cells were called with emptyDrops (FDR≤0.01), and nuclei with <500 genes and scDblFinder doublets were removed. Only nuclei kept by the authors' quality control were analyzed (≥700 genes, ≤20% MT, and removal of low-quality clusters). The authors' quality control was applied before cell type mapping, because MapMyCells assigned different labels to the kept nuclei when the removed nuclei were mapped in the same file. Results without the authors' quality control are shown in Supplementary Table S20. Cell type mapping (below) is not deterministic, and mapping the same file twice changed the supertype of 0.8% of nuclei, so the mapping results are deposited on Hugging Face. Age at death, sex, and PMI were from the authors' donor table (Synapse syn51105515).

External dataset 2 comprised the insular cortex snRNA-seq data of Rexach et al. [9]: 37 donors (AD, bvFTD, PSP, or controls) with one 10x Genomics 3′ v3 library each. Raw counts were from CELLxGENE (dataset aa87f914-07e0-48aa-b915-8de906c95baf), and age at death and PMI from the authors' Table S1. The authors' quality control was kept (nuclei with <200 genes or >8% MT removed), and nuclei with <500 genes were also removed.

Cell types were mapped to the SEA-AD taxonomy (CCN20230505) by MapMyCells Deep Generative Mapping without a confidence cutoff. The PVM_3 arm comprised Micro-PVM_3 nuclei with *MT-CO3*>0, and the HKG8 control its nuclei with *ACTB*>0 (main text Section 2.1). The R package cepip (0.1.0) was run at the SEA-AD settings on the composite of the eight SEA-AD DFGs. The DFGs replicated if at least five of the eight passed tier 1 with *MT-CO3* ($r_s$≥0.6 and $\tau_b$≥0.5), or otherwise tier 2 ($r_s$≥0.5 and $\tau_b$≥0.4) [5]. Module post-IP% was compared between diagnoses only if the arm reached an IPRS of at least 0.60.

In external dataset 1, Module post-IP% of FTLD and of ALS was compared with PN by OLS on cov2 (age at death and sex) and on cov3 (PMI missing for 7 of 16 PN donors). These comparisons used the 48 donors with at least 5 Micro-PVM_3 nuclei and a recorded age at death. In external dataset 2, Module post-IP% of AD, bvFTD, and PSP, combined and each alone, were compared with controls in the 33 donors with at least 5 Micro-PVM_3 nuclei, by OLS on cov3 (age at death, sex, and PMI). FDR was computed over the four comparisons of each dataset. Leave-one-donor-out refits, a donor bootstrap stratified by diagnosis (10,000 resamples), and the power to detect the SEA-AD NeoC cognitive-status β were added post hoc.

For external dataset 2, container reruns by the author and by a second person on Windows 11 agreed with all 113 original output tables. For external dataset 1, the author's rerun from the Cell Ranger count matrices yielded identical cell calls in all 63 libraries, and 151 of 152 output tables agreed; the other table differed only in scDblFinder scores (by ≤$3.7x10^{-4}$), with no change in doublet calls. A second person reran the steps after cell type mapping, and all 93 output tables agreed.

**Supplementary Methods A3. Code availability**

The analysis ran as a series of numbered scripts, in the following order:

1) Python Parts 1 to 8: These covered the data checks and cell selection, and ended with the export of the four arms' data as CP10K count matrices.
2) R Parts 9 to 10d: Fitted the GAMs, computed the SKCS scores, detected the IP, and derived the copula coordinates.
3) R Parts 10e to 11: These built the CMRF module and tested it, from its validation and clinical correlation to the boundary ablation.
4) Part12_01 to Part12_06: Extracted the inputs for the Batteries and called the Gerrits cells.
5) Part12_Fig15_Analysis to Part12_Fig19_Analysis: Conducted Batteries 1-5, one script per Battery.
6) R Part13 and Part13b: Performed the M4 share and random-subset analyses.

The container recipe was based on `rocker/r-ver:4.6.1` (Ubuntu 24.04), and `renv::restore()` installs the same R package versions without the container. The container holds no data, and the input folder is mounted at run time. Every script that drew random numbers used fixed seeds.

The container image was built from the recipe and also deposited on Zenodo. The one command that reran the whole pipeline from the SEA-AD h5ad to Figure 17 and Supplementary Tables S1 to S19 is as follows:

```
podman run --rm --shm-size=16g -v /path/to/SEA-AD:/data:Z seaad-cepip seaad-run all
```

Further modes rerun only the Python parts, the R parts (also from the Hugging Face files), or the supplementary tables, in Podman or Docker. `README_container.md` lists the inputs, commands, and run order. The two external validations (Supplementary Tables S20 and S21) ran in separate container images, which are also deposited on Zenodo, and their scripts are in the external_validation folder of the GitHub repository.

For the reproducibility check, Python Parts 1 to 8 were rerun from the SEA-AD h5ad and R Parts 9 to 13 from the Part 8 export, and `compare_rerun.py` compared every table with the original results. Of the 1,203 R tables, 1,150 were identical or differed only in sign, order, text, size, or by ≤0.1%, and the other 53 held the secondary statistics of Section 2.10. Of the 9,262 Python tables, 9,253 were identical or differed only in format. The results for each table are in `compare_all_detail.csv` deposited on GitHub.

The independent test (Section 2.10) was run on 21 October 2026 with Docker Desktop, cepip 0.1.0, and commit d4a82aa. Of the 1,073 tables compared, 1,020 agreed within 0.1% (i.e., every value differed from the original by at most 0.1% of its size), and the other 53 held the same secondary statistics as above, none of which changed a significance call. The clinical associations were identical (outputs in the GitHub repository, folder `verification/windows_test/`).

Some labels inside Supplementary Tables S13 to S17 used the figure numbers of the original analysis run. In these tables, Figures 12 to 19 correspond to Figures 9 to 16 of this paper, and Supplementary Figures 11 to 15 correspond to Supplementary Figures S15 to S19. The tables also used “soup index” for the ambient index and “second only” for the Module only and %MT only verdicts of Figure 15A. The original workbooks are available from the corresponding author on request.
